# Identifying miRNA Signatures Associated with Pancreatic Islet Dysfunction in a FOXA2-Deficient iPSC Model

**DOI:** 10.1101/2024.06.15.599142

**Authors:** Ahmed K. Elsayed, Noura Aldous, Nehad M. Alajez, Essam M. Abdelalim

## Abstract

The pathogenesis of diabetes involves complex changes in the expression profiles of mRNA and non-coding RNAs within pancreatic islet cells. Recent progress in induced pluripotent stem cell (iPSC) technology have allowed the modeling of diabetes-associated genes. Our recent study using FOXA2-deficient human iPSC models has highlighted an essential role for FOXA2 in the development of human pancreas. Here, we aimed to provide further insights on the role of microRNAs (miRNAs) by studying the miRNA-mRNA regulatory networks in iPSC-derived islets lacking the FOXA2 gene. Consistent with our previous findings, the absence of FOXA2 significantly downregulated the expression of islet hormones, INS, and GCG, alongside other key developmental genes in pancreatic islets. Concordantly, RNA-Seq analysis showed significant downregulation of genes related to pancreatic development and upregulation of genes associated with nervous system development and lipid metabolic pathways. Furthermore, the absence of FOXA2 in iPSC-derived pancreatic islets resulted in significant alterations in miRNA expression, with 61 miRNAs upregulated and 99 downregulated. The upregulated miRNAs targeted crucial genes involved in diabetes and pancreatic islet cell development. In contrary, the absence of FOXA2 in islets showed a network of downregulated miRNAs targeting genes related to nervous system development and lipid metabolism. These findings highlight the impact of FOXA2 absence on pancreatic islet development and suggesting intricate miRNA-mRNA regulatory networks affecting pancreatic islet cell development.

## Introduction

Diabetes mellitus (DM) is caused by the dysfunction or loss of insulin-producing pancreatic β-cells [1]. Understanding the molecular mechanisms governing human pancreas development holds potential for identifying novel strategies for DM treatment. Diverse signaling pathways and transcription factors (TFs) play essential roles in orchestrating the differentiation and maturation of endocrine pancreas [2]. While animal studies have long been instrumental in biomedical research, discrepancies in physiology and metabolism between animal models and humans hinder direct translation of findings to human diabetes pathophysiology. Differences in gene expression profiles of endocrine islet cells have been documented [3, 4]. Human pluripotent stem cells (hPSCs) present a renewable resource for generating functional insulin-producing β-cells and serve as a promising platform for disease modeling and drug discovery [5–7]. Recent advancements in differentiating functional β-cells from hPSCs, combined with genome-editing technologies, offer avenues to investigate diabetes-associated genes and discover novel therapeutic approaches [5–7].

MicroRNAs (miRNAs), a group of conserved non-coding RNAs consisting of 19-25 nucleotides, exert significant influence on diverse biological processes and are implicated in various diseases. Primarily, they function by suppressing the translation or facilitating the degradation of their target mRNAs [8]. While several miRNAs have been suggested to control multiple TFs of pancreatic β-cells during their development, many of these miRNA-mRNA interactions predicted through computational methods have yet to be confirmed through experimental validation. Numerous miRNAs have been recognized for their pivotal involvement in diabetes [9, 10], the development of all endocrine cell types [11, 12], and integral contribution to the proper functioning of mature β-cells [13, 14]. Certain miRNAs have been associated with processes relevant to type 2 diabetes (T2D), including apoptosis, response to cytokines, and insulin secretion [15]. Several studies support the idea that miRNAs play critical roles in maintaining β-cell identity [11, 13–16] and that their expression is changed in diabetic conditions [15, 17]. Also, various dietary and environmental factors have been shown to influence the miRNA profile within pancreatic islets [15].

FOXA2, a crucial TF, expression is initiated at the earliest stage of pancreatic development, and its presence continues throughout subsequent developmental phases. Recent studies conducted in our laboratory have unveiled that the absence of FOXA2 during the differentiation of induced pluripotent stem cells (iPSCs) into pancreatic and hepatic cells results in pancreatic islets’ and hepatic cells’ developmental defects [18, 19]. Furthermore, our findings have brought to light changes in mRNA profiles associated with FOXA2 deficiency that are accompanied by significant alterations in the expression of long non-coding RNAs (lncRNAs) at the pancreatic progenitor (PP) and pancreatic islet stages [20]. Recently, we have also unraveled the connection between dysregulated miRNAs and differentially expressed genes (DEGs) in PPs lacking FOXA2 [21]. However, a significant gap in knowledge exists concerning the consequences of FOXA2 deficiency on the expression profiles of miRNAs and their specific target genes within pancreatic islets. Individual miRNAs can suppress hundreds of mRNA targets, and several miRNAs frequently align to influence a specific pathway, facilitating a shared developmental result [22]. Therefore, in this study, to fully understand how miRNAs and their respective targets lead to the suppression of pancreatic islet development in the absence of FOXA2, we employed genome-wide sequencing to identify DEGs and differentially expressed miRNAs (DEmiRs) dysregulated in pancreatic islets lacking FOXA2. Through the integration analysis of DEmiRs and DEGs, we conducted a network analysis to uncover candidate miRNA-regulated genes. Our results showed that the absence of FOXA2 is linked to alterations in miRNA expression, particularly those miRNAs that target pivotal pancreatic genes, during pancreatic islet development. These shifts in miRNA expression, observed in iPSC-derived islets lacking FOXA2, may signify a disruption in the process of pancreatic differentiation.

## Materials and Methods

### Culture and differentiation of iPSCs into pancreatic islets

*FOXA2* knocked out iPSCs (*FOXA2^−/−^* iPSCs) as well as their wildtype (WT) control iPSCs (WT-iPSCs) previously established and fully characterized in our laboratory were used in this study [23]. All iPSCs lines were maintained in Stemflex media (ThermoFisher Scientific) on plates coated with hESC qualified Matrigel (Corning #354277). Differentiation of iPSCs into pancreatic islets was carried out following our previously published protocol (Supplementary Table 1) [18, 24, 25].

### Immunofluorescence

The cultured cells were rinsed twice with phosphate-buffered saline (PBS) and subsequently exposed to 4% paraformaldehyde (PFA) for 15-20 minutes at room temperature. The cells were subjected to three consecutive 10-minutes washes with tris-buffered saline containing 0.5% Tween 20 (TBST). Subsequently, the cells were permeabilized for 15 minutes at room temperature using PBS containing 0.5% Triton X-100 (PBST) in two rounds, followed by an overnight blocking step at 4°C in PBST with 6% Bovine Serum Albumin (BSA). The primary antibodies were diluted in PBST with 3% BSA and the cells were subjected to an overnight incubation with these antibodies at 4°C followed by three rinses with TBST. Afterward, the cells underwent a 1-hour incubation period with PBS-diluted secondary antibodies (1:500) at room temperature. To visualize the nuclei, the stained cells were treated with Hoechst 33258 (1:5000 in PBS) for 5 minutes, followed by three PBS washes prior to imaging. Images were acquired using Olympus inverted fluorescence microscope. The used primary antibodies were anti-insulin (GN-ID4, DSHB; 1:2000) and anti-glucagon (sc-514592, Santacruz; 1:1000). The secondary antibodies were Alexa Fluor 488 donkey anti-Rat IgG (H+L) (Thermo Fisher, A21208; 1:500) and Alexa Fluor 555 goat anti-Mouse IgG (H+L) (Thermo Fisher, A-21422; 1:500).

### Western blotting

Twenty micrograms of total protein, extracted from a single well of a 6-well plate using a mixture of RIPA lysis buffer and protease inhibitor then quantified using Pierce BCA kit (ThermoFisher Scientific), were separated via SDS-PAGE and subsequently transferred onto PVDF membranes. Following blocking of the membranes with 10% skimmed milk in TBST, the membranes underwent an overnight incubation at 4°C with primary antibodies diluted in 5% skimmed milk in TBST, rabbit anti-FOXA2 (1:4000, 8186S, Cell Signaling), and mouse GAPDH (1:10,000, sc-32233, Santa Cruz). After thorough TBST washing, horseradish peroxidase-conjugated secondary antibodies (Jackson Immunoresearch) were applied (1:10,000) for 1 hour at room temperature, followed by additional rounds of TBST washing. Finally, the membranes were developed using the SuperSignal West Pico Chemiluminescent substrate (Pierce) and imaged using the iBright™ CL 1000 Imaging System (Invitrogen).

### RNA extraction and expression analysis via RT-qPCR

Total RNA was extracted using the Direct-zol™ RNA Miniprep kit (Zymo Research, USA) from a single well of 6-well plate. For gene expression analysis, cDNA was synthesized from 1 µg of RNA using the SuperScript™ IV First-Strand Synthesis System following the manufacturer’s protocol (ThermoFisher Scientific, USA). RT-qPCR using the GoTaq qPCR SYBR Green Master Mix (Promega, USA) run in triplicates was done for mRNA expression analysis. To ensure accurate quantification, average Ct values were normalized to the corresponding values from WT samples for each tested gene, with GAPDH serving as the endogenous control (used primers are listed in Supplementary Table 2).

For reverse transcription of miRNA, 10 ng of total RNA was used for first strand cDNA synthesis using miRCURY LNA RT Kit (QIAGEN, Cat. No. 339340). Quantification of the relative miRNAs’ expression level was detected by specific miRCURY LNA miRNA PCR Assays and highly sensitive miRCURY LNA SYBR^®^ Green PCR Kit while adjusting the reverse transcriptase product dilution to 1:30 (QIAGEN, Cat. No. 339345). The calculation of relative miRNA expression was based on the -ΔΔCT method, where SNORD48 was employed as the endogenous control for miRNA expression normalization. Used miRCURY LNA miRNA PCR Assays are listed in Supplementary Table 2.

### Assessment of the differential gene expression

The quality and quantity of extracted RNA were assessed using on-chip electrophoresis with the Agilent RNA 6000 Nano Kit and Agilent 2100 Bioanalyzer, following the manufacturer’s instructions. Samples with an RNA Integrity Number (RIN) > 9.0 were selected for library preparation. In accordance with the manufacturer’s instructions, the NEBNext Poly(A) mRNA Magnetic Isolation Kit (NEB, E7490) was employed to capture mRNA, utilizing 1µg of total RNA. The RNA-sequencing (RNA-seq) libraries were generated using the NEBNext Ultra Directional RNA Library Prep Kit (NEB, E7420L), The yield of cDNA libraries was measured using the Qubit dsDNA HS Assay Kit, and the size distribution of the cDNA libraries was determined using the Agilent 2100 Bioanalyzer DNA1000 chip. Subsequently, libraries were pooled and subjected to sequencing on the Illumina Hiseq 4000 platform, resulting in an average read count of >8,400,000 per sample. To process the raw data, the Illumina BCL2Fastq Conversion Software v2.20 was utilized, with concurrent quality control procedures. Subsequently, using the built-in module and default settings within CLC Genomics Workbench v21.0.5, pair-end FASTQ files were aligned to the GRCh38 reference genome. For differential expression analysis, the normalized expression data represented as TPM (transcripts per million) mapped reads were imported into AltAnalyze v.2.1.3 as described before [26]. Differentially expressed genes (DEGs) were identified based on criteria where the log2 fold change (FC) exceeded 1 or was less than -1, accompanied by a *P-value* < 0.05. Functional enrichment analyses encompassing Gene Ontology (GO) and Kyoto Encyclopedia of Genes and Genomes (KEGG) pathways were conducted using the Database for Annotation, Visualization, and Integrated Discovery (DAVID) [27].

### Differential miRNA expression and potential target analysis in iPSC-derived pancreatic islets

miRNA library was prepared from approximately 100ng of the extracted total RNA, following the manufacturer’s guidelines provided with the library kit (E7560S, New England BioLabs Inc., USA). Monarch PCR purification kit (Biolabs, New England) used to purify the resulting amplified cDNA constructs. For miRNA analysis, FASTQ files were aligned to the miRBase v22 database (mature miRNAs), and miRNA expression (total counts) was estimated using the built-in small RNA analysis workflow with default settings in CLC Genomics Workbench 20.2, as previously described [28]. Over 70% of the annotated reads mapped to the miRBase v22 database, with more than 75% of those reads exhibiting a perfect match. Approximately 57% of miRBase-annotated miRNAs were detected in our samples. The CLC analysis pipeline did not provide information on other small RNAs, as this study focused specifically on miRNAs. Our analysis exclusively focused on reporting mature miRNAs (5p and 3p) according to miRBase v22 database annotations. To ensure accuracy, miRNA count reads were normalized using the TMM (trimmed mean of M values) method, and the resulting log2 CPM (Counts per Million) values were then subjected to subsequent differential analysis. Differentially expressed miRNAs in *FOXA2^−/−^* iPSCs compared to WT-iPSCs derived pancreatic islets were identified based on a log2 FC >1 with a *P-value* <0.05 threshold. Pathway analysis and the miRNA target filter identification were employed to unveil potential miRNA– mRNA networks, facilitated by Ingenuity Pathway Analysis (IPA) software (QIAGEN, Germany).

### Statistical analysis

Most experiments incorporated at least four biological replicates; in cases where this was not feasible, technical replicates were employed for subsequent statistical analysis. Prism 8 software was used to perform the statistical analysis, utilizing an unpaired two-tailed student’s t-test. The results are expressed as the mean ± standard deviation (SD).

## Results

### Profiling alterations in gene expression patterns in iPSC-derived pancreatic islets lacking *FOXA2*

To explore the impact of FOXA2 depletion on mRNA and miRNA expression profiles within pancreatic islets, we utilized two *FOXA2^−/−^* iPSC lines generated through CRISPR/Cas9 technology, along with their WT-iPSC control counterparts, as previously documented [18]. Using a stepwise protocol, the cell lines were differentiated into mature pancreatic islets (Fig. 1A and Supplementary Table 1). Our results demonstrated the complete absence of FOXA2 at protein level using western blotting (Fig. 1B) as well as clear demolishment in the number of INS- and GCG-expressing cells in *FOXA2^−/−^* islets (Fig. 1C). To identify the DEGs in pancreatic islets derived from both WT and *FOXA2^−/−^* iPSCs, we conducted RNA-Seq analysis. Our transcriptome analysis unveiled 719 DEGs that were significantly upregulated (Log2 FC > 1, *p* < 0.05), alongside 999 DEGs that were significantly downregulated (Log2 FC < −1, *p* < 0.05), when comparing *FOXA2^−/−^*islets to their WT-islet counterparts (Fig. 1D, E and Supplementary Tables 3 and 4). To visually represent the distinct transcriptomic profiles between WT-islets and *FOXA2^−/−^* islets, we performed PCA, which effectively clustered the data into two separate groups, underscoring the pronounced differences (Supplementary Fig. 1A-B). Further exploration of the downregulated DEGs through GO and KEGG pathways enrichment analyses revealed strong associations with pancreatic islet genes that are critical in processes such as diabetes mellitus, insulin secretion and processing, and glucose homeostasis (Fig. 1F, left panel and Supplementary Fig. 1C). The most important downregulated DEGs that play crucial roles in pancreatic islets’ development and functionality were enlisted in Fig. 2A. Conversely, when examining the upregulated DEGs, we observed enriched pathways primarily linked to nervous system development, lipid metabolism, bile acid and cholesterol metabolism, WNT signaling pathway, and various other metabolic signaling pathways (Fig. 1F, right panel). Absence of FOXA2 was correlated with the upregulation of many genes, primarily associated with processes related to fat and cholesterol metabolism, as well as lipid and neuronal development, among other metabolic pathways (Fig. 1F, Fig. 2A, and Fig. 3A). To validate the RNA-Seq findings, we conducted RT-qPCR analysis on a selection of crucial key pancreatic DEGs (Fig. 2B). Our RT-qPCR results provided robust confirmation of the RNA-Seq data, reaffirming the significant reduction of mRNA expression in pancreatic islet genes, including *INS, GCG, PPY, IAPP, ARX, NKX6.1, PDX1, GCGR, UCN3, NEUROG3, NEUROD1, NKX2.2, INSM1, PAX4, PAX6, RFX6, HES1, HES6,* and *PTF1A*, within *FOXA2^−/−^* islets (Fig. 2B). In addition, we validated some of the significantly upregulated DEGs at mRNA levels including *APOA2, APOC3, ABCC6, AFP,* and *OLIG3*. Also, some genes known to be expressed in pancreatic islets such as *LIN28A, NPY,* and delta cell-specific markers, *HHEX* and *SST* were significantly upregulated (Fig. 2C and Supplementary Table 3).

**Fig. 1.**
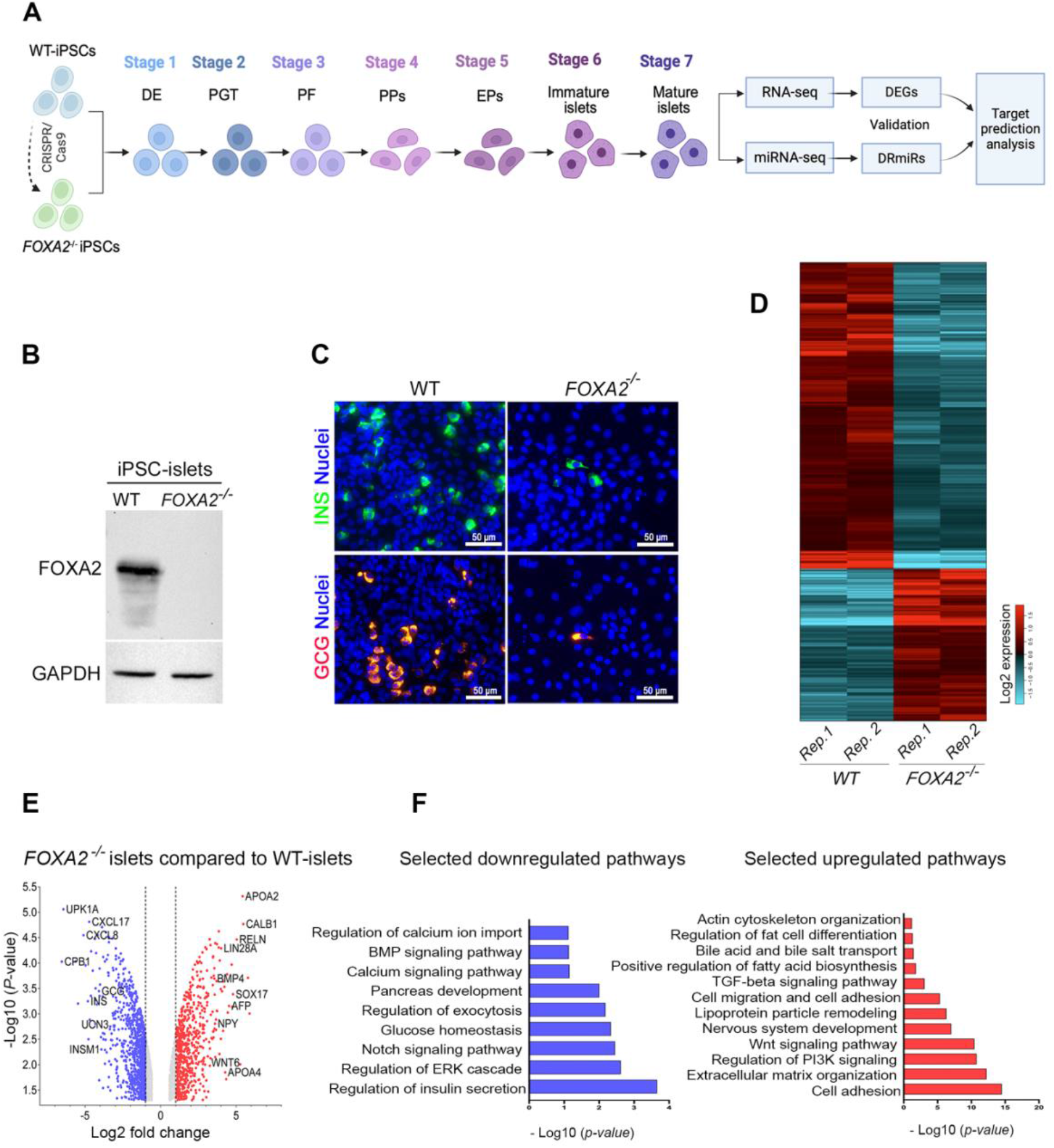
Effect of *FOXA2* loss on pancreatic islet differentiation. **(A)** Schematic representation of stepwise iPSC differentiation protocol into pancreatic islets. **(B)** Western blot analysis confirming the absence of FOXA2 expression in pancreatic islets derived from *FOXA2^−/−^* iPSCs (full-length blots/gels are presented in Supplementary Figure 1). **(C)** Immunofluorescence showing significant reduction in the expression of INS and GCG in *FOXA2^−/−^* islets. **(D)** General heatmap of differentially expressed genes (DEGs) in pancreatic islets derived from WT and *FOXA2*^−/−^ iPSCs. **(E)** Volcano plots representing the downregulated DEGs as blue dots while the red dots indicate upregulated DEGS in *FOXA2^−/−^* islets. **(F)** Gene ontology (GO) enrichment analysis terms for downregulated (blue) and upregulated (red) DEGs.

**Fig. 2.**
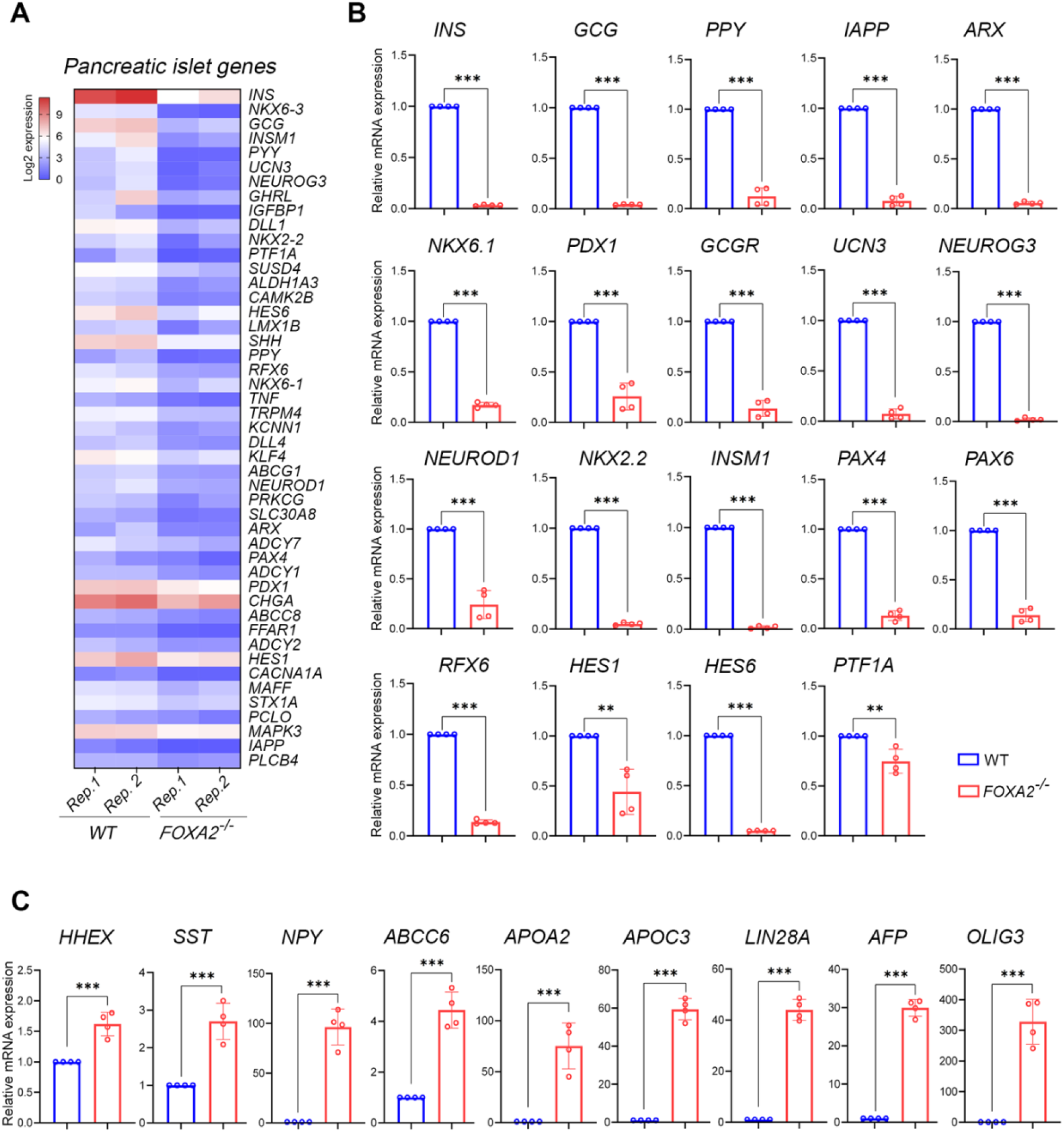
FOXA2 loss changes transcriptomic profile in pancreatic islets lacking FOXA2. **(A)** Heatmap for selected pancreatic islet-associated downregulated DEGs in *FOXA2*^−/−^ islets (n=2). RNA-Seq results were validated using RT-qPCR for selected downregulated DEGs (n=4) **(B)** and upregulated DEGs (n=4) **(C)**. Data are represented as mean ± SD.

**Fig. 3.**
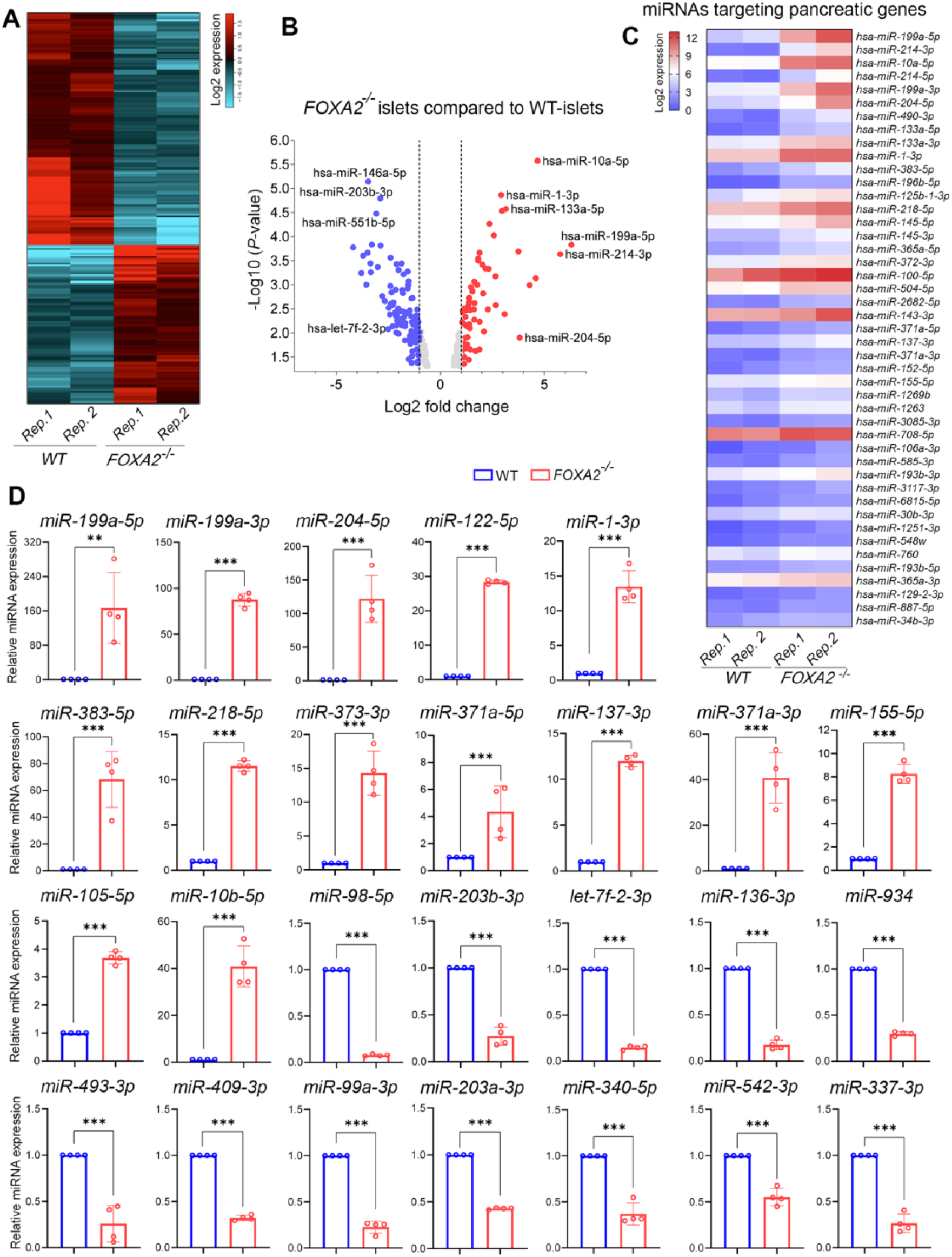
Differential miRNA expression profiling of *FOXA2^−/−^* islets and WT-islets. **(A)** General heatmap of dysregulated DEmiRs in *FOXA2^−/−^* islets compared to WT-islets (n=2). **(B)** Volcano plot of downregulated DEmiRs as blue dots and upregulated DEmiRs as red dots. **(C)** Heatmap of the selected upregulated DEmiRs targeting key pancreatic islet genes (n=2). **(D)** RT-qPCR validation of selected downregulated and upregulated DEmiRs (n=4). Data are represented as mean ± SD.

### Altered miRNA expression profile in iPSC-derived pancreatic islets lacking *FOXA2*

To unveil the alterations in the miRNA expression profile, we conducted miRNA-Seq analysis using the same RNA samples collected from iPSC-derived pancreatic islets of *FOXA2^−/−^* and WT. Our miRNA-Seq investigation identified 61 significantly upregulated miRNAs (Log2 FC > 1, *p* < 0.05) and 99 significantly downregulated miRNAs (Log2 FC < −1, *p* < 0.05), collectively referred to as DEmiRs, in *FOXA2^−/−^* pancreatic islets compared to WT counterparts (Fig. 3A, B). The top upregulated and downregulated DEmiRs are detailed in Supplementary Tables 5 and 6, respectively. Given the substantial suppression of β-cell development observed in the absence of FOXA2 [18], our study concentrated on the upregulated DEmiRs. Many of these DEmiRs are implicated in regulating the expression of key genes crucial for β-cell development [15, 29–31], and glucose regulation [15, 32] (Fig. 3C).

To validate our findings, we performed RT-qPCR analysis on selected DEmiRs. Our results demonstrated significant upregulation of miR-199a-5p, miR-199a-3p, miR-204-5p, miR-122-5p, miR-1-3p, miR-383-5p, miR-218-5p, miR-373-3p, miR-371a-5p, miR-137-3p, miR-371a-3p, miR-155-5p, miR-10b-5p, and miR-105-5p, along with significant downregulation of miR-98-5p, miR-203b-3p, let-7f-2-3p, miR-136-3p, miR-934, miR-493-3p, miR-409-3p, miR-99a-3p, miR-203a-3p, miR-340-5p, miR-542-3p, and miR-337-3p (Fig. 3D).

### Upregulated DEmiRs potentially target downregulated pancreatic genes in *FOXA2*^−/−^ *islets*

To gain insights into the roles of miRNAs during pancreatic islet cell development in the lack of FOXA2, we used the miRNA target filter in Ingenuity Pathway Analysis (IPA) Software to identify the potential target genes while focusing on the upregulated DEmiRs crosslinked with the downregulated DEGs from our datasets. The target prediction analysis revealed 55 significantly upregulated DEmiRs (Log2 FC > 1, *p* < 0.05) targeting 756 significantly downregulated DEGs (Log2 FC < −1, *p* < 0.05) in *FOXA2*^−/−^ islets. Among the 756 identified targeted DEGs, 32 were selectively identified as important pancreatic islet associated DEGs, which were targeted by 45 upregulated DEmiRs (Fig. 4 and Table 1). Hereafter, we focused on these selected DEGs, which are known for their pivotal roles in pancreatic islet development and function. Our investigation, with a particular emphasis on genes and TFs related to pancreatic endocrine specification, unveiled an intricate network of miRNA-mediated regulatory interactions. Notably, *NEUROG3* emerged as a *bona fide* target for the hsa-miR-372-3p, hsa-miR-106a-3p, hsa-miR-30b-3p, and hsa-miR-887-5p, while *NEUROD1* was a predicted target by hsa-miR-133a-5p, hsa-miR-1-3p, hsa-miR-372-3p, hsa-miR-2682-5p, hsa-miR-371a-5p, hsa-miR-137-3p, hsa-miR-1269b, and hsa-miR-1263. Furthermore, the NOTCH negative regulator TF, *HES6*, was predicted to be targeted by hsa-miR-214-3p, hsa-miR-371a-3p, hsa-miR-3085-3p, hsa-miR-6815-5p, and hsa-miR-30b-3p. Similarly, *LMX1B* was targeted by hsa-miR-214-3p, hsa-miR-3085-3p, hsa-miR-6815-5p, and hsa-miR-760. NKX6 TFs, including *NKX6.3* and *NKX6.1*, were targeted by several miRNAs, including hsa-miR-214-5p, hsa-miR-133a-3p, hsa-miR-365a-5p, and hsa-miR-504-5p for *NKX6.3*, and hsa-miR-548w, hsa-miR-193b-5p, and hsa-miR-34b-3p for *NKX6.1*. Moreover, *PDX1* and *PAX4* were anticipated targets for hsa-miR-193b-3p. *MAFF* was identified as a predicted target for 8 different DEmiRs including hsa-miR-214-3p, hsa-miR-133a-3p, hsa-miR-1269b, hsa-miR-3085-3p, hsa-miR-708-5p, hsa-miR-30b-3p, hsa-miR-760, and hsa-miR-193b-5p. *INSM1* was a predicted target for hsa-miR-125b-1-3p, and hsa-miR-100-5p. Additionally, *ARX* is predicted to be targeted by hsa-miR-204-5p, hsa-miR-372-3p, and hsa-miR-371a-3p.

**Fig. 4.**
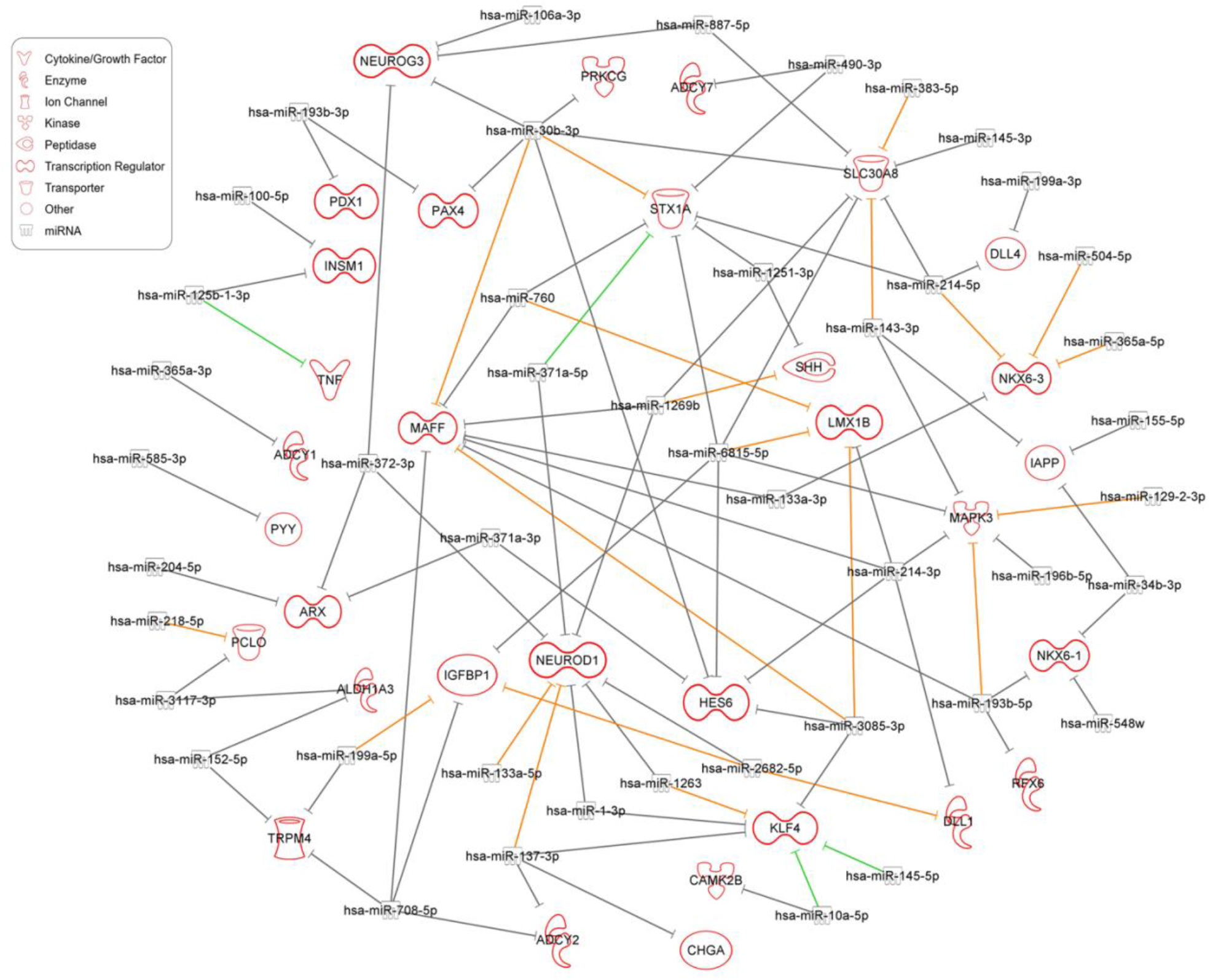
Target prediction analysis of upregulated DEmiRs targeting key downregulated pancreatic islet DEGs. The target prediction network between upregulated DEmiRs and downregulated DEGs was conducted using the Ingenuity Pathway Analysis (IPA) tool. Green lines indicate experimentally observed targets, orange lines represent a high prediction confidence level, while grey lines indicate a moderate prediction confidence level.

**Table 1.**
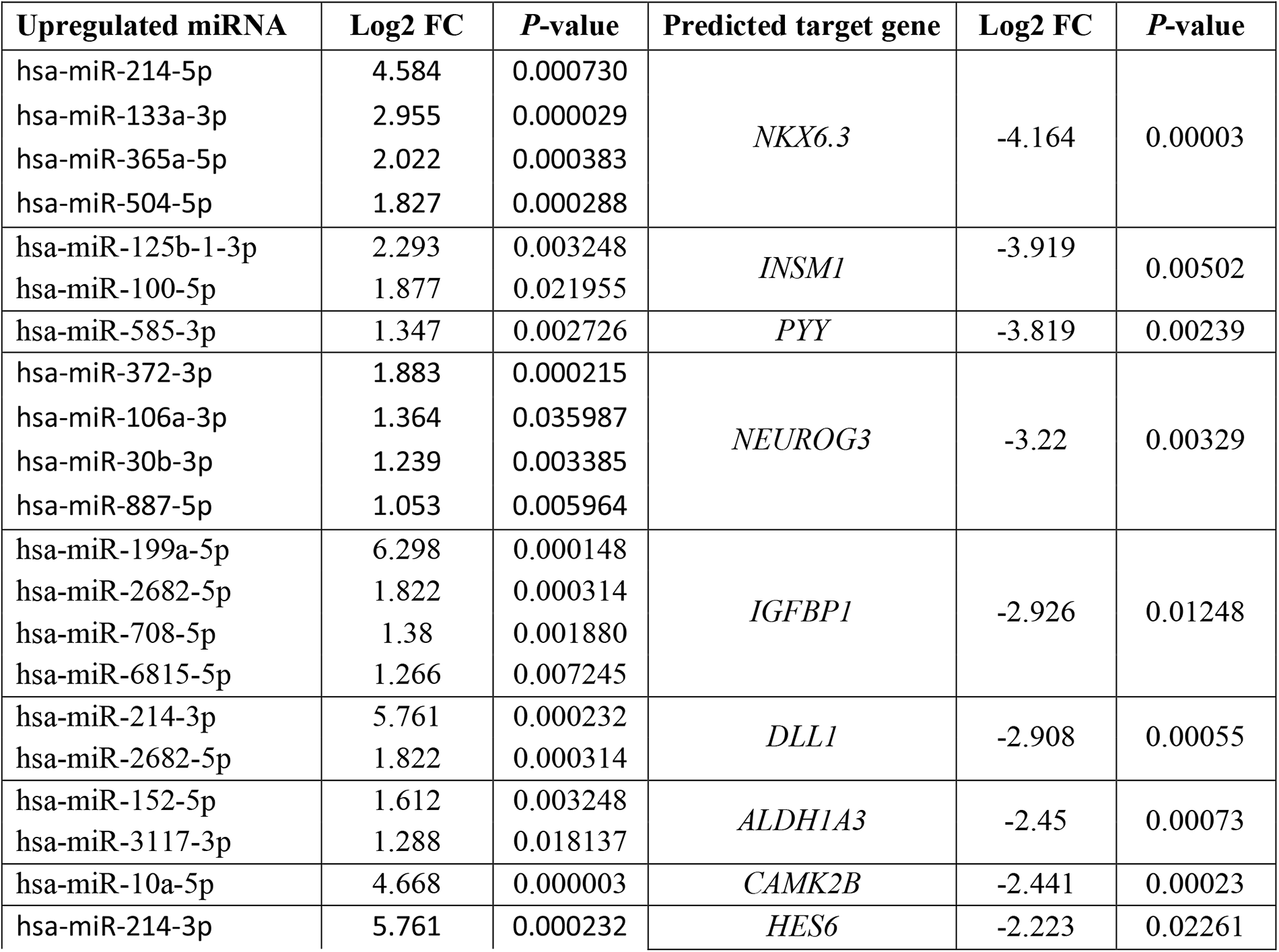
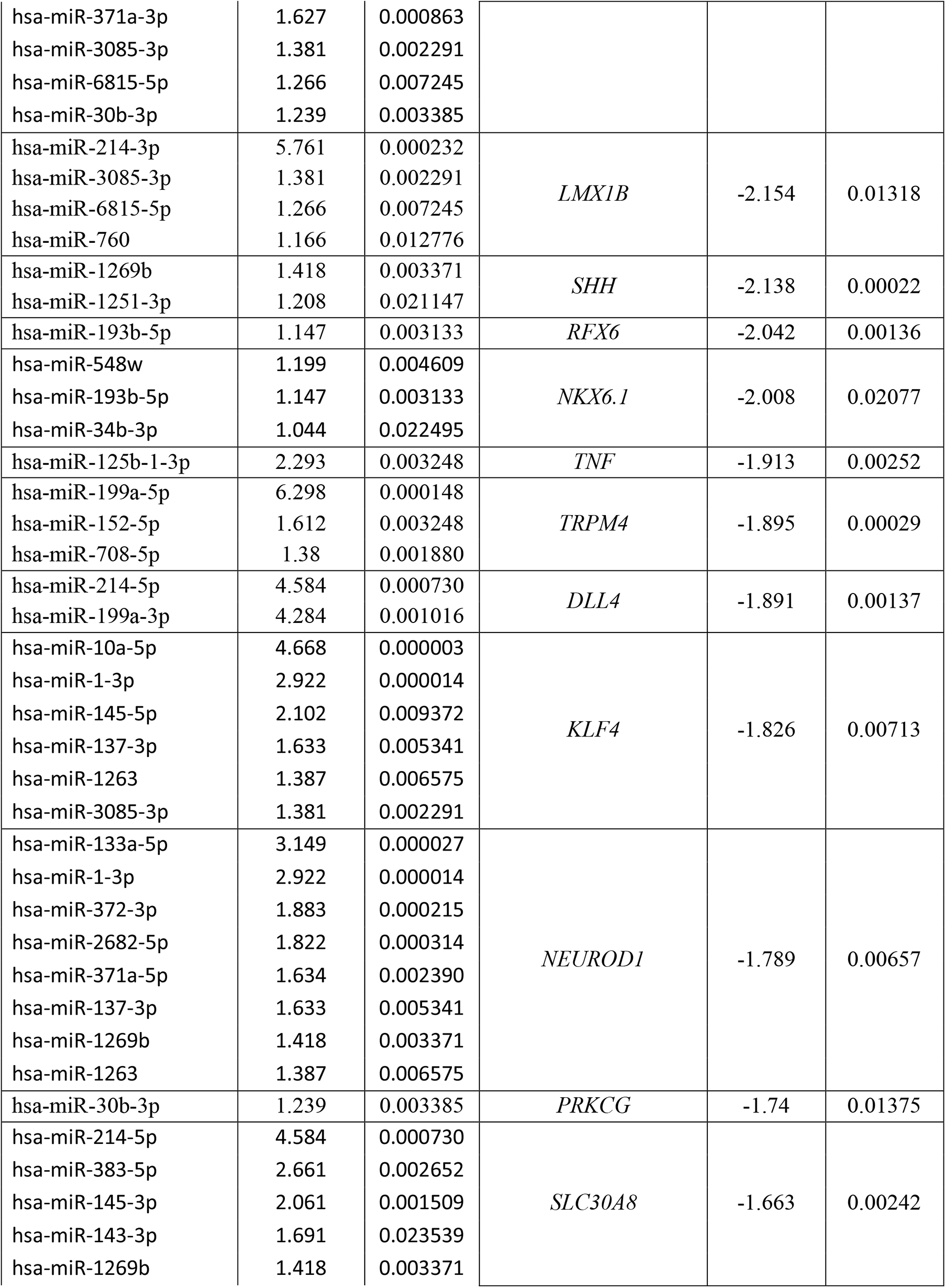
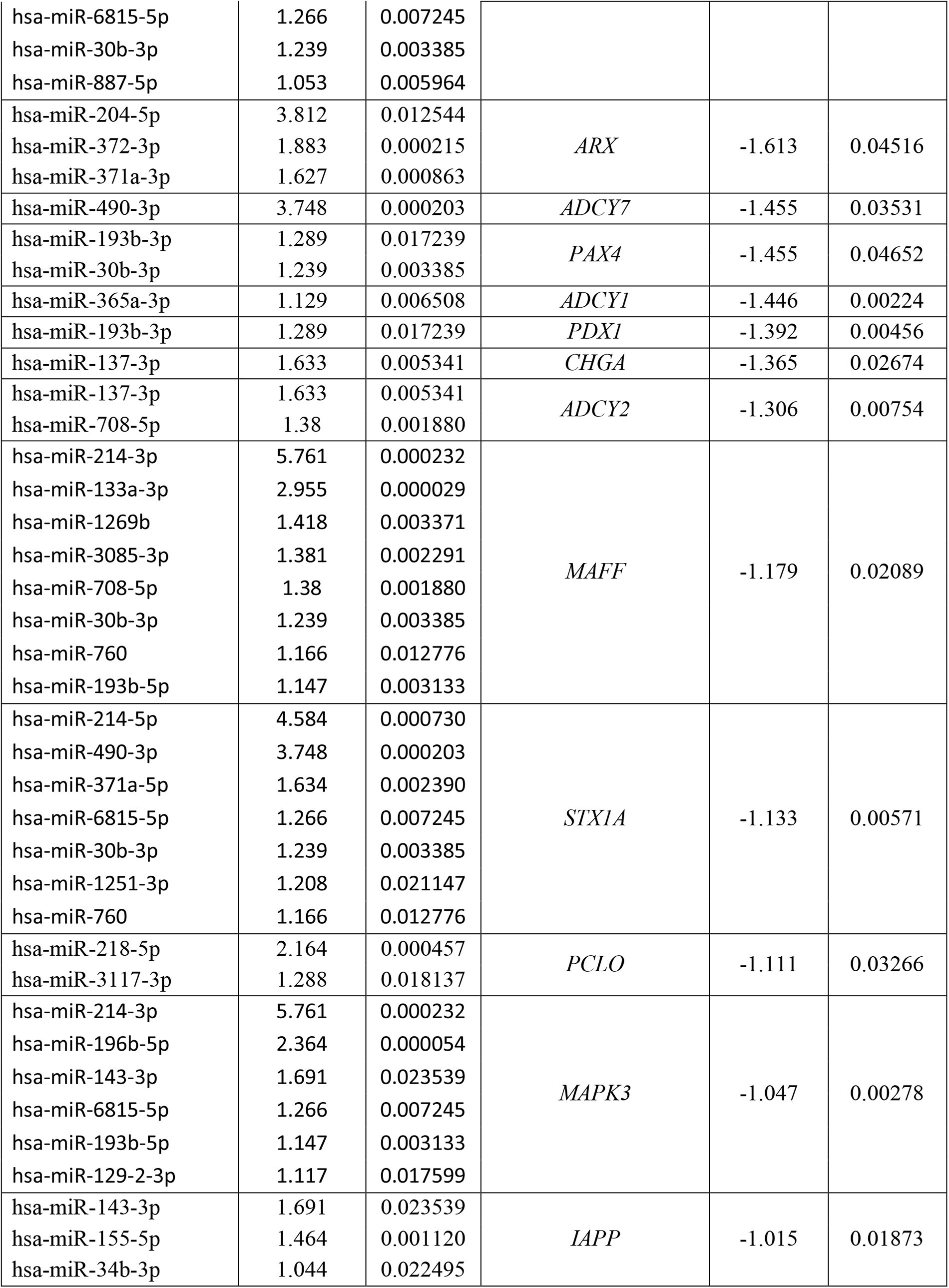
Upregulated DEmiRs (Log2 FC > 1, *P* < 0.05) and their predicted target DEGs (selected, Log2 FC < −1, *P* < 0.05) downregulated in *FOXA2*^−*/*−^ islets compared with WT-islets.

In addition to the TFs, other genes with diverse cellular roles, implicated in pancreatic islet and β-cell function, were also targeted by the significantly upregulated DEmiRs in the absence of FOXA2. For example, IGFBP1 was predicted to be targeted by hsa-miR-199a-5p, hsa-miR-2682-5p, hsa-miR-708-5p, and hsa-miR-6815-5p, while IAPP was predicted to be targeted by has-miR-143-3p, has-miR-155-5p, and has-miR-34b-3p. TRPM4 was targeted by hsa-miR-199a-5p, hsa-miR-152-5p, and hsa-miR-708-5p. SLC30A8, an important zinc transporter into insulin-containing granules [33, 34], was also targeted by a multitude of upregulated DEmiRs, including hsa-miR-214-5p, hsa-miR-383-5p, hsa-miR-145-3p, hsa-miR-143-3p, hsa-miR-1269b, hsa-miR-6815-5p, hsa-miR-30b-3p, and hsa-miR-887-5p. *PCLO*, which is known to be associated with diabetes [35], was predicted to be targeted by has-miR-218-5p and has-miR-3117-3p. The exocytotic gene [36, 37], *STX1A*, was also targeted by hsa-miR-214-5p, hsa-miR-490-3p, hsa-miR-371a-5p, hsa-miR-6815-5p, hsa-miR-30b-3p, hsa-miR-1251-3p, and hsa-miR-760. Furthermore, members of the adenylate cyclase family, including ADCY1, 2 and 7, which catalyze the cAMP generation, were targeted by hsa-miR-365a-3p, hsa-miR-137-3p, hsa-miR-708-5p, and hsa-miR-490-3p, respectively. Delta like canonical Notch ligands 1 and 4 (DLL1 and DLL4) were targeted by hsa-miR-214-3p and hsa-miR-2682-5p, and hsa-miR-214-5p and hsa-miR-199a-3p, respectively.

Numerous upregulated DEmiRs exhibited multiple predicted downregulated DEG targets, each with varying levels of prediction confidence. These prediction confidence levels, as determined by diverse sources within the IPA software, ranged from moderate to high and even extended to experimentally validated targets. For instance, the upregulated hsa-miR-371a-5p was experimentally confirmed to target STX1A, while both hsa-miR-145-5p and hsa-miR-10a-5p were experimentally validated to target KLF4. Additionally, hsa-miR-125b-1-3p was experimentally validated as a regulator of TNF. Moreover, 21 DEmiRs displayed high prediction levels for their target genes. Notably, the most upregulated hsa-miR-199a-5p was highly predicted to target *IGFBP1*. In this context, the miR-214 family, especially hsa-miR-214-3p, was found to target *DLL1*, *HES6*, *LMX1B*, *MAFF*, *MAPK3,* while hsa-miR-214-5p targeted *NKX6.3*, *DLL4*, *SLC30A8*, and *STX1A.* Furthermore, one significantly upregulated DEmiRs, hsa-miR-30b-3p, demonstrated regulatory influence on several critical genes, including *NEUROG3*, *PAX4*, *MAFF*, *HES6*, *STX1A*, *SLC30A8*, and *PRKCG*. In addition, hsa-miR-6815-5p, another significantly upregulated DEmiRs, exhibited a high number of DEG targets, encompassing *LMX1B*, *MAPK3*, *HES6*, *IGFBP1*, *STX1A*, and *SLC30A8*. Taken together, these findings strongly indicate the crucial involvement of the upregulated DEmiRs in regulating the expression of their target pancreatic genes and influencing the development and function of pancreatic islet cells. Nevertheless, it is essential to note that further experimental validation and functional studies are needed to solidify these predicted miRNA-mRNA networks.

### Downregulated DEmiRs potentially target upregulated genes in *FOXA2*^−/−^ islets

The cross-linked analysis between downregulated DEmiRs and upregulated DEGs in *FOXA2*^−/−^ islets revealed 79 significantly downregulated DEmiRs (Log2 FC < −1, *p* < 0.05), targeting 594 significantly upregulated DEGs (Log2 FC > 1, *p* < 0.05) in *FOXA2*^−/−^ islets. Notably, nervous system development associated upregulated DEGs were found among the highly predicted targets of the downregulated DEmiRs. 55 DEmiRs (Log2 FC < −1, *p* < 0.05) were found to be targeting 40 upregulated DEGs (Log2 FC > 2, *p* < 0.05) that are related to nervous system development (Fig. 5B and Supplementary Table 7). The most upregulated DEGs involved in this process, *RELN,* was predicted to be targeted by hsa-miR-429. Also, one of the most important DEGs for the neuron proliferation is *LIN28A* [38], which has been reported to ameliorate β-cell dysfunction and apoptosis [39]. *LIN28A* was targeted in our study by several DEmiRs such as hsa-let-7d-5p, hsa-miR-4510, hsa-miR-329-3p, hsa-miR-296-5p, and hsa-miR-181c-5p, which were downregulated in *FOXA2*^−/−^ islets. We noticed the upregulation of *EDNRB* gene, which is associated with hypoxia in pancreatic islets [40]. Our study showed clear downregulation of miRNAs targeting *EDNRB* in the absence of *FOXA2* such as hsa-miR-655-3p, hsa-miR-539-3p, hsa-miR-668-3p, hsa-miR-31-5p, and hsa-miR-340-5p. On the other hand, the same analysis revealed 34 significantly downregulated DEmiRs (Log2 FC < −1, *p* < 0.05) targeting 27 significantly upregulated DEGs (Log2 FC > 1, *p* < 0.05) in *FOXA2*^−/−^ islets associated with lipid and fat metabolism (Fig. 6B and Supplementary Table 8). For example, FOXA2 deletion was accompanied by downregulation in miRNAs targeting *APOC3* such as hsa-miR-429 and hsa-miR-4728-5p, and *APOC2* such as hsa-miR-4510 and hsa-miR-4728-5p. Upregulated *PDGFRA/B* were predicted targets of downregulated DEmiRs, including hsa-miR-654-5p, hsa-miR-892b, hsa-miR-146a-5p, and hsa-miR-301a-3p. Also, FOXA2 deletion led to the downregulation of miRNAs controlling the expression of the secretory phospholipase A2 (*PLA2G2A* and *PLA2G12B*), including hsa-miR-487a-5p, hsa-miR-4510, hsa-miR-4728-5p and hsa-miR-370-3p, hsa-miR-494-3p, hsa-miR-668-3p, and hsa-miR-892b.

**Fig. 5.**
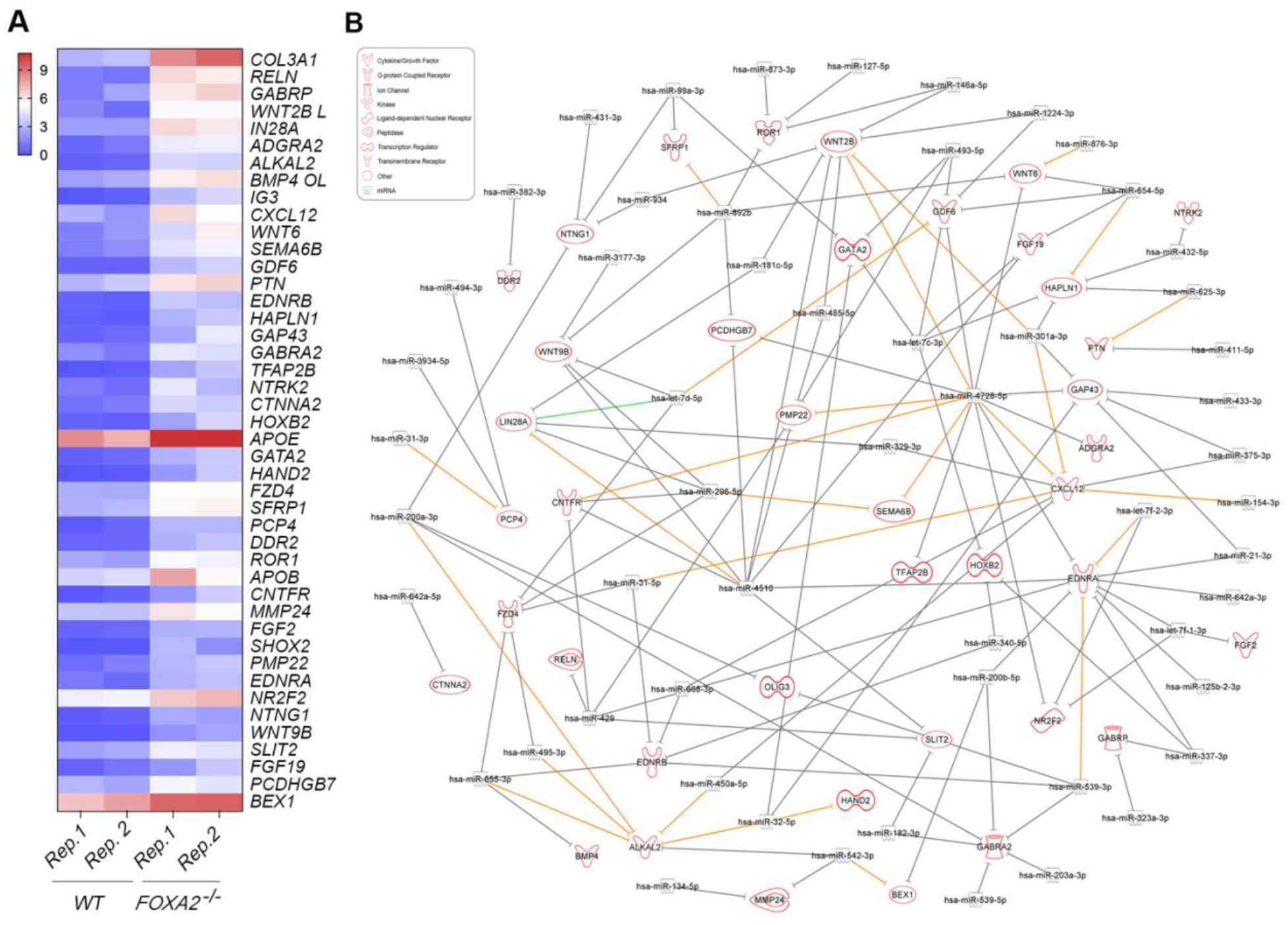
Lack of FOXA2 alters transcriptomic and miRNA profiles related to Nervous system development. **(A)** Heatmap of upregulated DEGs associated with nervous system development in *FOXA2^−/−^* islets compared to WT-islets (n=2). **(B)** Network reflecting target prediction analysis of downregulated DEmiRs and selected upregulated DEGs associated with nervous system development in *FOXA2^−/−^* islets. Green lines indicate experimentally observed targets, orange lines represent a high prediction confidence level, while grey lines indicate a moderate prediction confidence level.

**Fig. 6.**
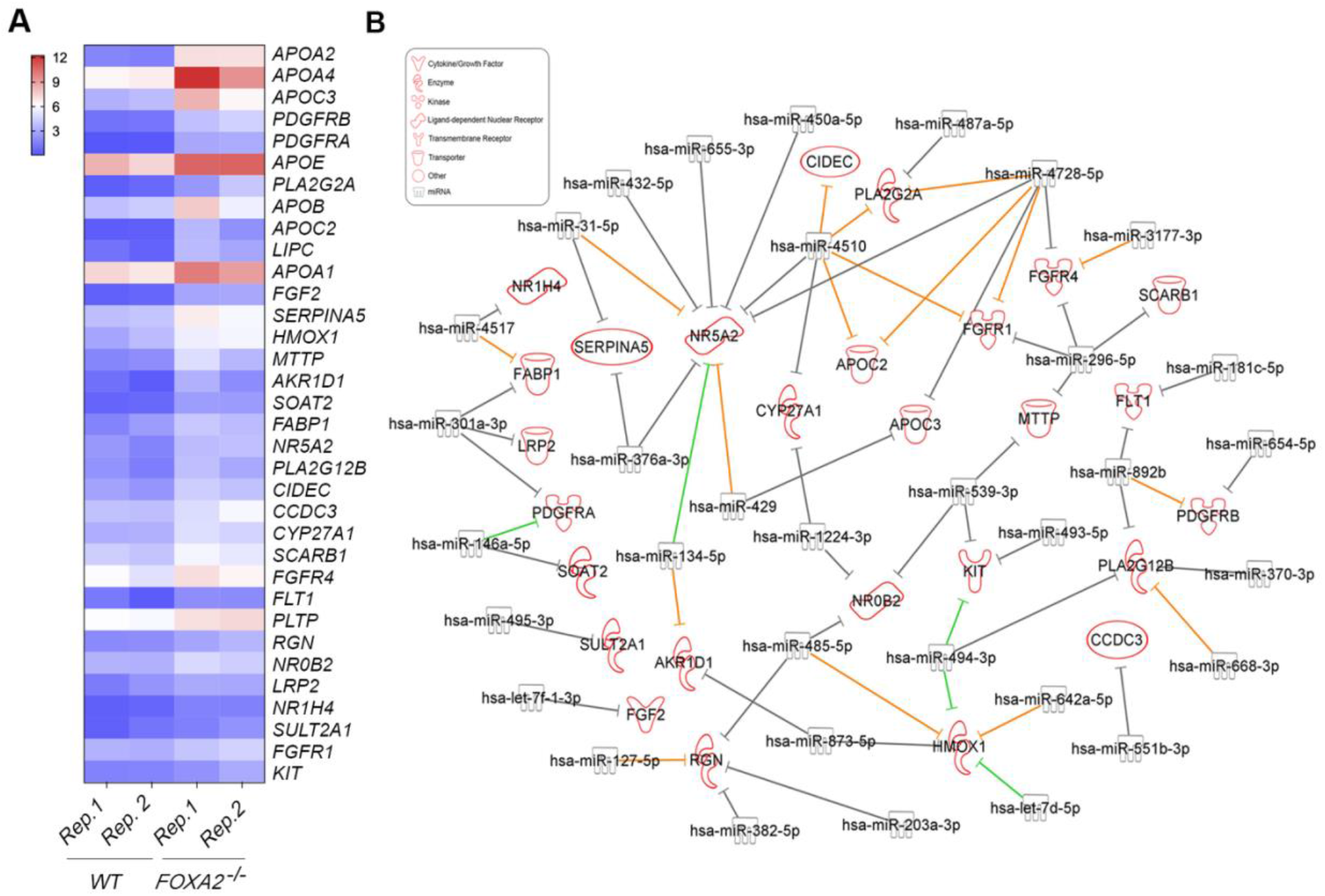
Loss of FOXA2 alters transcriptomic and miRNA profiles related to lipid metabolism. **(A)** Heatmap of upregulated DEGs associated with lipid metabolism in *FOXA2^−/−^*islets compared to WT-islets (n=2). **(B)** Network reflecting target prediction analysis of downregulated DEmiRs and selected upregulated DEGs associated with lipid metabolism in *FOXA2^−/−^* islets. Green lines indicate experimentally observed targets, orange lines represent a high prediction confidence level, while grey lines indicate a moderate prediction confidence level.

## Discussion

Our recent study reported that FOXA2 deficiency significantly reduces the number of pancreatic INS- and GCG-producing cells [18]. In agreement with these findings, RNA-Seq analysis of *FOXA2*^−/−^ islets has unveiled a substantial reduction in the expression of islet hormones and genes that play crucial roles in the development and function of pancreatic islet cells [2]. In this study, we identified several dysregulated miRNAs and mRNAs in pancreatic islets lacking FOXA2 and found that multiple DEmiRs were predicted to target several downregulated and upregulated DEGs. Conducting a pathway analysis on these target genes revealed an enrichment in biological processes, suggesting that miRNAs likely play significant roles in modulating pancreatic development and function. On the top list of enriched biological pathways affected by the downregulated DEGs in *FOXA2*^−/−^ islets, pancreatic islet development, glucose hemostasis, and diabetes were significantly enriched.

We identified a total of 45 out of 61 significantly upregulated DEmiRs, which showed significant degrees of prediction for targeting 32 downregulated pancreatic DEGs. Many of these potential targets of miRNAs, which were significantly downregulated by FOXA2 loss are known to impact insulin synthesis and secretion and islet development. These key targets include *PDX1, NKX6.1, PAX4, NKX6.3*, *INSM1*, *ARX, NEUROG3*, *NEUROD1*, *RFX6*, *MAFF*, *IGFBP1, TRPM4*, *PRKCG*, *SLC30A8*, *HES6, STX1A, DLL4, ADCY2, MAPK3*, *ALDH1A3, SHH,* and *IAPP.* Previous studies demonstrated that *IGFBP1* and *TRPM4,* playing critical roles in β-cell regeneration and insulin transport and secretion [41, 42], are predicted to be regulated by miR-199a-5p, which was the most upregulated miRNA in this study and is known to inhibit glucose metabolism [43]. Also, *NKX6.3*, *DLL4*, *SLC30A8*, and *STX1A*, which play key roles in insulin synthesis and release [33, 36, 44], are predicted to be targeted by miR-214-5p, which was significantly upregulated in FOXA2-deficient islets, and has been previously reported to be associated with type 1 diabetes (T1D) [45]. Furthermore, we found that *IGFBP1*, *TRPM4*, *ADCY2*, and *MAFF* were potential targets of miR-708-5p. It has been reported that increased miR-708 expression results in impaired insulin secretion and reduced proliferation of β-cells [9, 46]. Moreover, we observed a significant upregulation of miR-193b-3p, which was predicted to target *PAX4*, and *PDX1*. In addition, miR-193b-5p, was highly predicted to target *RFX6*, *NKX6.1*, *MAFF*, and *MAPK3*. Additionally, we observed an upregulation of miR-143-3p, which were predicted to target *SLC30A8*, *MAPK3*, and *IAPP*. Our study also identified several upregulated miRNAs associated with T1D, and T2D, including miR-372-3p [47], miR-100-5p [48], miR-152-5p [49], and miR-1269b [50]. In this study, miR-372-3p was predicted to target *NEUROG3*, *NEUROD1*, and *ARX*, while miR-100-5p was predicted to target *INSM1*, miR-152-5p targeted *ALDH1A3* and *TRPM4,* and hsa-miR-1269b was predicted to target *SHH*, *NEUROD1*, *SLC30A8*, and *MAFF*.

Previous studies have reported that dysregulations in miRNA expression are implicated in β-cell dysfunction and the autoimmune destruction of pancreatic β-cells in T1D, while also affecting insulin sensitivity, and glucose metabolism in T2D [51]. Furthermore, altered miRNA expression patterns serve as potential biomarkers for predicting and diagnosing diabetes [52]. Our study revealed that FOXA2 deletion resulted in upregulation in several miRNAs, which are known to be elevated in the blood of T1D and T2D patients. Increased levels of miR-199a-5p [53, 54], miR-199a-3p [55, 56], miR-145-5p [57, 58], miR-145-3p [59], and miR-143-3p [60, 61] have been associated with both T1D and T2D. Also, miR-100-5p is dysregulated in both T1D & T1D. [48, 56]. In T1D patients, multiple miRNAs have been shown to be increased such as hsa-miR-1-3p [57], miR-214 family, miR-152 family [62], miR-10a-5p [48], miR-618 [63], and miR-365a-3p [64]. Previous studies have indicated elevated levels of several miRNAs in the blood and tissues of T2D patients, such as miR-122-5p, miR-105-5p, and miR-193b-3p [65, 66], miR-125b [67, 68], miR-34b-3p [69], miR-206, miR-371a-5p, miR-133a [70–72], miR-204-5p [73, 74], and miR-133b [74, 75]. Also, some of the DEmiRs are associated with diabetes complications, such as miR-133a-3p, which is associated with myocardial steatosis in T2D patients [76], and miR-129-2-3p [77] and miR-129-5p, which have been detected in T2D patients with macrovascular complications [78]. Furthermore, hsa-miR-155-5p is closely associated with diabetic macular edema (DME) [79]. Several upregulated miRNAs are detected to be associated with diabetic nephropathy (DN) patients such as, miR-372-3p [47], miR-548ah-3p, miR-760 [80], and miR-1269b [81]. Increased expression of miR-196b-5p observed in serum of patients with diabetic kidney disease [82]. The identified miRNAs in our study play pivotal roles in various aspects of diabetes pathogenesis, impacting β-cell biology and insulin resistance in insulin target tissues. Some miRNAs influence glucose transporters expression, with miR-143 and miR-133b modulate GLUT4 protein expression, and miR-193b-3p inhibiting GLUT2 expression [61, 65, 72]. Also, other miRNAs are involved in diabetes by modulating key TFs, such as miR-204-5p, which downregulates MafA expression, and miR-372-3p, which targets growth factor-16 [47, 74]. Moreover, miR-199a-5p and miR-204-5p induce apoptosis, ROS generation, and ER stress in human islets [54, 74]. Notably, hsa-miR-1-3p exhibits a significant increase and strong association with glycated hemoglobin levels in patients diagnosed with T1D [57]. In our study, all of these miRNAs associated with T1D and/or T2D were significantly upregulated. This suggests that the absence of FOXA2 leads to an increase in the expression of diabetes-associated miRNAs, which could serve as biomarkers for predicting and diagnosing diabetes.

Although in this study we mainly focused on the downregulated DEGs associated with islet development and function, our findings demonstrated that several upregulated DEGs associated with nervous system development and lipid metabolism were predicted to be targeted by downregulated DEmiRs. This agrees with the previous study, which demonstrated that the deletion of Dicer, a critical factor in miRNA maturation, within PPs leads to significant impairments in the development of all islet cells, resulting in the upregulation of neuronal genes [83]. In our current study, we identified a significant downregulation of hsa-miR-375-3p in *FOXA2*^−/−^ islets compared to WT controls. In human islets, the expression of miR-375 is increased during development [84], and its inactivation leads to diabetes and reduces the pancreatic islet cell mass [30, 85]. Collectively, these findings suggest that the absence of FOXA2 disrupts the differentiation of pancreatic islets, partially by elevating and suppressing the expression of multiple miRNAs, thereby exerting a biological influence on pancreatic islet development through regulated key developmental genes. In addition, the data propose that pancreatic development is intricately governed by a complex miRNA network that targets crucial pancreatic and non-pancreatic genes during the pancreatic islet differentiation. This network enhances signaling pathways required for proper islet development.

## Conclusion

In conclusion, our findings demonstrate how the loss of FOXA2 disrupts the miRNA-mRNA regulatory network in iPSC-derived pancreatic islets, highlighting FOXA2’s crucial role in pancreatic endocrine cell generation and maturation. Our integration analysis revealed several dysregulated miRNAs targeting key pancreatic islet genes, shedding light on the intricate regulatory network controlling pancreatic development. While these insights are valuable, further validation studies are important to confirm the influence of miRNAs on pancreatic gene expression during islet differentiation. Moreover, deeper investigations into miRNA-mRNA interactions are essential for developing innovative diabetes therapeutic strategies.

## Acknowledgements

We thank the Genomic Core members at QBRI for their assistance for technical support in RNA and miRNA sequencing.

## Author contributions

A.K.E. and N.A. performed the experiments. A.K.E., N.A., and E.M.A. analyzed, interpreted the data, and wrote the manuscript. N.M.A. analyzed the RNA-Seq and miRNA-Seq data. All authors reviewed and approved the final version of the manuscript. E.M.A. conceived and designed the study and obtained research funding.

## Data Availability

The data that support the findings of this study are available from the corresponding author upon reasonable request.

## Ethical Approval

Not applicable.

## Consent to Participate

Not applicable.

## Consent for Publications

Not applicable.

