## Supplemental Figures and Tables for "Identifying miRNA Signatures Associated with Pancreatic Islet Dysfunction in a FOXA2-Deficient iPSC Model"

### SUPPLEMENTARY FIGURES AND TABLES

#### Supplementary Figure 1

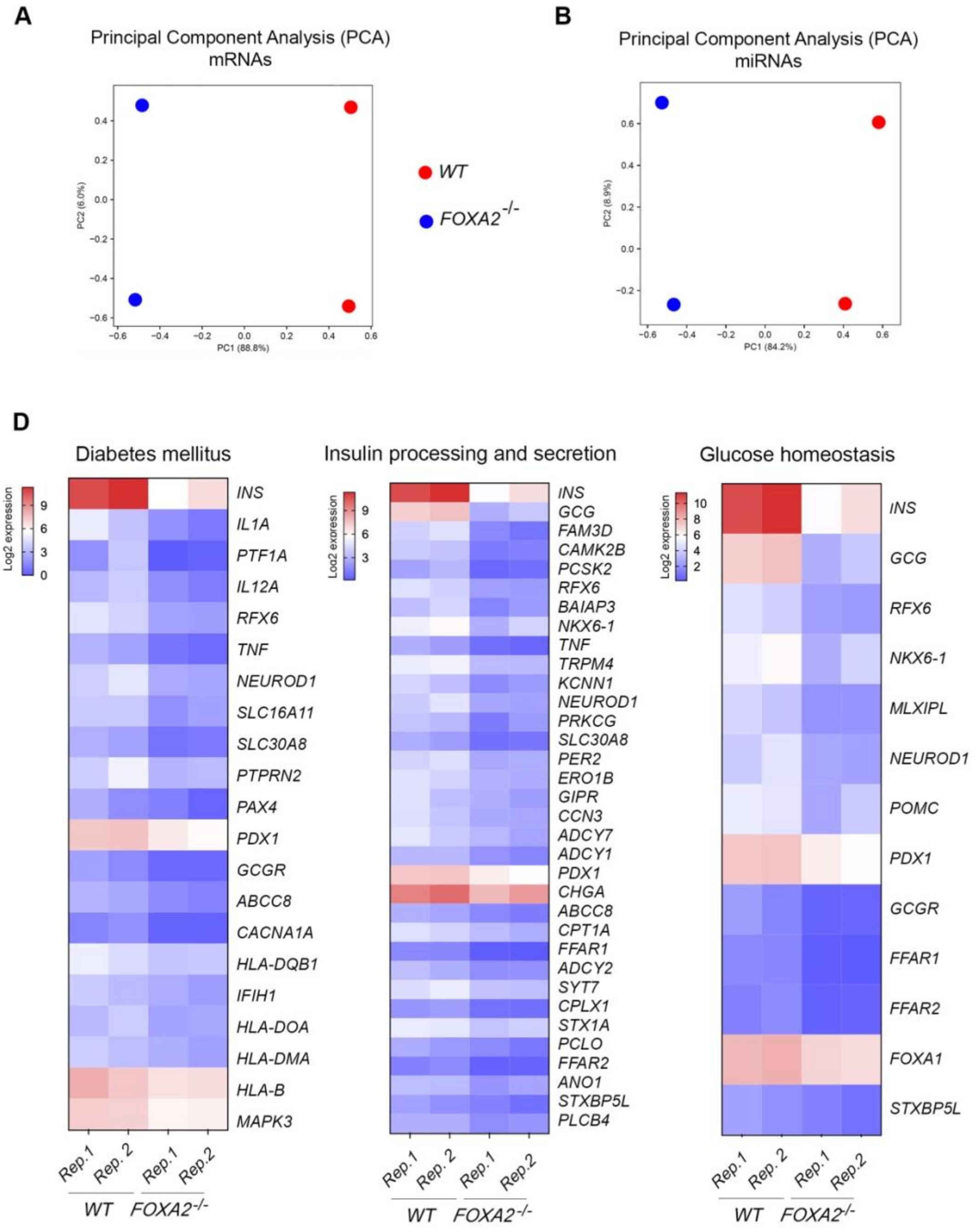


**Supplementary Figure 1. Enriched signaling pathways of downregulated DEGs in *FOXA2*$^{-/-}$ islets.** Principal component analyses (PCA) showing clear separation between DEGs **(A)** and DEmiRs **(B)** from *FOXA2*$^{-/-}$ islets and WT-islets. **(C)** Heatmaps of downregulated DEGs associated with diabetes mellitus, insulin processing, insulin secretion, and glucose homeostasis in FOXA2 absence using KEGG pathway and DAVID Tool.

**Supplementary Table 1.** Media formulations for pancreatic beta cell differentiation.

| Stage | Basal media |
|---|---|
| Stage 1 and 2 | MCDB 131 + 10 mM Glucose<br>1.5 g/L $NaHCO_3$<br>2 mM Glutamax<br>1% Pen/Strep<br>0.5% fatty acid free bovine serum albumin |
| Stage 3 and 4 | DMEM +<br>4.5g/l (25 mM) glucose<br>1% Pen/Strep<br>1% B27 supplement without vitamin A |
| Stage 5 | MCDB 131 + 20 mM Glucose<br>1.5 g/L $NaHCO_3$<br>2 mM Glutamax<br>1% Pen/Strep<br>2% fatty acid free bovine serum albumin<br>1:200 Insulin-Transferrin-Selenium-X (ITS)<br>10 μg/ mL Heparin<br>10 μM Zinc Sulfate |
| Stage 6 | MCDB 131 + 20 mM Glucose<br>1.5 g/L $NaHCO_3$<br>2 mM Glutamax<br>1% Pen/Strep<br>2% fatty acid free bovine serum albumin<br>1:200 Insulin-Transferrin-Selenium-X (ITS)<br>10 μg/ mL Heparin<br>10 μM Zinc Sulfate<br>1% non-Essential amino acids MEM-NEAA |
| Stage 7 | MCDB 131 +<br>1.5 g/L $NaHCO_3$<br>2 mM Glutamax<br>1% Pen/Strep<br>2% fatty acid free bovine serum albumin<br>1:200 Insulin-Transferrin<br>10 μg/ mL Heparin<br>10 μM Zinc Sulfate<br>1% non-Essential amino acids MEM-NEAA<br>1 mM N-acetyl cysteine |

| Stage | Cytokine | Final Concentration |
|---|---|---|
| Stage 1 day 1 | Activin A | 100 ng/ml |
| | CHIR99021 | 3 μM |
| | Rock inhibitor | 10 μM |
| | Vitamin C | 0.25 mM |
| Stage 1 day 2-4 | Activin A | 100 ng/ml |
| | Vitamin C | 0.25 mM |
| Stage 2 (2 days) | FGF10 | 50 ng/ml |
| | LDN | 0.2 μM |

| | | |
|---|---|---|
| | WNT3a | 3 ng/ml |
| | Vitamin C | 0.25 mM |
| Stage 3 (2 days) | LDN | 0.2 µM |
| | FGF10 | 50 ng/ml |
| | Retinoic acid | 2 µM |
| | Sant-1 | 0.25 µM |
| | Vitamin C | 0.25 mM |
| Stage 4 (4 days) | LDN | 0.2 µM |
| | EGF | 100 ng/ml |
| | Nicotinamide | 10 mM |
| | Vitamin C | 0.25 mM |
| Stage 5 (3 days) | SANT1 | 0.25 µM |
| | LDN | 0.1 µM |
| | RA | 0.05 µM |
| | Alk5i II | 10 µM |
| | T3 | 1 µM |
| | Vitamin C | 0.25 mM |
| Stage 6 (7 days) | LDN | 0.1 µM |
| | Alk5i II | 10 µM |
| | T3 | 1 µM |
| | Gamma-secretase inhibitor | 1 µM |
| | Vitamin C | 0.25 mM |
| Stage 7 (7-14 days) | Betacellulin | 20 ng/ mL |
| | Alk5i II | 10 µM |
| | T3 | 1 µM |
| | Gamma-secretase inhibitor | 1 µM |
| | Vitamin C | 0.25 mM |

**Supplementary Table 2.** List of primers and miRNA assays used for RT-qPCR validation.

| **List of primers** | | |
|---|---|---|
| **Gene** | **Forward** | **Reverse** |
| *INS* | AAGAGGCCATCAAGCAGATCA | CAGGAGGCGCATCCACA |
| *GCG* | CTCTTCACCTGCTCTGTTCTAC | TGGATTTCTCCTCTGTGTCTTG |
| *PPY* | AGGTGCTCGCTTGGTCTAGTG | ACCCAGCAGTGGCTGTAGTAAC |
| *IAPP* | TTGAGAAGCAATGGGCATCC | GGGTGTAGCTTTCAGATGGTTC |
| *ARX* | CTGCTGAAACGCAAACAGAGGC | CTCGGTCAAGTCCAGCCTCATG |
| *NKX6.1* | GGGCTCGTTTGGCCTATTCGTT | CCACTTGGTCCGGCGGTTCT |
| *PDX1* | CGTCCAGCTGCCTTTCCCAT | CCGTGAGATGTACTTGTTGAATAGGA |
| *GCGR* | AGGTGATGGACTTCCTGTTTGAG | TACTTGTCGAAGGTTCTGTTGC |
| *UCN3* | GATGGGCTTGGCTTTGTAGA | GGAGGGAAGTCCACTCTCG |
| *NEUROG3* | GGCTGTGGGTGCTAAGGGTAAG | CAGGGAGAAGCAGAAGGAACAA |
| *NEUROD1* | GCCCCAGGGTTATGAGACTAT | GAGAACTGAGACACTCGTCTGT |
| *NKX2.2* | AAACCATGTCACGCGCTCA | GGCGTTGTACTGCATGTGCT |
| *INSM1* | TTTGTCTCGTGGTTGGAAGC | CCAAAACAACCCGTACGCTA |
| *PAX4* | AGCAGAGGCACTGGAGAAAGAGTT | CAGCTGCATTTCCCACTTGAGCTT |
| *PAX6* | GCGGAAGCTGCAAAGAAATAG | GGGCAAACACATCTGGATAATG |
| *RFX6* | GTCGATGCATGGCTTGGACT | TGGGCCATAGCTAGACGGTG |
| *HES1* | AGTGAAGCACCTCCGGAAC | TCACCTCGTTCATGCACTC |
| *HES6* | AGCCCCTGGTGGAGAAGA | CAGCACTCCGGCGTTCTC |
| *PTF1A* | CCAGAAGGTCATCATCTGCC | AGAGAGTGTCCTGCTAGGGG |
| *ABCC6* | TGGATCATAGTGCTGGCAAATG | GTTTCTCCTTCTCCTCTATCTCC |
| *AFP* | ACAGAGGAACAACTTGAGGCTGTC | AGCAAAGCAGACTTCCTGTTCCTG |
| *APOA2* | GTTCGGAGACAGGCAAAGGA | TCAAAGTAAGACTTGGCCTCGG |
| *APOC3* | CTTCATGCAGGGTTACATGAAG | TTTCAGGGAACTGAAGCCATC |
| *HHEX* | GCGAGAGACAGGTCAAAACC | AGGGCGAACATTGAGAGCTA |
| *LIN28A* | GGTGCGGGCATCTGTAAGTG | GGAACCCTTCCATGTGCAGC |
| *NPY* | CGCTGCGACACTACATCAACC | AGGGTCTTCAAGCCGAGTTCTG |
| *OLIG3* | GCGAGAGAGATCAAGACCCA | ACTGCGTCGAAAGGAAAACC |
| *SST* | AGCTGCTGTCTGAACCCAAC | CCATAGCCGGGTTTGAGTTA |
| *GAPDH* | ACGACCACTTTGTCAAGCTCATTTC | GCAGTGAGGGTCTCTCTCTTCCTCT |

| **miRNA Assays** | |
|---|---|
| **miRNA PCR Assay** | **Assay ID** |
| hsa-miR-199a-5p | YP00204494 |

| | |
|---|---|
| hsa-miR-199a-3p | YP00204536 |
| hsa-miR-204-5p | YP00206072 |
| hsa-miR-1-3p | YP00204344 |
| hsa-miR-383-5p | YP00205904 |
| hsa-miR-10b-5p | YP00205637 |
| hsa-miR-218-5p | YP00206034 |
| hsa-miR-137-3p | YP00206062 |
| hsa-miR-155-5p | YP02119311 |
| hsa-miR-105-5p | YP00204389 |
| hsa-miR-337-3p | YP00205938 |
| hsa-miR-542-3p | YP00205444 |
| hsa-miR-340-5p | YP00206068 |
| hsa-miR-203a-3p | YP00205914 |
| hsa-miR-99a-3p | YP00204523 |
| hsa-miR-409-3p | YP00204358 |
| hsa-miR-934 | YP02119292 |
| hsa-miR-136-3p | YP00205503 |
| hsa-let-7f-2-3p | YP00204095 |
| hsa-miR-203b-3p | YP02103829 |
| hsa-miR-98-5p | YP00204640 |
| hsa-miR-493-3p | YP00204557 |
| hsa-miR-122-5p | YP00205664 |
| hsa-miR-371a-3p | YP00204299 |
| hsa-miR-371a-5p | YP00204493 |
| hsa-miR-373-3p | YP00204604 |
| SNORD48(hsa) | YP00203903 |

**Supplementary Table 3.** Top upregulated DEGs in *FOXA2*$^{-/-}$ islets compared with WT-islets (Log2 FC > 1, *P* < 0.05).

| **Gene ID** | **Log2 FC** | ***P*-value** |
|---|---|---|
| *KRT1* | 5.883013 | 0.000985 |
| *COL3A1* | 5.782581 | 0.000195 |
| *CALB1* | 5.479372 | 0.000017 |
| *APOA2* | 5.433256 | 0.000005 |
| *A2M* | 5.266395 | 0.009813 |
| *RELN* | 5.035494 | 0.000034 |
| *SOX17* | 4.794410 | 0.000411 |
| *COL1A2* | 4.736296 | 0.000725 |
| *FGG* | 4.704527 | 0.000107 |
| *AFP* | 4.526164 | 0.000698 |
| *GABRP* | 4.445883 | 0.000942 |
| *PTX3* | 4.402107 | 0.000166 |
| *CXCL14* | 4.340724 | 0.019216 |
| *APOA4* | 4.278651 | 0.014152 |
| *POSTN* | 4.219276 | 0.000294 |
| *HSD3B1* | 4.138513 | 0.002163 |
| *SPARCL1* | 4.125078 | 0.000231 |
| *WNT2B* | 4.066964 | 0.000359 |
| *LIN28A* | 4.004317 | 0.000051 |
| *ACTA2* | 3.986646 | 0.003101 |
| *CD248* | 3.980332 | 0.000324 |
| *ADGRA2* | 3.913052 | 0.000089 |
| *APOC3* | 3.897227 | 0.006166 |
| *MYBPC3* | 3.847898 | 0.000024 |
| *AHSG* | 3.843007 | 0.000050 |
| *KCNJ8* | 3.741418 | 0.000881 |
| *C6* | 3.725187 | 0.000042 |
| *RBP4* | 3.724246 | 0.000873 |
| *COL11A1* | 3.710116 | 0.000227 |
| *ACTC1* | 3.669441 | 0.002052 |
| *BGN* | 3.665861 | 0.001290 |
| *ALKAL2* | 3.619271 | 0.000041 |
| *HAND1* | 3.618144 | 0.007086 |
| *NPY* | 3.603045 | 0.001557 |
| *GATA5* | 3.581672 | 0.000418 |
| *GPC3* | 3.570344 | 0.001960 |
| *DUSP9* | 3.558037 | 0.000126 |
| *BMP4* | 3.557951 | 0.000201 |

| | | |
|---|---|---|
| *COL9A3* | 3.519734 | 0.000177 |
| *LCN15* | 3.506077 | 0.002915 |
| *DES* | 3.505785 | 0.002163 |
| *CDKN1C* | 3.481887 | 0.000074 |
| *DCN* | 3.477796 | 0.002240 |
| *CLMP* | 3.463822 | 0.000077 |
| *VAT1L* | 3.450420 | 0.000271 |
| *OLIG3* | 3.438869 | 0.000910 |
| *CXCL12* | 3.427706 | 0.003097 |
| *COL6A3* | 3.366678 | 0.002987 |
| *URAD* | 3.360436 | 0.000193 |
| *TTYH1* | 3.323021 | 0.001042 |
| *ASGR2* | 3.297619 | 0.000044 |
| *HTRA1* | 3.291037 | 0.007451 |
| *F2* | 3.257104 | 0.004818 |
| *WNT6* | 3.256271 | 0.010484 |
| *PITX2* | 3.241344 | 0.000034 |
| *SFRP5* | 3.205634 | 0.003585 |
| *FGB* | 3.147667 | 0.033733 |
| *SEMA6B* | 3.141794 | 0.000401 |
| *COL5A2* | 3.140943 | 0.000436 |
| *TGFBI* | 3.119701 | 0.006841 |
| *LGR5* | 3.099138 | 0.000449 |
| *GDF6* | 3.087368 | 0.000591 |
| *SRGN* | 3.075938 | 0.001392 |
| *PTN* | 3.070695 | 0.000677 |
| *EDNRB* | 3.070232 | 0.000141 |
| *PTH1R* | 3.041565 | 0.000053 |
| *PDGFRB* | 3.034967 | 0.000385 |
| *HAPLN1* | 3.029612 | 0.000470 |
| *KRTDAP* | 3.023072 | 0.043904 |
| *GAP43* | 3.021519 | 0.020357 |
| *ZNF521* | 3.010242 | 0.000458 |
| *DSG1* | 3.006862 | 0.003137 |
| *PDGFRA* | 2.958354 | 0.000049 |
| *MYL7* | 2.952391 | 0.009990 |
| *GLDC* | 2.927665 | 0.000490 |
| *GABRA2* | 2.926765 | 0.000792 |
| *HHIP* | 2.916994 | 0.000907 |
| *LGALS14* | 2.900540 | 0.033138 |
| *HMGCS2* | 2.891864 | 0.001677 |

| | | |
|---|---|---|
| *UGT2B11* | 2.879645 | 0.007328 |
| *FGA* | 2.869400 | 0.003365 |
| *TFAP2B* | 2.867028 | 0.002359 |
| *B3GALT1* | 2.864150 | 0.000771 |
| *NTRK2* | 2.860824 | 0.007615 |
| *OCA2* | 2.857906 | 0.000306 |
| *EMILIN1* | 2.854771 | 0.017253 |
| *CTNNA2* | 2.852362 | 0.000295 |
| *ANPEP* | 2.850059 | 0.001980 |
| *HOXB2* | 2.841496 | 0.008615 |
| *PCSK5* | 2.826800 | 0.000312 |
| *AMBP* | 2.817976 | 0.026671 |
| *PENK* | 2.814065 | 0.004658 |
| *APOE* | 2.811868 | 0.001774 |
| *GATA2* | 2.806731 | 0.001192 |
| *SLC22A10* | 2.804778 | 0.009672 |
| *ERVV-1* | 2.802207 | 0.000203 |
| *CRABP1* | 2.798465 | 0.049272 |
| *APCS* | 2.796480 | 0.001975 |
| *HSPB6* | 2.795951 | 0.001635 |
| *HAND2* | 2.788982 | 0.008251 |
| *IGFBP3* | 2.783111 | 0.002383 |
| *PLA2G2A* | 2.781383 | 0.012291 |
| *CA4* | 2.776809 | 0.012719 |
| *GAS1* | 2.775482 | 0.015586 |
| *FZD4* | 2.772512 | 0.000062 |
| *COL9A2* | 2.746627 | 0.000854 |
| *ADCYAP1* | 2.725630 | 0.000704 |
| *TMEM130* | 2.711961 | 0.004430 |
| *CAPN6* | 2.694053 | 0.000100 |
| *ACTG2* | 2.691711 | 0.008602 |
| *SFRP1* | 2.668163 | 0.000147 |
| *PCP4* | 2.650810 | 0.000294 |
| *DDR2* | 2.638976 | 0.000618 |
| *DPP4* | 2.635987 | 0.023592 |
| *TBX2* | 2.635329 | 0.001674 |
| *CNN1* | 2.632396 | 0.001149 |
| *TRABD2B* | 2.629211 | 0.000479 |
| *LOXL2* | 2.611407 | 0.000978 |
| *DAAM2* | 2.610841 | 0.001722 |
| *LOX* | 2.608137 | 0.000210 |

| | | |
|---|---|---|
| *MRC2* | 2.598063 | 0.002786 |
| *NRP2* | 2.595536 | 0.004361 |
| *TSPAN18* | 2.595432 | 0.000093 |
| *TPM2* | 2.593790 | 0.001563 |
| *ROR1* | 2.588043 | 0.000290 |
| *CST1* | 2.577324 | 0.000219 |
| *APOB* | 2.569907 | 0.031380 |
| *LIN7A* | 2.567268 | 0.000076 |
| *NKX1-2* | 2.564435 | 0.002282 |
| *CNTFR* | 2.555445 | 0.019535 |
| *APOC2* | 2.534667 | 0.008866 |
| *PDZK1* | 2.521182 | 0.019898 |
| *SPSB4* | 2.521021 | 0.000595 |
| *SLC19A3* | 2.520547 | 0.005894 |
| *LIPC* | 2.506663 | 0.002433 |
| *MYL4* | 2.504500 | 0.019959 |
| *SLC17A4* | 2.504090 | 0.005968 |
| *COL8A1* | 2.497091 | 0.000683 |
| *CPA2* | 2.492019 | 0.003022 |
| *CRB2* | 2.489605 | 0.000451 |
| *SBSPON* | 2.487491 | 0.003741 |
| *TNC* | 2.476923 | 0.000286 |
| *SALL3* | 2.471260 | 0.000098 |
| *KCNIP1* | 2.466084 | 0.009811 |
| *PRSS35* | 2.461024 | 0.000100 |
| *TSPAN8* | 2.441167 | 0.002133 |
| *TTR* | 2.434215 | 0.004597 |
| *PCDH17* | 2.430816 | 0.000176 |
| *MATN2* | 2.429981 | 0.000298 |
| *APOA1* | 2.415987 | 0.004667 |
| *MMP24* | 2.408768 | 0.000790 |
| *NEFM* | 2.407282 | 0.000774 |
| *FGF2* | 2.402290 | 0.000148 |
| *ENG* | 2.399158 | 0.000093 |
| *PKDCC* | 2.389005 | 0.005266 |
| *SCG2* | 2.388477 | 0.002145 |
| *CTHRC1* | 2.380752 | 0.006882 |
| *SELENOP* | 2.379562 | 0.000788 |
| *SERPINA5* | 2.376276 | 0.002514 |
| *DACT1* | 2.366551 | 0.003592 |
| *IGDCC3* | 2.360696 | 0.002721 |

| | | |
|---|---|---|
| *SHOX2* | 2.352232 | 0.009379 |
| *ENPP7* | 2.347244 | 0.035730 |
| *OLFML3* | 2.346314 | 0.002786 |
| *NPR3* | 2.342991 | 0.007742 |
| *HMOX1* | 2.339864 | 0.002408 |
| *MTTP* | 2.339240 | 0.011627 |
| *TIMP3* | 2.333180 | 0.001451 |
| *CD244* | 2.320870 | 0.031767 |
| *LRRTM1* | 2.317746 | 0.005312 |
| *EDN3* | 2.314412 | 0.021595 |
| *NPTX2* | 2.287396 | 0.006995 |
| *IGLON5* | 2.287204 | 0.006647 |
| *PMP22* | 2.283389 | 0.002400 |
| *NEFL* | 2.282019 | 0.004551 |
| *HOXB4* | 2.274054 | 0.002251 |
| *CLU* | 2.273067 | 0.006682 |
| *CST4* | 2.272256 | 0.000108 |
| *CYTL1* | 2.262476 | 0.015040 |
| *UGT2A3* | 2.252964 | 0.016734 |
| *PAH* | 2.238536 | 0.020612 |
| *PF4* | 2.237150 | 0.001896 |
| *TF* | 2.230508 | 0.001212 |
| *ELOVL2* | 2.225563 | 0.002885 |
| *EDNRA* | 2.220325 | 0.001145 |
| *RGS5* | 2.219339 | 0.001927 |
| *CLDN2* | 2.211100 | 0.004140 |
| *FBN1* | 2.209027 | 0.000430 |
| *BTNL3* | 2.202721 | 0.005767 |
| *NR2F2* | 2.198754 | 0.000651 |
| *CUX2* | 2.193826 | 0.042671 |
| *NTNG1* | 2.191506 | 0.000599 |
| *PREX1* | 2.191205 | 0.025568 |
| *FAM151A* | 2.188458 | 0.001172 |
| *SDK2* | 2.178335 | 0.001066 |
| *ITGA11* | 2.174331 | 0.002998 |
| *HOGA1* | 2.168276 | 0.002259 |
| *FLRT2* | 2.164832 | 0.046475 |
| *BAMBI* | 2.159477 | 0.000207 |
| *MMP2* | 2.157184 | 0.016318 |
| *LZTS1* | 2.147587 | 0.003166 |
| *SAMD11* | 2.147429 | 0.005941 |

| | | |
|---|---|---|
| *DSC3* | 2.144961 | 0.001131 |
| *DKK2* | 2.140417 | 0.009316 |
| *LRRN2* | 2.139285 | 0.000134 |
| *WNT9B* | 2.130923 | 0.001330 |
| *SLIT2* | 2.123351 | 0.000898 |
| *CTSC* | 2.114790 | 0.000272 |
| *SDC2* | 2.111606 | 0.014508 |
| *LAMA2* | 2.102461 | 0.000191 |
| *ANXA13* | 2.098008 | 0.004191 |
| *TUBB4A* | 2.097017 | 0.008472 |
| *FGF19* | 2.086062 | 0.026446 |
| *LUM* | 2.085002 | 0.011922 |
| *FST* | 2.082178 | 0.049763 |
| *FBLN5* | 2.076190 | 0.012916 |
| *PHYHIPL* | 2.065911 | 0.003713 |
| *PCDHGB7* | 2.061351 | 0.002564 |
| *TRH* | 2.061298 | 0.000752 |
| *BEX1* | 2.061178 | 0.003811 |
| *AKR1D1* | 2.047309 | 0.021353 |
| *MSN* | 2.046045 | 0.009079 |
| *PRTG* | 2.040664 | 0.007605 |
| *APCDD1* | 2.031217 | 0.014344 |
| *COL1A1* | 2.020142 | 0.006933 |
| *MGP* | 2.010206 | 0.031444 |
| *ADAMTS12* | 2.006529 | 0.000522 |
| *MMP16* | 2.004701 | 0.017031 |
| *SLC47A1* | 1.995437 | 0.000297 |
| *SLIT3* | 1.992052 | 0.000376 |
| *BAAT* | 1.987333 | 0.001823 |
| *NLGN4X* | 1.985241 | 0.000748 |
| *DACT2* | 1.981547 | 0.000480 |
| *TLX3* | 1.980741 | 0.000286 |
| *MAN1C1* | 1.979755 | 0.000433 |
| *NID2* | 1.961793 | 0.000978 |
| *GASK1B* | 1.961490 | 0.002402 |
| *SOAT2* | 1.950445 | 0.000241 |
| *LGALS2* | 1.942708 | 0.014338 |
| *UCHL1* | 1.941570 | 0.011738 |
| *SYDE1* | 1.940255 | 0.009649 |
| *ALPI* | 1.938137 | 0.022791 |
| *MCAM* | 1.938045 | 0.000854 |

| | | |
|---|---|---|
| *MYH6* | 1.937293 | 0.013701 |
| *GUCY2C* | 1.936311 | 0.012999 |
| *RHOJ* | 1.931813 | 0.000334 |
| *PDGFD* | 1.928530 | 0.000457 |
| *SYNPO2* | 1.927985 | 0.000906 |
| *GDF7* | 1.927483 | 0.001684 |
| *SLCO2B1* | 1.913000 | 0.005740 |
| *TBX18* | 1.910745 | 0.002291 |
| *CNTN6* | 1.906387 | 0.000490 |
| *L1CAM* | 1.901132 | 0.007481 |
| *DPEP1* | 1.899975 | 0.005106 |
| *ACTN3* | 1.892337 | 0.003327 |
| *COL2A1* | 1.884383 | 0.000490 |
| *DCHS2* | 1.879436 | 0.013558 |
| *NRK* | 1.877832 | 0.002455 |
| *KANK4* | 1.877680 | 0.011867 |
| *C7* | 1.873867 | 0.001195 |
| *LRRC32* | 1.863560 | 0.000418 |
| *SEMA5A* | 1.859240 | 0.007233 |
| *EGF* | 1.858628 | 0.002687 |
| *CTSF* | 1.856693 | 0.000620 |
| *ST6GAL2* | 1.851025 | 0.025164 |
| *FHL1* | 1.847895 | 0.034701 |
| *HOXB3* | 1.846972 | 0.003910 |
| *TENM3* | 1.846714 | 0.000394 |
| *LEFTY1* | 1.840215 | 0.001248 |
| *FAM20A* | 1.838669 | 0.026073 |
| *WNT5A* | 1.836846 | 0.011571 |
| *PAGE4* | 1.833917 | 0.027329 |
| *WSCD2* | 1.830264 | 0.027807 |
| *SLC5A9* | 1.827910 | 0.010260 |
| *ALDH1A1* | 1.827323 | 0.001934 |
| *CA14* | 1.825735 | 0.006133 |
| *DCDC2* | 1.825383 | 0.001056 |
| *KITLG* | 1.822008 | 0.003114 |
| *FABP1* | 1.817167 | 0.009621 |
| *AQP10* | 1.815680 | 0.004675 |
| *TFEC* | 1.814297 | 0.028170 |
| *NAALAD2* | 1.814110 | 0.000846 |
| *PLAT* | 1.813647 | 0.003309 |
| *ORM1* | 1.805846 | 0.005113 |

| | | |
|---|---|---|
| *HEG1* | 1.799395 | 0.033885 |
| *BNC1* | 1.797216 | 0.001933 |
| *SOSTDC1* | 1.795888 | 0.020262 |
| *MSX2* | 1.791604 | 0.013408 |
| *KCNQ1* | 1.791418 | 0.009810 |
| *LEFTY2* | 1.786042 | 0.000464 |
| *RSPO3* | 1.781372 | 0.016298 |
| *PTGIS* | 1.776587 | 0.000727 |
| *L1TD1* | 1.772977 | 0.000378 |
| *CSPG4* | 1.766219 | 0.009690 |
| *CPB2* | 1.761378 | 0.010662 |
| *VIM* | 1.761021 | 0.002130 |
| *OAF* | 1.757908 | 0.002238 |
| *GNG11* | 1.755994 | 0.032527 |
| *DAB2* | 1.754970 | 0.002584 |
| *NR5A2* | 1.753504 | 0.005532 |
| *GCNT2* | 1.752831 | 0.000640 |
| *SERPIND1* | 1.747175 | 0.001360 |
| *CPA4* | 1.743376 | 0.008392 |
| *GP6* | 1.740278 | 0.004107 |
| *ZEB1* | 1.727942 | 0.003952 |
| *ARHGAP24* | 1.724181 | 0.003787 |
| *TRIL* | 1.723680 | 0.014877 |
| *GABRB1* | 1.718909 | 0.007139 |
| *FXYD2* | 1.716178 | 0.008968 |
| *CA12* | 1.711974 | 0.012757 |
| *SLC1A1* | 1.705720 | 0.044881 |
| *CST5* | 1.705703 | 0.018085 |
| *UNC5CL* | 1.703325 | 0.033826 |
| *SLC26A2* | 1.701462 | 0.003734 |
| *IRX4* | 1.700900 | 0.035185 |
| *STRA6* | 1.700110 | 0.002655 |
| *ROR2* | 1.696830 | 0.000455 |
| *ANO4* | 1.696258 | 0.000379 |
| *SYNC* | 1.694059 | 0.004235 |
| *B3GALT5* | 1.693801 | 0.004749 |
| *GATA3* | 1.693652 | 0.022217 |
| *LGI2* | 1.692972 | 0.041934 |
| *TMEM72* | 1.686230 | 0.000849 |
| *SLC35G1* | 1.686037 | 0.008134 |
| *PHOX2A* | 1.681287 | 0.008623 |

| | | |
|---|---|---|
| *HTR1E* | 1.676899 | 0.000377 |
| *HSPB7* | 1.671943 | 0.019011 |
| *TNNT2* | 1.671525 | 0.002642 |
| *IRX2* | 1.669588 | 0.046334 |
| *MRAP2* | 1.659529 | 0.016440 |
| *ITGA9* | 1.657913 | 0.004900 |
| *ADCY8* | 1.646068 | 0.023116 |
| *CDH13* | 1.642811 | 0.022391 |
| *GRIK3* | 1.640550 | 0.000533 |
| *SULT1B1* | 1.639036 | 0.015702 |
| *CKMT2* | 1.633567 | 0.001144 |
| *LEF1* | 1.629145 | 0.012260 |
| *APBB1IP* | 1.625831 | 0.001241 |
| *SMLR1* | 1.625366 | 0.024146 |
| *CRYBB3* | 1.623460 | 0.021281 |
| *CRYAB* | 1.622985 | 0.040302 |
| *ACTA1* | 1.615135 | 0.008079 |
| *ENOX1* | 1.612433 | 0.003838 |
| *NEXN* | 1.612368 | 0.007263 |
| *ART5* | 1.610906 | 0.008203 |
| *SOBP* | 1.610676 | 0.002787 |
| *NCAM1* | 1.610379 | 0.000866 |
| *F2RL2* | 1.610100 | 0.000912 |
| *JPH2* | 1.609423 | 0.008151 |
| *ODAM* | 1.604262 | 0.034433 |
| *PPP1R14A* | 1.603590 | 0.034326 |
| *PCDH10* | 1.597152 | 0.005040 |
| *ASPHD1* | 1.590437 | 0.004026 |
| *LSAMP* | 1.587323 | 0.007760 |
| *ANGPTL2* | 1.581766 | 0.007601 |
| *SLITRK4* | 1.579966 | 0.000515 |
| *PLA2G12B* | 1.577241 | 0.020816 |
| *GPR50* | 1.575031 | 0.000436 |
| *LMOD1* | 1.574687 | 0.002533 |
| *GRIN2A* | 1.573834 | 0.001119 |
| *IGFL2* | 1.571905 | 0.028467 |
| *PLPPR3* | 1.567840 | 0.008860 |
| *TMEM86B* | 1.567016 | 0.013081 |
| *TGFB2* | 1.561504 | 0.046679 |
| *SVEP1* | 1.560967 | 0.003064 |
| *CIDEC* | 1.559138 | 0.005503 |

| | | |
|---|---|---|
| *MPV17L* | 1.558029 | 0.001749 |
| *CBLN2* | 1.558007 | 0.001284 |
| *MSRB3* | 1.550480 | 0.002755 |
| *GSTA2* | 1.547909 | 0.015894 |
| *FAM89A* | 1.547236 | 0.015393 |
| *ST3GAL5* | 1.546650 | 0.006655 |
| *TMEM37* | 1.545770 | 0.002387 |
| *PRAP1* | 1.545296 | 0.035490 |
| *MEDAG* | 1.544425 | 0.047564 |
| *MLN* | 1.543389 | 0.013358 |
| *ANXA6* | 1.542229 | 0.019519 |
| *ERVV-2* | 1.539897 | 0.012614 |
| *SLC22A8* | 1.536774 | 0.000518 |
| *RASSF5* | 1.536760 | 0.001798 |
| *GPR37* | 1.536382 | 0.001365 |
| *C8orf88* | 1.535774 | 0.041955 |
| *MRGPRF* | 1.535746 | 0.027817 |
| *SERPINA6* | 1.535530 | 0.001804 |
| *SST* | 1.532785 | 0.013788 |
| *LRRTM4* | 1.531449 | 0.001768 |
| *GC* | 1.527058 | 0.001295 |
| *GAL3ST3* | 1.526524 | 0.007559 |
| *ARSI* | 1.519843 | 0.001764 |
| *PLCXD3* | 1.516728 | 0.001865 |
| *DKK1* | 1.516249 | 0.002554 |
| *CCND2* | 1.513780 | 0.029444 |
| *TMEM132D* | 1.513040 | 0.000723 |
| *CCDC3* | 1.510894 | 0.016466 |
| *VCAN* | 1.510883 | 0.027490 |
| *BCHE* | 1.503736 | 0.024205 |
| *SNAI1* | 1.501577 | 0.011854 |
| *MME* | 1.499770 | 0.008825 |
| *UGT2B7* | 1.498604 | 0.035247 |
| *BMP5* | 1.494263 | 0.009161 |
| *CST2* | 1.493122 | 0.014870 |
| *GGT5* | 1.491470 | 0.015580 |
| *MAB21L2* | 1.488758 | 0.026530 |
| *TMCC3* | 1.487999 | 0.001294 |
| *FAM13C* | 1.486501 | 0.002018 |
| *SMPX* | 1.483645 | 0.015497 |
| *CYP27A1* | 1.482929 | 0.002474 |

| | | |
|---|---|---|
| *NGFR* | 1.480225 | 0.004048 |
| *AFF3* | 1.478310 | 0.002390 |
| *VEPH1* | 1.477019 | 0.011578 |
| *FIGNL2* | 1.476781 | 0.007346 |
| *SOX9* | 1.474219 | 0.025428 |
| *SLC39A14* | 1.471627 | 0.003748 |
| *MN1* | 1.470074 | 0.003879 |
| *NRXN3* | 1.463286 | 0.002595 |
| *BMERB1* | 1.462994 | 0.002466 |
| *EPDR1* | 1.462623 | 0.001620 |
| *DLC1* | 1.457579 | 0.002835 |
| *CNMD* | 1.450556 | 0.026732 |
| *VIP* | 1.449980 | 0.029939 |
| *PCOLCE* | 1.447663 | 0.019370 |
| *BICC1* | 1.447600 | 0.001326 |
| *LAMB1* | 1.443973 | 0.014131 |
| *SCD* | 1.439980 | 0.003656 |
| *SNAI2* | 1.439954 | 0.015293 |
| *C8B* | 1.438702 | 0.005437 |
| *DEPDC7* | 1.438655 | 0.002042 |
| *KDR* | 1.435198 | 0.000756 |
| *RNF175* | 1.432464 | 0.011432 |
| *SLC38A4* | 1.430860 | 0.035050 |
| *ORM2* | 1.430573 | 0.006692 |
| *DAB1* | 1.428542 | 0.000844 |
| *COL4A6* | 1.426642 | 0.003521 |
| *TNFRSF19* | 1.425169 | 0.002162 |
| *HRK* | 1.421023 | 0.002131 |
| *GSTM4* | 1.419109 | 0.000898 |
| *CD3D* | 1.418398 | 0.037139 |
| *TENM2* | 1.417099 | 0.000799 |
| *TUBB8B* | 1.416827 | 0.001046 |
| *ESM1* | 1.416149 | 0.002093 |
| *EBF3* | 1.415937 | 0.002777 |
| *THBS2* | 1.412757 | 0.037397 |
| *ISL1* | 1.410521 | 0.024911 |
| *TNMD* | 1.407704 | 0.001462 |
| *TMEM178A* | 1.406760 | 0.002389 |
| *ZCCHC24* | 1.406001 | 0.006648 |
| *REC8* | 1.405521 | 0.017390 |
| *GLT8D2* | 1.403017 | 0.047838 |

| | | |
|---|---|---|
| *TENM1* | 1.399052 | 0.005447 |
| *ZFPM2* | 1.397902 | 0.000837 |
| *TCEAL7* | 1.397624 | 0.002762 |
| *AQP1* | 1.396664 | 0.002310 |
| *CYP26A1* | 1.395342 | 0.042842 |
| *C21orf62* | 1.394456 | 0.040414 |
| *EDIL3* | 1.382223 | 0.014265 |
| *STARD8* | 1.380332 | 0.000924 |
| *AJAP1* | 1.379777 | 0.001106 |
| *HMCN1* | 1.379167 | 0.006607 |
| *HSD17B2* | 1.377997 | 0.004636 |
| *HOXA3* | 1.374232 | 0.000813 |
| *DACH1* | 1.373662 | 0.002949 |
| *GLTPD2* | 1.373597 | 0.005440 |
| *CSRP2* | 1.370522 | 0.005133 |
| *CPQ* | 1.369648 | 0.013889 |
| *SLC51B* | 1.368727 | 0.004343 |
| *HCN4* | 1.367373 | 0.006281 |
| *SCARB1* | 1.366789 | 0.007276 |
| *THY1* | 1.365312 | 0.040795 |
| *SLC18A3* | 1.361222 | 0.005370 |
| *GRK3* | 1.360033 | 0.008625 |
| *LRFN1* | 1.356770 | 0.002687 |
| *NDST3* | 1.353193 | 0.005164 |
| *SLC16A10* | 1.352834 | 0.001920 |
| *NELL1* | 1.351754 | 0.008777 |
| *CCDC160* | 1.351746 | 0.024057 |
| *FJX1* | 1.350620 | 0.044612 |
| *WIF1* | 1.348229 | 0.003484 |
| *CCBE1* | 1.346778 | 0.013889 |
| *NDP* | 1.346029 | 0.014030 |
| *SLN* | 1.341861 | 0.001724 |
| *ITLN2* | 1.340144 | 0.018575 |
| *KCNIP4* | 1.338315 | 0.004112 |
| *COL4A1* | 1.337453 | 0.008139 |
| *PCYT1B* | 1.335926 | 0.003960 |
| *KIF26B* | 1.334998 | 0.006668 |
| *CRTAC1* | 1.334596 | 0.004864 |
| *GGTLC2* | 1.331536 | 0.020216 |
| *F7* | 1.329207 | 0.002147 |
| *FADS6* | 1.327434 | 0.005543 |

| | | |
|---|---|---|
| *ARSL* | 1.322927 | 0.014098 |
| *CCDC88A* | 1.319947 | 0.001397 |
| *MGARP* | 1.318127 | 0.014296 |
| *IGFBP7* | 1.314383 | 0.006204 |
| *DPYSL4* | 1.312680 | 0.005336 |
| *MMP23B* | 1.309773 | 0.017805 |
| *C9orf64* | 1.309456 | 0.003274 |
| *TRPV4* | 1.307315 | 0.023423 |
| *ENPEP* | 1.304863 | 0.002700 |
| *SERPINB9* | 1.302230 | 0.004443 |
| *PEAR1* | 1.298009 | 0.001352 |
| *SORCS3* | 1.294748 | 0.009826 |
| *TMEM190* | 1.292417 | 0.002577 |
| *ETV1* | 1.290868 | 0.002758 |
| *FGF9* | 1.289809 | 0.005111 |
| *SCUBE3* | 1.288436 | 0.035331 |
| *WNT11* | 1.287608 | 0.013312 |
| *MFAP4* | 1.285477 | 0.005363 |
| *CACHD1* | 1.283157 | 0.002487 |
| *ITGA5* | 1.282483 | 0.025046 |
| *CNR1* | 1.282374 | 0.008660 |
| *BTNL8* | 1.280573 | 0.004640 |
| *SPHK1* | 1.277174 | 0.006250 |
| *PKNOX2* | 1.270282 | 0.010115 |
| *DDIT4L* | 1.268495 | 0.011158 |
| *TRPM6* | 1.267600 | 0.001635 |
| *ANXA9* | 1.266715 | 0.038345 |
| *BDNF* | 1.266096 | 0.042257 |
| *STXBP6* | 1.264670 | 0.004667 |
| *OGDHL* | 1.259711 | 0.001553 |
| *LAMA4* | 1.258547 | 0.007160 |
| *METTL24* | 1.257498 | 0.001946 |
| *FGFR4* | 1.255439 | 0.048591 |
| *CASR* | 1.254626 | 0.030478 |
| *NPFFR2* | 1.254124 | 0.007656 |
| *SRPX* | 1.253947 | 0.003574 |
| *SHC3* | 1.252854 | 0.006695 |
| *BNC2* | 1.252284 | 0.004811 |
| *ID2* | 1.251890 | 0.001499 |
| *SULF1* | 1.250796 | 0.024674 |
| *ITGA8* | 1.248090 | 0.001082 |

| | | |
|---|---|---|
| *IL1RAPL1* | 1.246909 | 0.030676 |
| *LDLRAD3* | 1.243640 | 0.003929 |
| *GNB4* | 1.242735 | 0.015706 |
| *P2RY6* | 1.241120 | 0.002476 |
| *ENPP1* | 1.238236 | 0.003533 |
| *C1QL2* | 1.237982 | 0.003077 |
| *ALKAL1* | 1.237043 | 0.024648 |
| *SFMBT2* | 1.235819 | 0.005136 |
| *COL21A1* | 1.233505 | 0.031903 |
| *COL4A2* | 1.233436 | 0.003361 |
| *RASGRF2* | 1.233187 | 0.007701 |
| *RALYL* | 1.232924 | 0.004688 |
| *HSD17B11* | 1.231445 | 0.023952 |
| *BACH2* | 1.231303 | 0.008818 |
| *NOX1* | 1.230367 | 0.017246 |
| *FZD8* | 1.228044 | 0.001372 |
| *SLC17A1* | 1.225831 | 0.021320 |
| *MYLK* | 1.225777 | 0.002820 |
| *FBXL7* | 1.225636 | 0.002627 |
| *FLT1* | 1.222638 | 0.041628 |
| *TNFSF4* | 1.222247 | 0.004209 |
| *PLTP* | 1.221591 | 0.002654 |
| *PTGER3* | 1.220634 | 0.031494 |
| *CHST13* | 1.219168 | 0.013457 |
| *TMEM200B* | 1.218870 | 0.006916 |
| *IQGAP2* | 1.218573 | 0.003175 |
| *PECAM1* | 1.213343 | 0.004343 |
| *CYP3A7* | 1.209276 | 0.047796 |
| *VSNL1* | 1.208692 | 0.004867 |
| *RGN* | 1.204175 | 0.011920 |
| *EPB41L2* | 1.203602 | 0.003557 |
| *PIEZO2* | 1.201767 | 0.025121 |
| *NR0B2* | 1.199482 | 0.004540 |
| *GDNF* | 1.197923 | 0.001319 |
| *COX7A1* | 1.196717 | 0.005430 |
| *ARHGEF28* | 1.195593 | 0.011563 |
| *FAM155A (NALF1 new name)* | 1.190124 | 0.001865 |
| *IHH* | 1.188725 | 0.002690 |
| *UGT3A2* | 1.186862 | 0.003269 |
| *MYL9* | 1.186368 | 0.002472 |
| *LRP2* | 1.186268 | 0.028674 |

| | | |
|---|---|---|
| *EDAR* | 1.184823 | 0.004146 |
| *DPF3* | 1.183963 | 0.004533 |
| *EID3* | 1.183501 | 0.023523 |
| *PRICKLE1* | 1.180543 | 0.004918 |
| *TRIM71* | 1.180531 | 0.005364 |
| *NXN* | 1.177594 | 0.001982 |
| *FIBIN* | 1.177311 | 0.005293 |
| *ATOH8* | 1.175704 | 0.024865 |
| *LIN28B* | 1.175057 | 0.043000 |
| *PPP2R2B* | 1.173115 | 0.007980 |
| *C1S* | 1.171392 | 0.007332 |
| *DTX1* | 1.171246 | 0.026545 |
| *XCL1* | 1.170676 | 0.002575 |
| *SMOC1* | 1.167883 | 0.036270 |
| *COL9A1* | 1.162996 | 0.048066 |
| *GLT1D1* | 1.161552 | 0.002278 |
| *TFAP2A* | 1.158099 | 0.014745 |
| *KCNJ5* | 1.154681 | 0.011994 |
| *PDZD4* | 1.151712 | 0.013978 |
| *OLFML1* | 1.151123 | 0.004268 |
| *ST6GALNAC3* | 1.150670 | 0.026719 |
| *CHRNA3* | 1.150586 | 0.006712 |
| *LIMCH1* | 1.150352 | 0.012102 |
| *LOXL1* | 1.150118 | 0.034656 |
| *CYYR1* | 1.147694 | 0.020485 |
| *ETV5* | 1.147362 | 0.010929 |
| *SGCD* | 1.147263 | 0.013189 |
| *HOXA2* | 1.146642 | 0.013043 |
| *SESN3* | 1.146384 | 0.001912 |
| *GLP1R* | 1.146365 | 0.020122 |
| *TOX2* | 1.145368 | 0.039433 |
| *CLIP3* | 1.143909 | 0.002595 |
| *ECSCR* | 1.143027 | 0.030705 |
| *MYC* | 1.142383 | 0.036299 |
| *MEGF6* | 1.141584 | 0.033582 |
| *ETV4* | 1.140824 | 0.013885 |
| *MRAS* | 1.140214 | 0.004249 |
| *ME1* | 1.138460 | 0.004307 |
| *GLRB* | 1.136878 | 0.009336 |
| *CYSLTR2* | 1.136503 | 0.002012 |
| *PALM2AKAP2* | 1.135205 | 0.005131 |

| | | |
|---|---|---|
| *SUCNR1* | 1.132610 | 0.003955 |
| *ID3* | 1.132337 | 0.010157 |
| *DPYSL3* | 1.132094 | 0.006862 |
| *ABCC6* | 1.131450 | 0.048896 |
| *GFRA1* | 1.130533 | 0.002478 |
| *ZEB2* | 1.125615 | 0.022652 |
| *PLXNA4* | 1.122851 | 0.023738 |
| *ISLR2* | 1.121884 | 0.014167 |
| *POGLUT2* | 1.119533 | 0.038331 |
| *MOXD1* | 1.118759 | 0.011910 |
| *RTN4RL1* | 1.117742 | 0.028155 |
| *PARVB* | 1.116722 | 0.014153 |
| *SOX11* | 1.113092 | 0.005153 |
| *SLC5A12* | 1.111262 | 0.003362 |
| *LCTL* | 1.111148 | 0.009203 |
| *SYNPO* | 1.110628 | 0.019308 |
| *CTSL* | 1.106017 | 0.014054 |
| *CPN1* | 1.104835 | 0.016934 |
| *HLA-DRB1* | 1.103876 | 0.033399 |
| *EDEM2* | 1.102468 | 0.018654 |
| *SLC2A8* | 1.101973 | 0.002129 |
| *NR1H4* | 1.100942 | 0.003411 |
| *PCDH7* | 1.100843 | 0.009500 |
| *SMAD9* | 1.100088 | 0.028507 |
| *ATP1B2* | 1.099960 | 0.021689 |
| *CTTNBP2* | 1.099684 | 0.003717 |
| *ADRA2C* | 1.099188 | 0.015726 |
| *SEMA3E* | 1.098580 | 0.025417 |
| *C5orf63* | 1.097868 | 0.017434 |
| *SULT2A1* | 1.095227 | 0.048243 |
| *HACD1* | 1.094155 | 0.002208 |
| *XPNPEP2* | 1.093186 | 0.033049 |
| *SERPINB3* | 1.092813 | 0.012963 |
| *EMILIN2* | 1.091034 | 0.003813 |
| *VCAM1* | 1.090626 | 0.021171 |
| *ANGPT1* | 1.089598 | 0.011522 |
| *B4GAT1* | 1.089196 | 0.002086 |
| *LGI1* | 1.088752 | 0.004579 |
| *ALDH3A2* | 1.088127 | 0.036411 |
| *SHISA9* | 1.086472 | 0.001960 |
| *GPRC5B* | 1.085821 | 0.004023 |

| | | |
|---|---|---|
| *GOLIM4* | 1.084406 | 0.041515 |
| *ABCB1* | 1.084224 | 0.002758 |
| *ISL2* | 1.082835 | 0.003601 |
| *POU3F4* | 1.082003 | 0.029126 |
| *SERPINF1* | 1.081361 | 0.010921 |
| *PPM1H* | 1.080813 | 0.011795 |
| *CPVL* | 1.080067 | 0.041940 |
| *ADH6* | 1.078284 | 0.035999 |
| *TNFSF12-TNFSF13* | 1.077806 | 0.001826 |
| *TAC1* | 1.076584 | 0.049640 |
| *NAV3* | 1.073733 | 0.012292 |
| *KIF5C* | 1.073621 | 0.022514 |
| *AGAP2* | 1.070165 | 0.018630 |
| *PRSS23* | 1.069263 | 0.012426 |
| *NAT2* | 1.067648 | 0.015780 |
| *MS4A10* | 1.067634 | 0.002235 |
| *VASN* | 1.063005 | 0.024604 |
| *PDLIM3* | 1.062079 | 0.024365 |
| *CPED1* | 1.061862 | 0.009735 |
| *KLHL4* | 1.057857 | 0.006783 |
| *GYPC* | 1.056041 | 0.004561 |
| *LAMP5* | 1.054734 | 0.007795 |
| *COL17A1* | 1.053660 | 0.013676 |
| *PRRT4* | 1.050463 | 0.004624 |
| *SPARC* | 1.046156 | 0.002819 |
| *LIFR* | 1.044593 | 0.032018 |
| *KHDRBS2* | 1.044511 | 0.040357 |
| *HHEX* | 1.040459 | 0.021090 |
| *EPB41L3* | 1.040217 | 0.004919 |
| *CSDC2* | 1.035182 | 0.006312 |
| *CA3* | 1.033633 | 0.024325 |
| *CCDC152* | 1.033573 | 0.008788 |
| *EMP3* | 1.032848 | 0.024143 |
| *PROCR* | 1.032278 | 0.003321 |
| *ENC1* | 1.030041 | 0.020673 |
| *AGMAT* | 1.029926 | 0.006360 |
| *CA10* | 1.026620 | 0.009652 |
| *C2CD4C* | 1.025759 | 0.033323 |
| *ASGR1* | 1.024341 | 0.009203 |
| *SMIM1* | 1.023396 | 0.025571 |
| *ABCC2* | 1.022091 | 0.021114 |

| | | |
|---|---|---|
| *ADAM23* | 1.021584 | 0.003464 |
| *NFE2* | 1.019920 | 0.012958 |
| *PDGFC* | 1.019591 | 0.014452 |
| *WDR86* | 1.019500 | 0.006281 |
| *PREX2* | 1.018390 | 0.010148 |
| *MYO18B* | 1.018386 | 0.003189 |
| *VIT* | 1.017858 | 0.016619 |
| *GRIK1* | 1.017587 | 0.018332 |
| *KCNE4* | 1.013894 | 0.017643 |
| *GPC5* | 1.013371 | 0.013080 |
| *OLFM2* | 1.012845 | 0.002536 |
| *ATP1B1* | 1.010252 | 0.003330 |
| *FGFR1* | 1.007798 | 0.014626 |
| *MYO16* | 1.007168 | 0.009633 |
| *MEGF10* | 1.006160 | 0.014980 |
| *ARPP21* | 1.006118 | 0.015371 |
| *FKBP10* | 1.005879 | 0.003947 |
| *PCDHAC2* | 1.004188 | 0.004193 |
| *PCDH18* | 1.003608 | 0.030795 |
| *KIT* | 1.003465 | 0.044687 |
| *RAB29* | 1.003001 | 0.020488 |
| *TMEM131L* | 1.002776 | 0.004259 |
| *GABRA4* | 1.001421 | 0.002916 |
| *MASP1* | 1.000861 | 0.027190 |
| *PCDH9* | 1.000114 | 0.005019 |

**Supplementary Table 4.** Top downregulated DEGs in *FOXA2*$^{-/-}$ islets compared with WT-islets (Log2 FC < −1, $P < 0.05$).

| **Gene ID** | **Log2 FC** | ***P*-value** |
|---|---|---|
| *CPB1* | -6.545 | 0.000094 |
| *UPK1A* | -6.440 | 0.000009 |
| *MUC5AC* | -5.636 | 0.000093 |
| *FRZB* | -5.472 | 0.000627 |
| *CXCL8* | -5.110 | 0.000028 |
| *LRRN3* | -4.972 | 0.000111 |
| *INS* | -4.814 | 0.000564 |
| *TMEM265* | -4.780 | 0.003155 |
| *CXCL17* | -4.728 | 0.000015 |
| *PRSS1* | -4.690 | 0.001305 |

| | | |
|---|---|---|
| *SCGB1A1* | -4.614 | 0.000443 |
| *PKHD1L1* | -4.600 | 0.000059 |
| *UBD* | -4.498 | 0.000141 |
| *BMP3* | -4.353 | 0.000033 |
| *PPBP* | -4.327 | 0.001568 |
| *RGPD2* | -4.308 | 0.000314 |
| *UPK2* | -4.307 | 0.000533 |
| *ATP2A3* | -4.273 | 0.000063 |
| *PRSS2* | -4.223 | 0.000653 |
| *NKX6.3* | -4.164 | 0.000028 |
| *GCG* | -4.083 | 0.000356 |
| *FER1L6* | -4.067 | 0.020197 |
| *KIAA1324* | -4.040 | 0.000128 |
| *OR2I1P* | -4.011 | 0.000264 |
| *SOX21* | -3.950 | 0.002865 |
| *INSM1* | -3.919 | 0.005020 |
| *MBOAT1* | -3.882 | 0.000020 |
| *COMP* | -3.830 | 0.000453 |
| *PYY* | -3.819 | 0.002394 |
| *MMP10* | -3.812 | 0.028758 |
| *ITM2A* | -3.802 | 0.000076 |
| *HPGD* | -3.786 | 0.003154 |
| *CDH12* | -3.727 | 0.002478 |
| *HS3ST6* | -3.691 | 0.000093 |
| *MUC4* | -3.690 | 0.009579 |
| *LYPD2* | -3.686 | 0.005129 |
| *PAX9* | -3.648 | 0.003233 |
| *ZNF559* | -3.620 | 0.000025 |
| *RPRM* | -3.573 | 0.005719 |
| *MUC1* | -3.572 | 0.000084 |
| *MS4A8* | -3.490 | 0.000069 |
| *MUC20* | -3.490 | 0.000030 |
| *MSMB* | -3.478 | 0.000154 |
| *NTS* | -3.477 | 0.027544 |
| *LRAT* | -3.451 | 0.000176 |
| *CLDN18* | -3.449 | 0.000156 |
| *RGS10* | -3.426 | 0.017489 |
| *PEG3* | -3.422 | 0.000039 |
| *DEGS2* | -3.406 | 0.000031 |
| *EPHA7* | -3.405 | 0.008581 |
| *SLC6A4* | -3.360 | 0.002002 |

| | | |
|---|---|---|
| *UCN3* | -3.333 | 0.001773 |
| *CCNO* | -3.329 | 0.032135 |
| *NPW* | -3.316 | 0.004834 |
| *ANXA10* | -3.291 | 0.009944 |
| *TACR1* | -3.282 | 0.003254 |
| *SOX2* | -3.250 | 0.002448 |
| *NEUROG3* | -3.220 | 0.003294 |
| *GMNC* | -3.194 | 0.000033 |
| *AGR3* | -3.173 | 0.000973 |
| *SYK* | -3.167 | 0.000124 |
| *KLK12* | -3.157 | 0.004112 |
| *KLK11* | -3.155 | 0.000450 |
| *RASSF9* | -3.154 | 0.001075 |
| *DEPTOR* | -3.134 | 0.000139 |
| *ZBTB7C* | -3.116 | 0.001055 |
| *GDPD3* | -3.105 | 0.001694 |
| *PCDH19* | -3.100 | 0.001241 |
| *FAM3D* | -3.089 | 0.000950 |
| *ATP13A4* | -3.079 | 0.001806 |
| *LMO3* | -3.078 | 0.002070 |
| *CXCL1* | -3.048 | 0.001018 |
| *ACSL1* | -3.046 | 0.001702 |
| *MMP7* | -3.008 | 0.007840 |
| *GRHL3* | -3.006 | 0.000965 |
| *S100P* | -2.987 | 0.001245 |
| *ZNF208* | -2.981 | 0.004176 |
| *ZNF471* | -2.975 | 0.000890 |
| *ZNF667* | -2.966 | 0.000051 |
| *KLK10* | -2.965 | 0.000084 |
| *MSLN* | -2.964 | 0.000478 |
| *SGPP2* | -2.958 | 0.000072 |
| *PERCC1* | -2.941 | 0.000131 |
| *GJB7* | -2.940 | 0.000225 |
| *IGFBP1* | -2.926 | 0.012484 |
| *RARRES2* | -2.924 | 0.014587 |
| *DLL1* | -2.908 | 0.000551 |
| *NKX2-2* | -2.897 | 0.008452 |
| *GABRA1* | -2.890 | 0.011704 |
| *UGT2B15* | -2.888 | 0.045793 |
| *PAPLN* | -2.883 | 0.003091 |
| *IL1A* | -2.878 | 0.010967 |

| | | |
|---|---|---|
| *MYBPC1* | -2.877 | 0.003957 |
| *UPK3B* | -2.865 | 0.018825 |
| *EHF* | -2.841 | 0.000909 |
| *ZACN* | -2.830 | 0.027083 |
| *CNTN3* | -2.825 | 0.021109 |
| *TMSB15A* | -2.819 | 0.016865 |
| *AFF2* | -2.791 | 0.000194 |
| *FUT3* | -2.789 | 0.000167 |
| *C10orf99* | -2.782 | 0.001155 |
| *KRT7* | -2.780 | 0.002027 |
| *NTN1* | -2.767 | 0.000277 |
| *KLK1* | -2.767 | 0.000763 |
| *CXCL3* | -2.763 | 0.000079 |
| *HEPACAM2* | -2.755 | 0.000262 |
| *LCK* | -2.734 | 0.000246 |
| *MLPH* | -2.701 | 0.003326 |
| *RND1* | -2.695 | 0.000588 |
| *FA2H* | -2.678 | 0.000104 |
| *ZNF676* | -2.657 | 0.000369 |
| *ASCL1* | -2.654 | 0.025424 |
| *SAMD13* | -2.647 | 0.000092 |
| *PDE8B* | -2.646 | 0.000349 |
| *NXPH2* | -2.644 | 0.000310 |
| *RGPD1* | -2.638 | 0.000907 |
| *FAM149A* | -2.638 | 0.000065 |
| *SYBU* | -2.637 | 0.001298 |
| *ITGA4* | -2.618 | 0.027904 |
| *ATP6V1B1* | -2.598 | 0.000273 |
| *IL23A* | -2.592 | 0.006872 |
| *CRIP1* | -2.573 | 0.005462 |
| *MCIDAS* | -2.570 | 0.044623 |
| *KCNK16* | -2.564 | 0.000834 |
| *TMPRSS4* | -2.558 | 0.003784 |
| *GGT6* | -2.550 | 0.004181 |
| *NIBAN1* | -2.549 | 0.000257 |
| *GNA15* | -2.539 | 0.006887 |
| *PLCE1* | -2.535 | 0.001097 |
| *ZNF736* | -2.534 | 0.000090 |
| *UNC13C* | -2.530 | 0.001950 |
| *PCDHA6* | -2.515 | 0.000315 |
| *GFRA3* | -2.497 | 0.005359 |

| | | |
|---|---|---|
| *PTF1A* | -2.496 | 0.022354 |
| *TNFRSF11B* | -2.494 | 0.003990 |
| *PTPRT* | -2.480 | 0.004472 |
| *RNF223* | -2.474 | 0.000085 |
| *SLC30A2* | -2.472 | 0.000162 |
| *KRTAP2-3* | -2.463 | 0.020646 |
| *AR* | -2.454 | 0.000132 |
| *SUSD4* | -2.451 | 0.001329 |
| *ALDH1A3* | -2.450 | 0.000733 |
| *CAMK2B* | -2.441 | 0.000227 |
| *ATP10B* | -2.437 | 0.005857 |
| *BIK* | -2.429 | 0.008291 |
| *GUCA2B* | -2.424 | 0.003863 |
| *COL22A1* | -2.423 | 0.003374 |
| *ZNF726* | -2.414 | 0.000436 |
| *APOL4* | -2.397 | 0.000208 |
| *CELSR1* | -2.396 | 0.000290 |
| *ADGRF1* | -2.376 | 0.000223 |
| *VSIG1* | -2.366 | 0.042825 |
| *FAM83E* | -2.343 | 0.000128 |
| *TRIB3* | -2.340 | 0.001052 |
| *ZNF177* | -2.338 | 0.006792 |
| *IL20RA* | -2.321 | 0.000292 |
| *SMIM5* | -2.317 | 0.001982 |
| *UPK3BL1* | -2.316 | 0.003461 |
| *TRIM36* | -2.311 | 0.008072 |
| *MYB* | -2.299 | 0.010419 |
| *ALDH3B2* | -2.299 | 0.012061 |
| *SDCBP2* | -2.296 | 0.008590 |
| *NHS* | -2.294 | 0.034727 |
| *UGT2B17* | -2.287 | 0.021883 |
| *RNASE4* | -2.286 | 0.000478 |
| *PALMD* | -2.282 | 0.044211 |
| *PNPO* | -2.281 | 0.001244 |
| *SPTSSB* | -2.276 | 0.010386 |
| *C16orf89* | -2.273 | 0.000258 |
| *ABO* | -2.269 | 0.011387 |
| *TMPRSS13* | -2.268 | 0.001549 |
| *LPAR3* | -2.243 | 0.012714 |
| *LINGO2* | -2.237 | 0.000351 |
| *LURAP1L* | -2.228 | 0.001379 |

| | | |
|---|---|---|
| *CXCL5* | -2.226 | 0.001216 |
| *CAMK1D* | -2.225 | 0.007096 |
| *ANG* | -2.223 | 0.000668 |
| *HES6* | -2.223 | 0.022610 |
| *VSIG2* | -2.222 | 0.005441 |
| *OTX1* | -2.221 | 0.000320 |
| *SMIM22* | -2.220 | 0.000620 |
| *LHX5* | -2.216 | 0.000359 |
| *LPAR5* | -2.206 | 0.000290 |
| *NECTIN4* | -2.203 | 0.001683 |
| *GAST* | -2.192 | 0.026778 |
| *P2RX1* | -2.179 | 0.001702 |
| *RNF224* | -2.178 | 0.001098 |
| *LAMB3* | -2.176 | 0.026730 |
| *RASSF2* | -2.163 | 0.004210 |
| *LMX1B* | -2.154 | 0.013177 |
| *KRT6B* | -2.153 | 0.015350 |
| *CHRM1* | -2.141 | 0.001047 |
| *SHH* | -2.138 | 0.000222 |
| *FOLH1* | -2.130 | 0.014639 |
| *PPY* | -2.130 | 0.006051 |
| *SEMA4A* | -2.121 | 0.001269 |
| *CLDN10* | -2.111 | 0.034158 |
| *BCAS1* | -2.104 | 0.001418 |
| *RIMKLA* | -2.102 | 0.001543 |
| *EVPL* | -2.100 | 0.000685 |
| *RNF186* | -2.099 | 0.001328 |
| *SMPD3* | -2.097 | 0.000398 |
| *PCSK2* | -2.093 | 0.001923 |
| *SERPINB2* | -2.090 | 0.047196 |
| *KIF19* | -2.085 | 0.015270 |
| *MYO5B* | -2.082 | 0.000829 |
| *CPM* | -2.079 | 0.018724 |
| *S100A2* | -2.075 | 0.000153 |
| *SPACA4* | -2.073 | 0.027058 |
| *TMEM100* | -2.071 | 0.017443 |
| *LTB* | -2.068 | 0.006267 |
| *SLC38A3* | -2.068 | 0.000314 |
| *IRF5* | -2.066 | 0.000324 |
| *IL12A* | -2.061 | 0.005140 |
| *DPY19L2* | -2.051 | 0.006344 |

| | | |
|---|---|---|
| *ASTL* | -2.049 | 0.000373 |
| *SEMA3A* | -2.042 | 0.001207 |
| *EFEMP1* | -2.042 | 0.001534 |
| *RFX6* | -2.042 | 0.001358 |
| *TFF1* | -2.040 | 0.029253 |
| *ZNF835* | -2.037 | 0.000277 |
| *BAIAP3* | -2.033 | 0.008062 |
| *AOC1* | -2.030 | 0.003640 |
| *FAM167A* | -2.029 | 0.015739 |
| *C2orf72* | -2.028 | 0.000444 |
| *TAAR1* | -2.027 | 0.002781 |
| *PLCL2* | -2.024 | 0.000212 |
| *UCP2* | -2.013 | 0.006024 |
| *RIPPLY3* | -2.011 | 0.004640 |
| *SEMA3F* | -2.008 | 0.001696 |
| *NKX6.1* | -2.008 | 0.020767 |
| *NOL4* | -2.007 | 0.000217 |
| *SLC25A12* | -2.001 | 0.001692 |
| *MLXIPL* | -1.997 | 0.001945 |
| *GNA14* | -1.996 | 0.024093 |
| *SPINK1* | -1.995 | 0.002861 |
| *KLF5* | -1.994 | 0.000617 |
| *CACNG4* | -1.994 | 0.000419 |
| *LRRC61* | -1.977 | 0.001890 |
| *OVOL2* | -1.976 | 0.035332 |
| *TMCC2* | -1.962 | 0.020002 |
| *CNTNAP3* | -1.961 | 0.000230 |
| *PTGS2* | -1.960 | 0.003701 |
| *LRRC10B* | -1.959 | 0.041431 |
| *ZXDA* | -1.957 | 0.000187 |
| *VWA5B2* | -1.952 | 0.001372 |
| *THRB* | -1.950 | 0.009821 |
| *GDF15* | -1.947 | 0.000501 |
| *FUT1* | -1.945 | 0.000782 |
| *SEMA3C* | -1.938 | 0.011795 |
| *CYP4F12* | -1.938 | 0.022220 |
| *CLIC5* | -1.935 | 0.017618 |
| *PLA2G4B* | -1.932 | 0.015796 |
| *SLC34A3* | -1.930 | 0.001550 |
| *ARHGAP30* | -1.929 | 0.000384 |
| *SPIB* | -1.928 | 0.003433 |

| | | |
|---|---|---|
| *MYLPF* | -1.927 | 0.047784 |
| *C6orf141* | -1.921 | 0.014020 |
| *PID1* | -1.920 | 0.002186 |
| *GPBAR1* | -1.914 | 0.000765 |
| *EMID1* | -1.914 | 0.010140 |
| *TNF* | -1.913 | 0.002522 |
| *LRRC4* | -1.913 | 0.000229 |
| *ACSS1* | -1.908 | 0.000465 |
| *HIF3A* | -1.902 | 0.004680 |
| *HSPE1-MOB4* | -1.902 | 0.030515 |
| *PLAUR* | -1.899 | 0.003615 |
| *CDH9* | -1.896 | 0.014012 |
| *TRPM4* | -1.895 | 0.000285 |
| *CDH7* | -1.895 | 0.025479 |
| *KCNN1* | -1.892 | 0.006227 |
| *LCN2* | -1.892 | 0.000541 |
| *DLL4* | -1.891 | 0.001372 |
| *FRY* | -1.890 | 0.002506 |
| *COL14A1* | -1.889 | 0.019458 |
| *FERMT1* | -1.885 | 0.027903 |
| *PCDHGA10* | -1.881 | 0.002355 |
| *GABRB2* | -1.881 | 0.023961 |
| *PLCH2* | -1.880 | 0.000802 |
| *DMBT1* | -1.877 | 0.000429 |
| *PANX2* | -1.874 | 0.002072 |
| *TPBG* | -1.869 | 0.000695 |
| *MUC5B* | -1.867 | 0.017076 |
| *CCL20* | -1.861 | 0.020581 |
| *SNTB1* | -1.852 | 0.001054 |
| *TMEM156* | -1.850 | 0.005598 |
| *PRODH* | -1.847 | 0.000411 |
| *MISP* | -1.844 | 0.001439 |
| *ALOX15* | -1.838 | 0.028500 |
| *KIAA0319* | -1.835 | 0.005467 |
| *SPRR2A* | -1.834 | 0.001655 |
| *BMPR1B* | -1.833 | 0.000304 |
| *EBF4* | -1.833 | 0.002013 |
| *KLF4* | -1.826 | 0.007130 |
| *IRAK2* | -1.822 | 0.023421 |
| *UPP1* | -1.821 | 0.040542 |
| *ATG16L2* | -1.820 | 0.013471 |

| | | |
|---|---|---|
| *IQANK1* | -1.819 | 0.004081 |
| *UPK3BL2* | -1.819 | 0.000265 |
| *CDHR1* | -1.818 | 0.002940 |
| *PHACTR3* | -1.818 | 0.010215 |
| *PCDH1* | -1.813 | 0.027435 |
| *EXPH5* | -1.812 | 0.001166 |
| *COL6A2* | -1.807 | 0.006240 |
| *CAPS* | -1.803 | 0.000588 |
| *ABCG1* | -1.797 | 0.001020 |
| *NR3C2* | -1.796 | 0.005904 |
| *SIDT1* | -1.795 | 0.000771 |
| *ARAP3* | -1.794 | 0.018435 |
| *SCNN1A* | -1.794 | 0.028341 |
| *DACH2* | -1.792 | 0.001283 |
| *DSTN* | -1.790 | 0.019380 |
| *FOXQ1* | -1.790 | 0.021329 |
| *NEUROD1* | -1.789 | 0.006574 |
| *CXCL2* | -1.786 | 0.005516 |
| *SEMA7A* | -1.783 | 0.030450 |
| *MT1G* | -1.780 | 0.049183 |
| *NPY1R* | -1.775 | 0.033571 |
| *JUNB* | -1.772 | 0.001907 |
| *RELB* | -1.770 | 0.003850 |
| *KNDC1* | -1.768 | 0.001944 |
| *TNFRSF14* | -1.759 | 0.003019 |
| *RAPGEFL1* | -1.758 | 0.000874 |
| *THBD* | -1.757 | 0.000390 |
| *IL32* | -1.756 | 0.007821 |
| *NRXN1* | -1.755 | 0.001037 |
| *DOCK3* | -1.750 | 0.010172 |
| *VGLL1* | -1.750 | 0.018871 |
| *ADGRG2* | -1.749 | 0.019929 |
| *LTBP4* | -1.749 | 0.011985 |
| *ICAM4* | -1.748 | 0.046341 |
| *DNAJC12* | -1.748 | 0.006754 |
| *PRKCG* | -1.740 | 0.013747 |
| *APOH* | -1.739 | 0.036507 |
| *NXPH4* | -1.737 | 0.022630 |
| *PAQR6* | -1.735 | 0.010609 |
| *GRB14* | -1.731 | 0.004872 |
| *UGDH* | -1.729 | 0.001938 |

| | | |
|---|---|---|
| *PLAAT4* | -1.728 | 0.007241 |
| *GALNT18* | -1.728 | 0.008630 |
| *TGM1* | -1.726 | 0.001106 |
| *CRACR2B* | -1.721 | 0.000633 |
| *SCPEP1* | -1.721 | 0.015748 |
| *GPM6B* | -1.721 | 0.006094 |
| *TCTEX1D4* | -1.718 | 0.000696 |
| *SLFN13* | -1.718 | 0.007944 |
| *FAM174B* | -1.716 | 0.000811 |
| *POMC* | -1.715 | 0.019984 |
| *SYNE2* | -1.714 | 0.002197 |
| *TNNT1* | -1.711 | 0.018566 |
| *KIF21B* | -1.708 | 0.021579 |
| *KLHL35* | -1.704 | 0.001337 |
| *USP6* | -1.702 | 0.005130 |
| *F3* | -1.700 | 0.030851 |
| *C19orf33* | -1.699 | 0.001135 |
| *WNT4* | -1.690 | 0.024940 |
| *RAMP1* | -1.689 | 0.005853 |
| *SMAGP* | -1.688 | 0.006273 |
| *MAP3K5* | -1.688 | 0.002158 |
| *ITGA2B* | -1.687 | 0.013580 |
| *TNFSF9* | -1.686 | 0.001146 |
| *ZBED6CL* | -1.681 | 0.002556 |
| *SLC22A9* | -1.675 | 0.004789 |
| *CGB8* | -1.673 | 0.012099 |
| *TSPAN12* | -1.671 | 0.015625 |
| *C9orf152* | -1.671 | 0.002636 |
| *GALR2* | -1.670 | 0.043973 |
| *SLC16A11* | -1.668 | 0.002473 |
| *PDLIM4* | -1.666 | 0.038097 |
| *METTL7A* | -1.665 | 0.003765 |
| *SLC30A8* | -1.663 | 0.002418 |
| *ACP7* | -1.660 | 0.000585 |
| *KLHL1* | -1.652 | 0.002767 |
| *STYK1* | -1.651 | 0.001026 |
| *SGSM1* | -1.650 | 0.000517 |
| *ZNF257* | -1.650 | 0.000362 |
| *PCDHB5* | -1.647 | 0.002952 |
| *TAFA3* | -1.643 | 0.023058 |
| *EPB41L4A* | -1.642 | 0.013227 |

| | | |
|---|---|---|
| *PLEK2* | -1.641 | 0.016646 |
| *ADAM8* | -1.639 | 0.004164 |
| *PLAU* | -1.637 | 0.008937 |
| *GPSM3* | -1.637 | 0.002378 |
| *ZNF229* | -1.637 | 0.000381 |
| *TMEM79* | -1.636 | 0.000400 |
| *ATF3* | -1.635 | 0.000555 |
| *CELF3* | -1.633 | 0.005517 |
| *TACC2* | -1.629 | 0.002510 |
| *SLC15A2* | -1.624 | 0.013754 |
| *DOCK5* | -1.615 | 0.031724 |
| *ARX* | -1.613 | 0.045159 |
| *SLFN5* | -1.612 | 0.004258 |
| *ZC3H12A* | -1.609 | 0.000533 |
| *TEN1-CDK3* | -1.607 | 0.003615 |
| *PER2* | -1.600 | 0.001656 |
| *AOX1* | -1.595 | 0.019656 |
| *EPN3* | -1.595 | 0.000573 |
| *ZNF69* | -1.590 | 0.003002 |
| *TGFA* | -1.590 | 0.000986 |
| *LFNG* | -1.589 | 0.002395 |
| *ZBTB42* | -1.588 | 0.001452 |
| *EPHA10* | -1.587 | 0.004934 |
| *TREX1* | -1.584 | 0.017851 |
| *ZG16B* | -1.577 | 0.000451 |
| *TMEM176A* | -1.575 | 0.003318 |
| *TRIM47* | -1.571 | 0.006807 |
| *ENTPD2* | -1.571 | 0.006145 |
| *SLC9A2* | -1.570 | 0.004095 |
| *ADH1A* | -1.569 | 0.003276 |
| *REEP1* | -1.568 | 0.004048 |
| *TNNC1* | -1.567 | 0.033675 |
| *IGFL1* | -1.563 | 0.001179 |
| *TRIM38* | -1.561 | 0.002410 |
| *TPH1* | -1.560 | 0.001016 |
| *C20orf204* | -1.560 | 0.030023 |
| *ARRDC3* | -1.556 | 0.002320 |
| *KRT81* | -1.556 | 0.001395 |
| *RAB3D* | -1.553 | 0.001045 |
| *IKZF2* | -1.551 | 0.002351 |
| *FXYD4* | -1.550 | 0.001318 |

| | | |
|---|---|---|
| *IFI44* | -1.550 | 0.007901 |
| *HPSE* | -1.550 | 0.001533 |
| *OAS3* | -1.549 | 0.019931 |
| *FAM160A1* | -1.548 | 0.001030 |
| *SYT5* | -1.546 | 0.016911 |
| *ERO1B* | -1.544 | 0.003526 |
| *CAPN5* | -1.544 | 0.001028 |
| *GJA1* | -1.540 | 0.001632 |
| *ACTL10* | -1.539 | 0.009968 |
| *TFF3* | -1.538 | 0.001947 |
| *TNFRSF11A* | -1.538 | 0.003172 |
| *STX19* | -1.535 | 0.000482 |
| *OXGR1* | -1.534 | 0.025014 |
| *GALNT12* | -1.532 | 0.001709 |
| *POF1B* | -1.531 | 0.022017 |
| *FBP1* | -1.531 | 0.008191 |
| *CYP2C18* | -1.529 | 0.018006 |
| *OLFM4* | -1.526 | 0.017430 |
| *B4GALNT3* | -1.526 | 0.008377 |
| *TMCO4* | -1.524 | 0.003658 |
| *CDKN1B* | -1.524 | 0.014230 |
| *KLK13* | -1.521 | 0.003769 |
| *ATP13A5* | -1.519 | 0.002778 |
| *GCNT1* | -1.519 | 0.015917 |
| *RASSF6* | -1.517 | 0.033911 |
| *ZFP36L1* | -1.515 | 0.001275 |
| *TRIM45* | -1.514 | 0.003811 |
| *WNK2* | -1.514 | 0.000870 |
| *E2F7* | -1.514 | 0.002304 |
| *GRHL1* | -1.510 | 0.000923 |
| *GALNT14* | -1.506 | 0.002348 |
| *ADGRG5* | -1.503 | 0.004548 |
| *AMOT* | -1.503 | 0.008357 |
| *ANGPT2* | -1.501 | 0.009879 |
| *UBA7* | -1.500 | 0.009183 |
| *AREG* | -1.500 | 0.012832 |
| *RUNX2* | -1.499 | 0.010266 |
| *SCG3* | -1.498 | 0.028335 |
| *MFNG* | -1.496 | 0.003282 |
| *HSH2D* | -1.493 | 0.004141 |
| *RNF183* | -1.488 | 0.023980 |

| | | |
|---|---|---|
| *C1orf127* | -1.486 | 0.000841 |
| *NEXMIF* | -1.485 | 0.035935 |
| *C20orf194* | -1.483 | 0.002156 |
| *TBC1D8* | -1.483 | 0.000997 |
| *CHD3* | -1.478 | 0.001971 |
| *TNFAIP3* | -1.477 | 0.004433 |
| *ZFP36* | -1.476 | 0.002616 |
| *TMEM255A* | -1.476 | 0.038011 |
| *SLC10A5* | -1.474 | 0.005407 |
| *C15orf39* | -1.474 | 0.000970 |
| *GIPR* | -1.471 | 0.046349 |
| *RARB* | -1.471 | 0.024582 |
| *KCNH6* | -1.471 | 0.012661 |
| *OXTR* | -1.470 | 0.005707 |
| *SPDEF* | -1.469 | 0.003887 |
| *S100A6* | -1.468 | 0.001581 |
| *HS6ST2* | -1.463 | 0.004457 |
| *CDO1* | -1.462 | 0.003938 |
| *VNN2* | -1.462 | 0.013233 |
| *FOXN1* | -1.461 | 0.003661 |
| *SLITRK6* | -1.461 | 0.009643 |
| *USP41* | -1.460 | 0.025163 |
| *PTPRN2* | -1.458 | 0.044138 |
| *EGLN3* | -1.456 | 0.021643 |
| *CCN3* | -1.455 | 0.014954 |
| *ADCY7* | -1.455 | 0.035309 |
| *PAX4* | -1.455 | 0.046518 |
| *FSTL5* | -1.453 | 0.015670 |
| *SULT1A3* | -1.453 | 0.007054 |
| *GAB2* | -1.452 | 0.004117 |
| *LBH* | -1.451 | 0.003241 |
| *TRIM16* | -1.449 | 0.001965 |
| *ADCY1* | -1.446 | 0.002243 |
| *IRF1* | -1.442 | 0.001858 |
| *AMIGO2* | -1.441 | 0.001760 |
| *JPH1* | -1.439 | 0.036780 |
| *ZNF572* | -1.439 | 0.002002 |
| *MYOM3* | -1.438 | 0.015313 |
| *CDKN1A* | -1.438 | 0.002046 |
| *TMEM176B* | -1.437 | 0.007239 |
| *PRSS16* | -1.437 | 0.009003 |

| | | |
|---|---|---|
| *ABCA3* | -1.437 | 0.004396 |
| *ZNF528* | -1.437 | 0.035965 |
| *CDCP1* | -1.435 | 0.043382 |
| *ZNF860* | -1.432 | 0.017763 |
| *VWA2* | -1.432 | 0.004627 |
| *MTMR7* | -1.431 | 0.000798 |
| *C3orf14* | -1.431 | 0.046570 |
| *GBP6* | -1.430 | 0.034570 |
| *STAC* | -1.426 | 0.002697 |
| *CKMT1A* | -1.425 | 0.001904 |
| *KCNS3* | -1.424 | 0.001009 |
| *S100A4* | -1.419 | 0.018572 |
| *FAM167B* | -1.418 | 0.023868 |
| *SLC7A2* | -1.418 | 0.002289 |
| *B3GALT4* | -1.418 | 0.000975 |
| *PRR15L* | -1.417 | 0.004639 |
| *SCGN* | -1.417 | 0.002077 |
| *SFMBT1* | -1.415 | 0.001304 |
| *LGR4* | -1.414 | 0.002253 |
| *SLC52A3* | -1.413 | 0.003278 |
| *SUSD6* | -1.413 | 0.010001 |
| *MYH14* | -1.412 | 0.002230 |
| *IQSEC2* | -1.409 | 0.024659 |
| *IFIT3* | -1.404 | 0.005309 |
| *ANKRD22* | -1.401 | 0.007175 |
| *CRYBG1* | -1.400 | 0.002074 |
| *FGD6* | -1.399 | 0.044040 |
| *SPIRE2* | -1.398 | 0.029198 |
| *PDX1* | -1.392 | 0.004558 |
| *SPPL2B* | -1.391 | 0.003989 |
| *TTC9* | -1.389 | 0.006876 |
| *RIBC2* | -1.389 | 0.017886 |
| *PAQR4* | -1.388 | 0.040263 |
| *GJB3* | -1.388 | 0.012975 |
| *SIT1* | -1.387 | 0.023822 |
| *BFSP1* | -1.382 | 0.041800 |
| *NBL1* | -1.382 | 0.010305 |
| *CADM2* | -1.382 | 0.001277 |
| *AHNAK2* | -1.380 | 0.045924 |
| *GCGR* | -1.380 | 0.013826 |
| *BCL2L14* | -1.378 | 0.006057 |

| | | |
|---|---|---|
| *SLAMF7* | -1.377 | 0.019499 |
| *CD9* | -1.376 | 0.005930 |
| *MCTS2P* | -1.374 | 0.009316 |
| *IKBKE* | -1.372 | 0.011074 |
| *APOL1* | -1.372 | 0.044488 |
| *C21orf58* | -1.371 | 0.000787 |
| *MEOX1* | -1.367 | 0.000992 |
| *CHGA* | -1.365 | 0.026737 |
| *GCOM1* | -1.364 | 0.016783 |
| *SLC37A1* | -1.363 | 0.010094 |
| *ARRDC2* | -1.360 | 0.001894 |
| *HAUS4* | -1.360 | 0.005784 |
| *RUNX1* | -1.357 | 0.001564 |
| *TFCP2L1* | -1.356 | 0.003096 |
| *KCNH8* | -1.355 | 0.023042 |
| *ESRP2* | -1.353 | 0.002671 |
| *TACSTD2* | -1.350 | 0.003080 |
| *MYEOV* | -1.350 | 0.040870 |
| *TXNIP* | -1.350 | 0.015325 |
| *LRP11* | -1.350 | 0.029113 |
| *DOCK11* | -1.349 | 0.001677 |
| *NR4A2* | -1.345 | 0.006674 |
| *VPS37B* | -1.345 | 0.017607 |
| *ACPP* | -1.345 | 0.023512 |
| *ABCC8* | -1.342 | 0.004895 |
| *STAP2* | -1.341 | 0.000924 |
| *MAST4* | -1.340 | 0.019528 |
| *KCNK13* | -1.339 | 0.033592 |
| *CCDC187* | -1.338 | 0.023572 |
| *GPCPD1* | -1.337 | 0.002040 |
| *ZDHHC14* | -1.335 | 0.031918 |
| *KISS1* | -1.335 | 0.003537 |
| *MYZAP* | -1.334 | 0.012427 |
| *NR2F1* | -1.334 | 0.017207 |
| *MYO10* | -1.334 | 0.001229 |
| *AQP3* | -1.333 | 0.003207 |
| *ST8SIA2* | -1.333 | 0.007189 |
| *LTK* | -1.332 | 0.018947 |
| *FAM43A* | -1.328 | 0.016143 |
| *N4BP3* | -1.328 | 0.001296 |
| *NLRP2* | -1.328 | 0.001010 |

| | | |
|---|---|---|
| *XK* | -1.326 | 0.001311 |
| *CKMT1B* | -1.325 | 0.009791 |
| *CHST15* | -1.325 | 0.005334 |
| *SGSM2* | -1.324 | 0.000864 |
| *IER5* | -1.322 | 0.014909 |
| *PTPN13* | -1.320 | 0.002214 |
| *BLNK* | -1.319 | 0.008030 |
| *CPT1A* | -1.317 | 0.008920 |
| *FFAR1* | -1.316 | 0.000939 |
| *MGME1* | -1.312 | 0.001774 |
| *SLC16A5* | -1.311 | 0.004397 |
| *CLDN4* | -1.310 | 0.001050 |
| *SLC4A11* | -1.307 | 0.033926 |
| *BSPRY* | -1.307 | 0.001753 |
| *ADCY2* | -1.306 | 0.007542 |
| *CLCNKB* | -1.303 | 0.014630 |
| *ARNTL2* | -1.303 | 0.001964 |
| *TMTC2* | -1.302 | 0.014278 |
| *TRIM22* | -1.302 | 0.003237 |
| *PCDHB8* | -1.301 | 0.008582 |
| *KISS1R* | -1.300 | 0.024383 |
| *ENO2* | -1.299 | 0.031085 |
| *ALG1L* | -1.298 | 0.042150 |
| *CBLC* | -1.296 | 0.003577 |
| *CASZ1* | -1.296 | 0.007891 |
| *SYT7* | -1.296 | 0.006145 |
| *HID1* | -1.292 | 0.004379 |
| *LYPD6B* | -1.292 | 0.004131 |
| *CPLX1* | -1.291 | 0.002596 |
| *C15orf62* | -1.291 | 0.003111 |
| *C1orf116* | -1.290 | 0.014497 |
| *CCDC154* | -1.289 | 0.001366 |
| *PSCA* | -1.288 | 0.003341 |
| *AQP5* | -1.286 | 0.046170 |
| *SLCO3A1* | -1.286 | 0.020220 |
| *RECQL5* | -1.285 | 0.024966 |
| *MYCL* | -1.285 | 0.025711 |
| *HES1* | -1.284 | 0.037253 |
| *TLR5* | -1.284 | 0.036120 |
| *PHYHD1* | -1.283 | 0.007589 |
| *IL17RE* | -1.281 | 0.001809 |

| | | |
|---|---|---|
| *SPRR2D* | -1.280 | 0.003418 |
| *PLSCR1* | -1.278 | 0.038482 |
| *CYP2D6* | -1.278 | 0.028252 |
| *TASP1* | -1.277 | 0.004311 |
| *CHGB* | -1.277 | 0.024924 |
| *NUDT19* | -1.277 | 0.038553 |
| *ANGPTL4* | -1.277 | 0.026593 |
| *ZFP28* | -1.276 | 0.001575 |
| *MTUS2* | -1.275 | 0.001711 |
| *MEIS3* | -1.275 | 0.037301 |
| *TM4SF1* | -1.274 | 0.001406 |
| *QSOX1* | -1.274 | 0.029289 |
| *PITX1* | -1.272 | 0.001694 |
| *ENTPD8* | -1.272 | 0.044908 |
| *A4GALT* | -1.271 | 0.006219 |
| *PLAAT5* | -1.269 | 0.019926 |
| *PTAFR* | -1.268 | 0.031502 |
| *RBCK1* | -1.266 | 0.003171 |
| *MPP7* | -1.266 | 0.009161 |
| *CACNA1A* | -1.261 | 0.004077 |
| *JMJD7-PLA2G4B* | -1.261 | 0.005453 |
| *EPHX3* | -1.261 | 0.012838 |
| *MCF2L* | -1.260 | 0.005826 |
| *DUSP10* | -1.258 | 0.005786 |
| *TMEM121B* | -1.257 | 0.005964 |
| *TM6SF1* | -1.257 | 0.047085 |
| *MSH5* | -1.256 | 0.011934 |
| *CBFA2T3* | -1.255 | 0.006803 |
| *OVOL1* | -1.255 | 0.009826 |
| *NOTCH2NLC* | -1.255 | 0.001353 |
| *GPHA2* | -1.255 | 0.038673 |
| *CDHR3* | -1.254 | 0.004311 |
| *FAM102B* | -1.251 | 0.011376 |
| *ARSJ* | -1.249 | 0.021586 |
| *MOCOS* | -1.247 | 0.001293 |
| *CBLN3* | -1.247 | 0.003203 |
| *NOTCH2NLB* | -1.246 | 0.019310 |
| *TAP1* | -1.243 | 0.022793 |
| *OTULINL* | -1.240 | 0.002993 |
| *TFAP2E* | -1.240 | 0.003936 |
| *PAIP2B* | -1.240 | 0.005252 |

| | | |
|---|---|---|
| *ELF3* | -1.240 | 0.001102 |
| *FAM110A* | -1.240 | 0.001217 |
| *LMO7* | -1.239 | 0.008362 |
| *SLC41A2* | -1.238 | 0.023930 |
| *PCED1A* | -1.238 | 0.007308 |
| *ZNF559-ZNF177* | -1.236 | 0.001133 |
| *IGSF11* | -1.236 | 0.004428 |
| *DGKD* | -1.235 | 0.003855 |
| *TCF19* | -1.235 | 0.012547 |
| *NR3C1* | -1.233 | 0.001518 |
| *PLEKHG4B* | -1.233 | 0.001188 |
| *TPD52* | -1.231 | 0.003784 |
| *ITPKA* | -1.231 | 0.011748 |
| *CRYBA2* | -1.231 | 0.019914 |
| *BHLHE40* | -1.228 | 0.047768 |
| *TK1* | -1.228 | 0.045019 |
| *ARHGAP32* | -1.228 | 0.008827 |
| *PPP1R13L* | -1.228 | 0.010462 |
| *TRIM14* | -1.225 | 0.014026 |
| *TMEM163* | -1.223 | 0.003059 |
| *EBAG9* | -1.223 | 0.005029 |
| *PSD4* | -1.222 | 0.006607 |
| *LYPD6* | -1.221 | 0.005875 |
| *IFITM1* | -1.220 | 0.015091 |
| *MACC1* | -1.218 | 0.018311 |
| *SALL4* | -1.217 | 0.003436 |
| *ZNF844* | -1.217 | 0.001193 |
| *SH3RF1* | -1.217 | 0.002883 |
| *ARRB1* | -1.216 | 0.001201 |
| *GSDMD* | -1.214 | 0.005216 |
| *TMEM54* | -1.214 | 0.001855 |
| *USP54* | -1.214 | 0.010158 |
| *LRRC26* | -1.214 | 0.042423 |
| *PLA2G4F* | -1.213 | 0.003122 |
| *LTBP2* | -1.212 | 0.006728 |
| *ACE* | -1.212 | 0.009953 |
| *MISP3* | -1.211 | 0.003912 |
| *ETV7* | -1.211 | 0.024882 |
| *DOCK6* | -1.210 | 0.007924 |
| *TRIM25* | -1.210 | 0.004433 |
| *ISG20* | -1.209 | 0.038940 |

| | | |
|---|---|---|
| *SMIM14* | -1.209 | 0.007232 |
| *PSMB8* | -1.206 | 0.004600 |
| *CCDC188* | -1.205 | 0.010635 |
| *SCN3A* | -1.203 | 0.001632 |
| *IQGAP3* | -1.202 | 0.016648 |
| *H2BC21* | -1.202 | 0.006356 |
| *DENND11* | -1.201 | 0.013209 |
| *CHRNB1* | -1.200 | 0.001406 |
| *ZNF714* | -1.199 | 0.001774 |
| *DEPP1* | -1.195 | 0.011827 |
| *FAM83B* | -1.192 | 0.001950 |
| *VANGL1* | -1.191 | 0.009471 |
| *CDKN2C* | -1.191 | 0.041811 |
| *ARHGAP6* | -1.189 | 0.013606 |
| *PLEKHM3* | -1.188 | 0.006671 |
| *BRINP2* | -1.187 | 0.004569 |
| *P2RY1* | -1.186 | 0.002905 |
| *DEFB4A* | -1.185 | 0.030601 |
| *SP100* | -1.185 | 0.003582 |
| *MSH5-SAPCD1* | -1.183 | 0.003686 |
| *CACNG6* | -1.183 | 0.007517 |
| *IRF6* | -1.183 | 0.002395 |
| *TRIM34* | -1.182 | 0.027458 |
| *SQOR* | -1.181 | 0.015999 |
| *TRIM4* | -1.181 | 0.004677 |
| *MAGI2* | -1.180 | 0.002286 |
| *MAFF* | -1.179 | 0.020888 |
| *ANO8* | -1.178 | 0.015659 |
| *ZBED2* | -1.176 | 0.001721 |
| *NFKB2* | -1.176 | 0.021729 |
| *FOSL2* | -1.176 | 0.003779 |
| *H2BU1* | -1.172 | 0.040346 |
| *CCDC6* | -1.168 | 0.011111 |
| *FAM50B* | -1.167 | 0.001700 |
| *STN1* | -1.167 | 0.001454 |
| *SMOX* | -1.167 | 0.010008 |
| *RGS2* | -1.167 | 0.031553 |
| *EFS* | -1.166 | 0.004532 |
| *NFKBIA* | -1.166 | 0.005398 |
| *TENT5C* | -1.163 | 0.039488 |
| *PIPOX* | -1.162 | 0.003542 |

| | | |
|---|---|---|
| *TSPAN5* | -1.162 | 0.006943 |
| *INAVA* | -1.160 | 0.005666 |
| *ABLIM3* | -1.160 | 0.033103 |
| *PPP1R3E* | -1.158 | 0.003770 |
| *CDCA8* | -1.158 | 0.014445 |
| *NAP1L5* | -1.158 | 0.045184 |
| *DCLK2* | -1.158 | 0.037648 |
| *DDRGK1* | -1.158 | 0.014803 |
| *MUC16* | -1.156 | 0.047306 |
| *AGAP9* | -1.156 | 0.040016 |
| *NFATC4* | -1.155 | 0.009510 |
| *TAS1R3* | -1.155 | 0.004181 |
| *CAB39L* | -1.153 | 0.030330 |
| *NET1* | -1.153 | 0.011302 |
| *CELF6* | -1.151 | 0.024046 |
| *EPHB6* | -1.150 | 0.010863 |
| *ESPL1* | -1.149 | 0.001475 |
| *OSBPL7* | -1.148 | 0.040400 |
| *CENPB* | -1.148 | 0.031031 |
| *PKIA* | -1.147 | 0.003623 |
| *TNP1* | -1.146 | 0.025200 |
| *TMTC1* | -1.146 | 0.021656 |
| *RASA4B* | -1.144 | 0.016447 |
| *HERC6* | -1.143 | 0.002236 |
| *BCL2L11* | -1.143 | 0.004175 |
| *CDC7* | -1.142 | 0.006329 |
| *PYROXD2* | -1.141 | 0.021991 |
| *KCNK1* | -1.140 | 0.003410 |
| *TIPARP* | -1.140 | 0.003242 |
| *MANEAL* | -1.140 | 0.007787 |
| *PRSS50* | -1.139 | 0.031454 |
| *MXD1* | -1.139 | 0.025131 |
| *ZNF677* | -1.138 | 0.004461 |
| *PRDM5* | -1.138 | 0.003082 |
| *TSPOAP1* | -1.138 | 0.012591 |
| *CCDC51* | -1.137 | 0.014794 |
| *ERAP2* | -1.137 | 0.022797 |
| *CD36* | -1.136 | 0.024288 |
| *PGBD5* | -1.135 | 0.001688 |
| *ZNF589* | -1.135 | 0.007098 |
| *CXXC4* | -1.135 | 0.005809 |

| | | |
|---|---|---|
| *PPFIA3* | -1.135 | 0.003804 |
| *KCNH3* | -1.135 | 0.003162 |
| *TNFSF13* | -1.134 | 0.007644 |
| *STX1A* | -1.133 | 0.005714 |
| *CYP2J2* | -1.133 | 0.001703 |
| *ABHD17C* | -1.133 | 0.007580 |
| *DENND6B* | -1.133 | 0.008980 |
| *NEURL1* | -1.133 | 0.001661 |
| *TP53I11* | -1.132 | 0.002984 |
| *UBALD2* | -1.131 | 0.001977 |
| *TENT5A* | -1.131 | 0.039769 |
| *YPEL3* | -1.127 | 0.023515 |
| *H1-0* | -1.127 | 0.007382 |
| *FAM111B* | -1.126 | 0.004474 |
| *FYCO1* | -1.126 | 0.012252 |
| *STK35* | -1.125 | 0.004519 |
| *NAPB* | -1.125 | 0.005449 |
| *ZNF747* | -1.123 | 0.003309 |
| *TMEM238* | -1.123 | 0.030335 |
| *TSPAN2* | -1.122 | 0.019874 |
| *MTURN* | -1.122 | 0.009915 |
| *NOD1* | -1.121 | 0.002754 |
| *PARP12* | -1.121 | 0.004177 |
| *SFR1* | -1.121 | 0.003083 |
| *SAT1* | -1.119 | 0.004349 |
| *ZIK1* | -1.117 | 0.012928 |
| *PRRG3* | -1.116 | 0.003458 |
| *FRMPD3* | -1.113 | 0.002883 |
| *CFAP70* | -1.112 | 0.020633 |
| *AP5S1* | -1.111 | 0.002061 |
| *PCLO* | -1.111 | 0.032656 |
| *TMPRSS3* | -1.110 | 0.048472 |
| *BGLAP* | -1.110 | 0.011874 |
| *CA8* | -1.110 | 0.048710 |
| *MUC15* | -1.110 | 0.041553 |
| *NXT2* | -1.110 | 0.008700 |
| *FKBP1A* | -1.107 | 0.003741 |
| *ST18* | -1.107 | 0.014267 |
| *HLA-DQB1* | -1.107 | 0.019533 |
| *DYNLRB2* | -1.105 | 0.042977 |
| *BCO1* | -1.105 | 0.014861 |

| | | |
|---|---|---|
| *SAPCD2* | -1.104 | 0.001975 |
| *CSNK2A3* | -1.104 | 0.017902 |
| *SPAG16* | -1.103 | 0.005831 |
| *SATB1* | -1.103 | 0.003608 |
| *CENPM* | -1.103 | 0.012549 |
| *FAM13A* | -1.102 | 0.001988 |
| *TMEM205* | -1.102 | 0.002802 |
| *GTF2IRD2* | -1.100 | 0.007866 |
| *FANCE* | -1.100 | 0.010834 |
| *PTGDR2* | -1.100 | 0.005320 |
| *IQCN* | -1.099 | 0.032183 |
| *MAOA* | -1.097 | 0.004888 |
| *C1orf74* | -1.096 | 0.004851 |
| *IFIH1* | -1.095 | 0.026572 |
| *CXCL16* | -1.093 | 0.025421 |
| *LGALS3* | -1.092 | 0.021948 |
| *TEKT2* | -1.092 | 0.024026 |
| *RORA* | -1.090 | 0.013971 |
| *PLCL1* | -1.090 | 0.003464 |
| *RAB15* | -1.089 | 0.002777 |
| *NBEA* | -1.088 | 0.012756 |
| *FFAR2* | -1.088 | 0.004623 |
| *SH2D3A* | -1.088 | 0.003981 |
| *DOK7* | -1.087 | 0.014936 |
| *CYB561* | -1.086 | 0.005159 |
| *NOTCH2NLA* | -1.085 | 0.001840 |
| *TNIP1* | -1.085 | 0.002604 |
| *GUCA2A* | -1.085 | 0.018723 |
| *TMC8* | -1.085 | 0.019826 |
| *DHX58* | -1.083 | 0.010420 |
| *FAM95C* | -1.083 | 0.029638 |
| *PLEKHA7* | -1.083 | 0.033197 |
| *NIPAL3* | -1.082 | 0.038081 |
| *GAPVD1* | -1.082 | 0.002471 |
| *TJP3* | -1.081 | 0.001974 |
| *HLA-DOA* | -1.081 | 0.022698 |
| *DMTN* | -1.081 | 0.003152 |
| *ANKRD27* | -1.081 | 0.011375 |
| *KCNK17* | -1.080 | 0.003792 |
| *IRF7* | -1.080 | 0.039550 |
| *HLA-DMA* | -1.079 | 0.023634 |

| | | |
|---|---|---|
| *AKAP12* | -1.078 | 0.031537 |
| *C2CD4A* | -1.078 | 0.030742 |
| *TLCD2* | -1.077 | 0.017456 |
| *TUBA4A* | -1.074 | 0.019704 |
| *RASAL1* | -1.074 | 0.009397 |
| *SI* | -1.073 | 0.021445 |
| *PPP1R32* | -1.072 | 0.019605 |
| *ICOSLG* | -1.072 | 0.008046 |
| *GINS1* | -1.071 | 0.039429 |
| *ALCAM* | -1.071 | 0.017525 |
| *BCL6* | -1.071 | 0.026053 |
| *FOXA1* | -1.070 | 0.004061 |
| *TLR3* | -1.068 | 0.006402 |
| *TOM1L2* | -1.067 | 0.002647 |
| *HLA-B* | -1.066 | 0.022461 |
| *CARD6* | -1.066 | 0.045003 |
| *PIGA* | -1.066 | 0.005783 |
| *GZF1* | -1.064 | 0.002542 |
| *ELF4* | -1.063 | 0.002897 |
| *ANO1* | -1.062 | 0.010913 |
| *CAMKK1* | -1.062 | 0.014468 |
| *WDR90* | -1.062 | 0.037748 |
| *TP73* | -1.060 | 0.003336 |
| *CMTM8* | -1.059 | 0.027066 |
| *PSMF1* | -1.059 | 0.005417 |
| *BICDL1* | -1.059 | 0.046845 |
| *SOWAHB* | -1.059 | 0.012329 |
| *PPL* | -1.058 | 0.006826 |
| *RABGAP1L* | -1.058 | 0.005737 |
| *NYNRIN* | -1.057 | 0.010082 |
| *LYNX1* | -1.056 | 0.004470 |
| *IL17C* | -1.056 | 0.013912 |
| *ARRB2* | -1.055 | 0.002581 |
| *DLX3* | -1.053 | 0.007630 |
| *S100A16* | -1.053 | 0.022403 |
| *SEPTIN5* | -1.053 | 0.019551 |
| *IFI35* | -1.052 | 0.002066 |
| *H4C15* | -1.051 | 0.017775 |
| *PPM1L* | -1.051 | 0.044578 |
| *SPTB* | -1.051 | 0.002942 |
| *ZNF232* | -1.050 | 0.003169 |

| | | |
|---|---|---|
| *OSBPL10* | -1.050 | 0.002895 |
| *PARP10* | -1.048 | 0.007827 |
| *CNKSR1* | -1.048 | 0.035606 |
| *COMTD1* | -1.048 | 0.017919 |
| *RIMBP2* | -1.048 | 0.004973 |
| *MAPK3* | -1.047 | 0.002777 |
| *OLFML2A* | -1.046 | 0.016444 |
| *DTX2* | -1.044 | 0.005688 |
| *AKR1C2* | -1.044 | 0.007760 |
| *MAPK15* | -1.042 | 0.003379 |
| *PCDHGA3* | -1.042 | 0.002507 |
| *CASP7* | -1.042 | 0.008853 |
| *TAC3* | -1.042 | 0.025088 |
| *USP53* | -1.041 | 0.048687 |
| *KIF24* | -1.040 | 0.002417 |
| *NAALADL2* | -1.039 | 0.035220 |
| *ZNF98* | -1.039 | 0.003560 |
| *CBX6* | -1.038 | 0.033778 |
| *PPFIBP2* | -1.038 | 0.020609 |
| *PDE4A* | -1.037 | 0.004173 |
| *EMP1* | -1.037 | 0.013589 |
| *NPY5R* | -1.037 | 0.020334 |
| *DDX11* | -1.036 | 0.020608 |
| *VSTM5* | -1.035 | 0.004869 |
| *ZNF816-ZNF321P* | -1.035 | 0.006324 |
| *TRERF1* | -1.034 | 0.005835 |
| *ITIH4* | -1.034 | 0.013320 |
| *ESAM* | -1.033 | 0.026807 |
| *UBOX5* | -1.033 | 0.015142 |
| *LTB4R* | -1.032 | 0.044307 |
| *ERMAP* | -1.032 | 0.024775 |
| *FREM2* | -1.032 | 0.027608 |
| *RALGAPA1* | -1.031 | 0.025632 |
| *BAIAP2L1* | -1.031 | 0.023945 |
| *ACBD7* | -1.031 | 0.033720 |
| *MUTYH* | -1.031 | 0.028227 |
| *SYCP2* | -1.030 | 0.015361 |
| *PROCA1* | -1.030 | 0.030550 |
| *USP43* | -1.030 | 0.002844 |
| *CNTNAP3B* | -1.028 | 0.020015 |
| *AKNA* | -1.028 | 0.009663 |

| | | |
|---|---|---|
| *MCM8* | -1.028 | 0.005665 |
| *KRTCAP3* | -1.028 | 0.003431 |
| *BCL3* | -1.028 | 0.019358 |
| *NFIB* | -1.028 | 0.007351 |
| *SNPH* | -1.027 | 0.029018 |
| *CCDC57* | -1.027 | 0.029050 |
| *CNGA4* | -1.027 | 0.002497 |
| *NBPF10* | -1.026 | 0.039699 |
| *PAK6* | -1.025 | 0.003863 |
| *MT1F* | -1.023 | 0.007834 |
| *STXBP5L* | -1.023 | 0.027422 |
| *TBC1D30* | -1.020 | 0.029165 |
| *KRT15* | -1.018 | 0.021203 |
| *USP18* | -1.018 | 0.008775 |
| *ENO4* | -1.017 | 0.044005 |
| *IKZF1* | -1.016 | 0.036303 |
| *KLHL26* | -1.016 | 0.012073 |
| *DGKE* | -1.016 | 0.005883 |
| *PPP1R15A* | -1.015 | 0.003252 |
| *SRGAP1* | -1.015 | 0.027961 |
| *IZUMO4* | -1.015 | 0.004785 |
| *IAPP* | -1.015 | 0.018731 |
| *CDC25B* | -1.013 | 0.011099 |
| *ZNF337* | -1.013 | 0.048963 |
| *SPOPL* | -1.012 | 0.007724 |
| *CDKL5* | -1.011 | 0.043983 |
| *SCGB3A1* | -1.011 | 0.042434 |
| *KRT19* | -1.009 | 0.003439 |
| *NUP210* | -1.009 | 0.035957 |
| *SEC23B* | -1.008 | 0.007018 |
| *LLGL2* | -1.008 | 0.009128 |
| *PLP2* | -1.008 | 0.011470 |
| *CCDC159* | -1.007 | 0.027061 |
| *TICAM1* | -1.007 | 0.047406 |
| *FBXO32* | -1.006 | 0.002379 |
| *NBPF19* | -1.006 | 0.023508 |
| *PRXL2B* | -1.005 | 0.010893 |
| *DTD1* | -1.005 | 0.048617 |
| *ADRA2B* | -1.003 | 0.003941 |
| *CTSS* | -1.003 | 0.008804 |
| *SPATA12* | -1.002 | 0.018480 |

| | | |
|---|---|---|
| *PLCB4* | -1.002 | 0.004344 |
| *PCDHGA7* | -1.002 | 0.003411 |
| *DUSP7* | -1.001 | 0.044091 |
| *RGS11* | -1.001 | 0.019069 |
| *ZNF880* | -1.001 | 0.003628 |

**Supplementary Table 5.** Top upregulated DEmiRs in *FOXA2*$^{-/-}$ islets compared with WT-islets (Log2 FC > 1, *P* < 0.05).

| **miRNA ID** | **Log2 FC** | ***P*-value** |
|---|---|---|
| hsa-miR-199a-5p | 6.29761 | 0.000148 |
| hsa-miR-214-3p | 5.76148 | 0.000232 |
| hsa-miR-10a-5p | 4.66772 | 0.000003 |
| hsa-miR-214-5p | 4.58377 | 0.000730 |
| hsa-miR-199a-3p | 4.28411 | 0.001016 |
| hsa-miR-204-5p | 3.81157 | 0.012544 |
| hsa-miR-490-3p | 3.74824 | 0.000203 |
| hsa-miR-133a-5p | 3.14915 | 0.000027 |
| hsa-miR-122-5p | 3.09211 | 0.004089 |
| hsa-miR-133a-3p | 2.95520 | 0.000029 |
| hsa-miR-1-3p | 2.92150 | 0.000014 |
| hsa-miR-383-5p | 2.66077 | 0.002652 |
| hsa-miR-10b-5p | 2.64874 | 0.000668 |
| hsa-miR-618 | 2.57778 | 0.000094 |
| hsa-miR-196b-5p | 2.36430 | 0.000054 |
| hsa-miR-133b | 2.32345 | 0.000464 |
| hsa-miR-125b-1-3p | 2.29296 | 0.003248 |
| hsa-miR-218-5p | 2.16414 | 0.000457 |
| hsa-miR-145-5p | 2.10212 | 0.009372 |
| hsa-miR-145-3p | 2.06116 | 0.001509 |
| hsa-miR-365a-5p | 2.02153 | 0.000383 |
| hsa-miR-206 | 1.88778 | 0.005863 |
| hsa-miR-372-3p | 1.88276 | 0.000215 |
| hsa-miR-100-5p | 1.87694 | 0.021955 |
| hsa-miR-504-5p | 1.82716 | 0.000288 |
| hsa-miR-2682-5p | 1.82199 | 0.000314 |
| hsa-miR-373-3p | 1.76612 | 0.000992 |
| hsa-miR-143-3p | 1.69133 | 0.023539 |
| hsa-miR-371a-5p | 1.63440 | 0.002390 |

| | | |
|---|---|---|
| hsa-miR-137-3p | 1.63327 | 0.005341 |
| hsa-miR-371a-3p | 1.62716 | 0.000863 |
| hsa-miR-152-5p | 1.61188 | 0.003248 |
| hsa-miR-10a-3p | 1.54962 | 0.001363 |
| hsa-miR-155-5p | 1.46370 | 0.001120 |
| hsa-miR-105-5p | 1.43233 | 0.021987 |
| hsa-miR-129-5p | 1.41903 | 0.007961 |
| hsa-miR-1269b | 1.41806 | 0.003371 |
| hsa-miR-1263 | 1.38661 | 0.006575 |
| hsa-miR-3085-3p | 1.38094 | 0.002291 |
| hsa-miR-708-5p | 1.38026 | 0.001880 |
| hsa-miR-106a-3p | 1.36400 | 0.035987 |
| hsa-miR-585-3p | 1.34695 | 0.002726 |
| hsa-miR-9983-3p | 1.30991 | 0.012296 |
| hsa-miR-193b-3p | 1.28944 | 0.017239 |
| hsa-miR-3117-3p | 1.28800 | 0.018137 |
| hsa-miR-6815-5p | 1.26580 | 0.007245 |
| hsa-miR-30b-3p | 1.23908 | 0.003385 |
| hsa-miR-1251-3p | 1.20758 | 0.021147 |
| hsa-miR-548w | 1.19917 | 0.004609 |
| hsa-miR-760 | 1.16573 | 0.012776 |
| hsa-miR-193b-5p | 1.14683 | 0.003133 |
| hsa-miR-10401-3p | 1.13964 | 0.031666 |
| hsa-miR-6843-3p | 1.13868 | 0.031624 |
| hsa-miR-548ah-3p | 1.13642 | 0.043918 |
| hsa-miR-365a-3p | 1.12864 | 0.006508 |
| hsa-miR-129-2-3p | 1.11738 | 0.017599 |
| hsa-miR-514a-3p | 1.09436 | 0.016005 |
| hsa-miR-489-3p | 1.07911 | 0.003577 |
| hsa-miR-887-5p | 1.05318 | 0.005964 |
| hsa-miR-34b-3p | 1.04424 | 0.022495 |
| hsa-miR-887-3p | 1.03743 | 0.027094 |

**Supplementary Table 6.** Top downregulated DEmiRs in *FOXA2*$^{-/-}$ islets compared with WT-islets (Log2 FC < −1, *P* < 0.05).

| **miRNA ID** | **Log2 FC** | ***P*-value** |
|---|---|---|
| hsa-let-7d-5p | -4.18566 | 0.00017 |
| hsa-miR-98-5p | -3.79267 | 0.00057 |
| hsa-let-7d-3p | -3.69696 | 0.00025 |
| hsa-let-7i-5p | -3.55578 | 0.00099 |
| hsa-let-7g-5p | -3.48416 | 0.00021 |
| hsa-miR-146a-5p | -3.46109 | 0.00001 |
| hsa-let-7c-5p | -3.35395 | 0.00036 |
| hsa-let-7b-5p | -3.33790 | 0.00055 |
| hsa-miR-3059-5p | -3.29272 | 0.00015 |
| hsa-miR-551b-5p | -3.07552 | 0.00003 |
| hsa-let-7f-5p | -3.00945 | 0.00043 |
| hsa-miR-203b-3p | -2.87888 | 0.00002 |
| hsa-miR-551b-3p | -2.86735 | 0.00015 |
| hsa-miR-127-3p | -2.78930 | 0.00172 |
| hsa-let-7f-1-3p | -2.66048 | 0.00217 |
| hsa-let-7f-2-3p | -2.50539 | 0.00827 |
| hsa-miR-136-3p | -2.47541 | 0.00301 |
| hsa-miR-433-3p | -2.41518 | 0.00417 |
| hsa-let-7a-5p | -2.39032 | 0.00106 |
| hsa-miR-934 | -2.38368 | 0.00027 |
| hsa-let-7a-3p | -2.37988 | 0.00250 |
| hsa-let-7c-3p | -2.34103 | 0.00145 |
| hsa-miR-493-5p | -2.29658 | 0.00725 |
| hsa-miR-370-3p | -2.28731 | 0.00262 |
| hsa-miR-127-5p | -2.25616 | 0.00232 |
| hsa-miR-493-3p | -2.15845 | 0.00375 |
| hsa-miR-411-5p | -2.13715 | 0.00126 |
| hsa-miR-409-5p | -2.12795 | 0.00445 |
| hsa-miR-487b-3p | -2.11730 | 0.00365 |
| hsa-miR-494-3p | -2.11026 | 0.00703 |
| hsa-miR-382-3p | -2.11023 | 0.00981 |
| hsa-miR-412-5p | -2.09312 | 0.00037 |
| hsa-miR-382-5p | -2.08008 | 0.00453 |
| hsa-miR-381-3p | -2.06745 | 0.00549 |
| hsa-miR-432-5p | -2.04222 | 0.00370 |
| hsa-let-7b-3p | -2.02456 | 0.00311 |
| hsa-miR-873-3p | -2.01125 | 0.00049 |
| hsa-miR-409-3p | -2.00956 | 0.01033 |

| | | |
|---|---|---|
| hsa-miR-134-5p | -1.98735 | 0.00342 |
| hsa-miR-654-5p | -1.93030 | 0.00616 |
| hsa-miR-99a-3p | -1.89894 | 0.00349 |
| hsa-miR-323a-3p | -1.87578 | 0.00520 |
| hsa-miR-873-5p | -1.85647 | 0.00036 |
| hsa-miR-495-3p | -1.84195 | 0.00700 |
| hsa-miR-655-3p | -1.81740 | 0.00128 |
| hsa-miR-487a-5p | -1.79663 | 0.00149 |
| hsa-miR-543 | -1.76665 | 0.00359 |
| hsa-miR-200a-3p | -1.76574 | 0.00061 |
| hsa-miR-1224-3p | -1.75385 | 0.00473 |
| hsa-miR-375-3p | -1.70244 | 0.01426 |
| hsa-miR-98-3p | -1.69519 | 0.00722 |
| hsa-miR-539-3p | -1.69198 | 0.00620 |
| hsa-miR-485-3p | -1.67730 | 0.00407 |
| hsa-miR-642a-5p | -1.58584 | 0.00374 |
| hsa-miR-625-3p | -1.56552 | 0.00053 |
| hsa-miR-431-3p | -1.56424 | 0.00120 |
| hsa-let-7i-3p | -1.56051 | 0.01407 |
| hsa-miR-429 | -1.54112 | 0.00101 |
| hsa-miR-141-3p | -1.52582 | 0.00084 |
| hsa-miR-203a-3p | -1.51655 | 0.03505 |
| hsa-miR-758-3p | -1.48987 | 0.01376 |
| hsa-let-7e-5p | -1.46834 | 0.00342 |
| hsa-miR-4510 | -1.46180 | 0.01151 |
| hsa-miR-876-3p | -1.43503 | 0.01689 |
| hsa-miR-668-3p | -1.42099 | 0.00443 |
| hsa-miR-182-3p | -1.38475 | 0.00920 |
| hsa-miR-200b-3p | -1.38147 | 0.00193 |
| hsa-miR-889-3p | -1.37138 | 0.01140 |
| hsa-miR-485-5p | -1.37054 | 0.04171 |
| hsa-miR-125b-2-3p | -1.34979 | 0.00590 |
| hsa-miR-329-3p | -1.34386 | 0.03001 |
| hsa-miR-200b-5p | -1.32878 | 0.00189 |
| hsa-miR-1185-1-3p | -1.32778 | 0.02417 |
| hsa-miR-3934-5p | -1.29827 | 0.00676 |
| hsa-miR-891a-5p | -1.26090 | 0.00617 |
| hsa-miR-642a-3p | -1.23877 | 0.00540 |
| hsa-miR-31-5p | -1.23331 | 0.01249 |
| hsa-miR-539-5p | -1.22424 | 0.01525 |
| hsa-miR-340-5p | -1.17002 | 0.01578 |

| | | |
|---|---|---|
| hsa-miR-542-3p | -1.15544 | 0.00960 |
| hsa-miR-1250-5p | -1.15045 | 0.00437 |
| hsa-miR-146b-3p | -1.13802 | 0.00789 |
| hsa-miR-154-3p | -1.12851 | 0.02181 |
| hsa-miR-21-3p | -1.12340 | 0.00495 |
| hsa-miR-200a-5p | -1.11795 | 0.00462 |
| hsa-miR-31-3p | -1.11183 | 0.00729 |
| hsa-miR-337-3p | -1.10772 | 0.00680 |
| hsa-miR-296-5p | -1.10493 | 0.00326 |
| hsa-miR-99a-5p | -1.09727 | 0.01150 |
| hsa-miR-32-5p | -1.08862 | 0.03895 |
| hsa-miR-301a-3p | -1.07225 | 0.04241 |
| hsa-miR-450a-5p | -1.07125 | 0.00751 |
| hsa-miR-376a-3p | -1.06003 | 0.02819 |
| hsa-miR-4517 | -1.05251 | 0.01394 |
| hsa-miR-181c-5p | -1.04678 | 0.00587 |
| hsa-miR-3177-3p | -1.04512 | 0.02936 |
| hsa-miR-4728-5p | -1.01159 | 0.01216 |
| hsa-miR-146b-5p | -1.01154 | 0.01420 |
| hsa-miR-892b | -1.00092 | 0.00963 |

**Supplementary Table 7.** Top downregulated DEmiRs (Log2 FC < −1, $P$ < 0.05) and their predicted upregulated target DEGs (Log2 FC > 2, $P$ < 0.05) associated with nervous system development in *FOXA2*$^{-/-}$ islets compared with WT-islets.

| Upregulated miRNA | Log2 FC | *P*-value | Predicted target gene | Log2 FC | *P*-value |
|---|---|---|---|---|---|
| hsa-miR-429 | -1.541 | 0.001015 | *RELN* | 5.035 | 0.00003 |
| hsa-miR-323a-3p | -1.876 | 0.005197 | *GABRP* | 4.446 | 0.00094 |
| hsa-miR-337-3p | -1.108 | 0.006802 | | | |
| hsa-miR-146a-5p | -3.461 | 0.000007 | *WNT2B* | 4.067 | 0.00036 |
| hsa-miR-934 | -2.384 | 0.000274 | | | |
| hsa-miR-1224-3p | -1.754 | 0.004729 | | | |
| hsa-miR-4510 | -1.462 | 0.01151 | | | |
| hsa-miR-485-5p | -1.371 | 0.041705 | | | |
| hsa-miR-301a-3p | -1.072 | 0.042411 | | | |
| hsa-miR-181c-5p | -1.047 | 0.005868 | | | |
| hsa-miR-4728-5p | -1.012 | 0.012164 | | | |
| hsa-let-7d-5p | -4.186 | 0.000168 | *LIN28A* | 4.004 | 0.00005 |
| hsa-miR-4510 | -1.462 | 0.01151 | | | |

| | | | | | |
|---|---|---|---|---|---|
| hsa-miR-329-3p<br>hsa-miR-296-5p<br>hsa-miR-181c-5p | -1.344<br>-1.105<br>-1.047 | 0.030007<br>0.00326<br>0.005868 | | | |
| hsa-miR-4728-5p | -1.012 | 0.012164 | *ADGRA2* | 3.913 | 0.00009 |
| hsa-miR-495-3p<br>hsa-miR-655-3p<br>hsa-miR-200a-3p<br>hsa-miR-542-3p<br>hsa-miR-32-5p<br>hsa-miR-450a-5p | -1.842<br>-1.817<br>-1.766<br>-1.155<br>-1.089<br>-1.071 | 0.007002<br>0.001278<br>0.000607<br>0.009603<br>0.038947<br>0.00751 | *ALKAL2* | 3.619 | 0.00004 |
| hsa-miR-655-3p | -1.817 | 0.001278 | *BMP4* | 3.558 | 0.00020 |
| hsa-miR-200a-3p<br>hsa-miR-539-3p | -1.766<br>-1.692 | 0.000607<br>0.006197 | *OLIG3* | 3.439 | 0.00091 |
| hsa-miR-375-3p<br>hsa-miR-668-3p<br>hsa-miR-329-3p<br>hsa-miR-31-5p<br>hsa-miR-154-3p<br>hsa-miR-301a-3p<br>hsa-miR-450a-5p<br>hsa-miR-4728-5p | -1.702<br>-1.421<br>-1.344<br>-1.233<br>-1.129<br>-1.072<br>-1.071<br>-1.012 | 0.014259<br>0.004435<br>0.030007<br>0.012494<br>0.021808<br>0.042411<br>0.00751<br>0.012164 | *CXCL12* | 3.428 | 0.00310 |
| hsa-miR-654-5p<br>hsa-miR-876-3p<br>hsa-miR-4728-5p<br>hsa-miR-892b | -1.93<br>-1.435<br>-1.012<br>-1.001 | 0.006161<br>0.016891<br>0.012164<br>0.00963 | *WNT6* | 3.256 | 0.01048 |
| hsa-miR-296-5p<br>hsa-miR-4728-5p | -1.105<br>-1.012 | 0.003260<br>0.012164 | *SEMA6B* | 3.142 | 0.00040 |
| hsa-let-7d-5p<br>hsa-let-7c-3p<br>hsa-miR-493-5p<br>hsa-miR-654-5p<br>hsa-miR-1224-3p<br>hsa-miR-4728-5p | -4.186<br>-2.341<br>-2.297<br>-1.93<br>-1.754<br>-1.012 | 0.000168<br>0.001446<br>0.007254<br>0.006161<br>0.004729<br>0.012164 | *GDF6* | 3.087 | 0.00059 |
| hsa-miR-411-5p<br>hsa-miR-625-3p | -2.137<br>-1.566 | 0.001262<br>0.000532 | *PTN* | 3.071 | 0.00068 |
| hsa-miR-655-3p<br>hsa-miR-539-3p<br>hsa-miR-668-3p<br>hsa-miR-31-5p<br>hsa-miR-340-5p | -1.817<br>-1.692<br>-1.421<br>-1.233<br>-1.17 | 0.001278<br>0.006197<br>0.004435<br>0.012494<br>0.015782 | *EDNRB* | 3.07 | 0.00014 |

| | | | | | |
|---|---|---|---|---|---|
| hsa-let-7c-3p<br>hsa-miR-432-5p<br>hsa-miR-654-5p<br>hsa-miR-625-3p<br>hsa-miR-301a-3p | -2.341<br>-2.042<br>-1.93<br>-1.566<br>-1.072 | 0.001446<br>0.0037<br>0.006161<br>0.000532<br>0.042411 | *HAPLN1* | 3.03 | 0.00047 |
| hsa-miR-433-3p<br>hsa-miR-375-3p<br>hsa-miR-21-3p<br>hsa-miR-32-5p<br>hsa-miR-301a-3p<br>hsa-miR-4728-5p | -2.415<br>-1.702<br>-1.123<br>-1.089<br>-1.072<br>-1.012 | 0.004174<br>0.014259<br>0.004947<br>0.038947<br>0.042411<br>0.012164 | *GAP43* | 3.022 | 0.02036 |
| hsa-miR-200a-3p<br>hsa-miR-539-3p<br>hsa-miR-203a-3p<br>hsa-miR-182-3p<br>hsa-miR-200b-5p<br>hsa-miR-539-5p | -1.766<br>-1.692<br>-1.517<br>-1.385<br>-1.329<br>-1.224 | 0.000607<br>0.006197<br>0.035053<br>0.009204<br>0.00189<br>0.015253 | *GABRA2* | 2.927 | 0.00079 |
| hsa-miR-4728-5p | -1.012 | 0.012164 | *TFAP2B* | 2.867 | 0.00236 |
| hsa-miR-432-5p | -2.042 | 0.003700 | *NTRK2* | 2.861 | 0.00761 |
| hsa-miR-642a-5p | -1.586 | 0.003740 | *CTNNA2* | 2.852 | 0.00030 |
| hsa-let-7c-3p<br>hsa-miR-340-5p<br>hsa-miR-337-3p | -2.341<br>-1.17<br>-1.108 | 0.001446<br>0.015782<br>0.006802 | *HOXB2* | 2.841 | 0.00862 |
| hsa-let-7c-3p<br>hsa-miR-493-5p<br>hsa-miR-99a-3p<br>hsa-miR-32-5p | -2.341<br>-2.297<br>-1.899<br>-1.089 | 0.001446<br>0.007254<br>0.003487<br>0.038947 | *GATA2* | 2.807 | 0.00119 |
| hsa-miR-32-5p | -1.089 | 0.038947 | *HAND2* | 2.789 | 0.00825 |
| hsa-let-7d-5p<br>hsa-miR-495-3p<br>hsa-miR-655-3p<br>hsa-miR-31-5p<br>hsa-miR-296-5p | -4.186<br>-1.842<br>-1.817<br>-1.233<br>-1.105 | 0.000168<br>0.007002<br>0.001278<br>0.012494<br>0.00326 | *FZD4* | 2.773 | 0.00006 |
| hsa-miR-99a-3p<br>hsa-miR-892b | -1.899<br>-1.001 | 0.003487<br>0.009630 | *SFRP1* | 2.668 | 0.00015 |
| hsa-miR-494-3p<br>hsa-miR-3934-5p<br>hsa-miR-31-3p | -2.11<br>-1.298<br>-1.112 | 0.007032<br>0.006761<br>0.007288 | *PCP4* | 2.651 | 0.00029 |
| hsa-miR-382-3p | -2.11 | 0.009809 | *DDR2* | 2.639 | 0.00062 |
| hsa-miR-146a-5p | -3.461 | 0.000007 | *ROR1* | 2.588 | 0.00029 |

| | | | | | |
|---|---|---|---|---|---|
| hsa-miR-127-5p<br>hsa-miR-873-3p<br>hsa-miR-892b | -2.256<br>-2.011<br>-1.001 | 0.002322<br>0.000493<br>0.00963 | | | |
| hsa-miR-429<br>hsa-miR-4510<br>hsa-miR-296-5p<br>hsa-miR-4728-5p | -1.541<br>-1.462<br>-1.105<br>-1.012 | 0.001015<br>0.01151<br>0.00326<br>0.012164 | *CNTFR* | 2.555 | 0.01953 |
| hsa-miR-134-5p<br>hsa-miR-542-3p | -1.987<br>-1.155 | 0.003423<br>0.009603 | *MMP24* | 2.409 | 0.00079 |
| hsa-let-7f-1-3p | -2.66 | 0.002172 | *FGF2* | 2.402 | 0.00015 |
| hsa-miR-493-5p<br>hsa-miR-429<br>hsa-miR-4510<br>hsa-miR-485-5p<br>hsa-miR-4728-5p | -2.297<br>-1.541<br>-1.462<br>-1.371<br>-1.012 | 0.007254<br>0.001015<br>0.01151<br>0.041705<br>0.012164 | *PMP22* | 2.283 | 0.00240 |
| hsa-let-7f-1-3p<br>hsa-let-7f-2-3p<br>hsa-miR-539-3p<br>hsa-miR-429 | -2.66<br>-2.505<br>-1.692<br>-1.541 | 0.002172<br>0.008274<br>0.006197<br>0.001015 | *EDNRA* | 2.22 | 0.00114 |
| hsa-miR-4510<br>hsa-miR-125b-2-3p<br>hsa-miR-200b-5p<br>hsa-miR-642a-3p<br>hsa-miR-21-3p<br>hsa-miR-337-3p<br>hsa-miR-4728-5p | -1.462<br>-1.35<br>-1.329<br>-1.239<br>-1.123<br>-1.108<br>-1.012 | 0.01151<br>0.005899<br>0.00189<br>0.005401<br>0.004947<br>0.006802<br>0.012164 | | | |
| hsa-let-7f-1-3p<br>hsa-let-7f-2-3p<br>hsa-miR-4728-5p | -2.66<br>-2.505<br>-1.012 | 0.002172<br>0.008274<br>0.012164 | *NR2F2* | 2.199 | 0.00065 |
| hsa-miR-934<br>hsa-miR-99a-3p<br>hsa-miR-200a-3p<br>hsa-miR-431-3p | -2.384<br>-1.899<br>-1.766<br>-1.564 | 0.000274<br>0.003487<br>0.000607<br>0.001198 | *NTNG1* | 2.192 | 0.00060 |
| hsa-let-7d-5p<br>hsa-miR-4510<br>hsa-miR-296-5p<br>hsa-miR-3177-3p<br>hsa-miR-892b | -4.186<br>-1.462<br>-1.105<br>-1.045<br>-1.001 | 0.000168<br>0.011510<br>0.003260<br>0.029363<br>0.009630 | *WNT9B* | 2.131 | 0.00133 |
| hsa-miR-429<br>hsa-miR-4510 | -1.541<br>-1.462 | 0.001015<br>0.011510 | *SLIT2* | 2.123 | 0.00090 |

| | | | | | |
|---|---|---|---|---|---|
| hsa-miR-182-3p | -1.385 | 0.009204 | | | |
| hsa-let-7c-3p | -2.341 | 0.001446 | *FGF19* | 2.086 | 0.02645 |
| hsa-miR-654-5p | -1.93 | 0.006161 | | | |
| hsa-miR-4510 | -1.462 | 0.011510 | | | |
| hsa-miR-200b-5p | -1.329 | 0.001890 | *BEX1* | 2.061 | 0.00381 |
| hsa-miR-542-3p | -1.155 | 0.009603 | | | |
| hsa-miR-4510 | -1.462 | 0.011510 | *PCDHGB7* | 2.061 | 0.00256 |
| hsa-miR-4728-5p | -1.012 | 0.012164 | | | |
| hsa-miR-892b | -1.001 | 0.009630 | | | |

**Supplementary Table 8.** Top downregulated DEmiRs (Log2 FC < -1, *P* < 0.05) and their predicted upregulated target DEGs (Log2 FC > 1, *P* < 0.05) associated with lipid metabolism in *FOXA2*$^{-/-}$ islets compared with WT-islets.

| Upregulated miRNA | Log2 FC | *P*-value | Predicted target gene | Log2 FC | *P*-value |
|---|---|---|---|---|---|
| hsa-miR-429 | -1.541 | 0.001015 | *APOC3* | 3.897 | 0.006166 |
| hsa-miR-4728-5p | -1.012 | 0.012164 | | | |
| hsa-miR-654-5p | -1.93 | 0.006161 | *PDGFRB* | 3.035 | 0.000385 |
| hsa-miR-892b | -1.001 | 0.009630 | | | |
| hsa-miR-146a-5p | -3.461 | 0.000007 | *PDGFRA* | 2.958 | 0.000049 |
| hsa-miR-301a-3p | -1.072 | 0.042411 | | | |
| hsa-miR-487a-5p | -1.797 | 0.001493 | *PLA2G2A* | 2.781 | 0.012291 |
| hsa-miR-4510 | -1.462 | 0.011510 | | | |
| hsa-miR-4728-5p | -1.012 | 0.012164 | | | |
| hsa-miR-4510 | -1.462 | 0.011510 | *APOC2* | 2.535 | 0.008866 |
| hsa-miR-4728-5p | -1.012 | 0.012164 | | | |
| hsa-let-7f-1-3p | -2.66 | 0.002172 | *FGF2* | 2.402 | 0.000148 |
| hsa-miR-31-5p | -1.233 | 0.012494 | *SERPINA5* | 2.376 | 0.002514 |
| hsa-miR-376a-3p | -1.06 | 0.028186 | | | |
| hsa-let-7d-5p | -4.186 | 0.00017 | *HMOX1* | 2.34 | 0.002408 |
| hsa-miR-494-3p | -2.11 | 0.00703 | | | |
| hsa-miR-873-5p | -1.856 | 0.00036 | | | |
| hsa-miR-642a-5p | -1.586 | 0.00374 | | | |
| hsa-miR-485-5p | -1.371 | 0.04171 | | | |
| hsa-miR-539-3p | -1.692 | 0.006197 | *MTTP* | 2.339 | 0.011627 |
| hsa-miR-296-5p | -1.105 | 0.003260 | | | |
| hsa-miR-134-5p | -1.987 | 0.003423 | *AKR1D1* | 2.047 | 0.021353 |
| hsa-miR-873-5p | -1.856 | 0.000360 | | | |
| hsa-miR-146a-5p | -3.461 | 0.000007 | *SOAT2* | 1.95 | 0.000241 |
| hsa-miR-301a-3p | -1.072 | 0.042411 | *FABP1* | 1.817 | 0.009621 |
| hsa-miR-4517 | -1.053 | 0.013944 | | | |
| hsa-miR-432-5p | -2.042 | 0.0037 | *NR5A2* | 1.754 | 0.005532 |
| hsa-miR-134-5p | -1.987 | 0.00342 | | | |
| hsa-miR-655-3p | -1.817 | 0.00128 | | | |

| | | | | | |
|---|---|---|---|---|---|
| hsa-miR-429 | -1.541 | 0.00102 | | | |
| hsa-miR-4510 | -1.462 | 0.01151 | | | |
| hsa-miR-31-5p | -1.233 | 0.01249 | | | |
| hsa-miR-450a-5p | -1.071 | 0.00751 | | | |
| hsa-miR-4728-5p | -1.012 | 0.01216 | | | |
| hsa-miR-376a-3p | -1.06 | 0.02819 | | | |
| hsa-miR-370-3p | -2.287 | 0.002624 | *PLA2G12B* | 1.577 | 0.020816 |
| hsa-miR-494-3p | -2.11 | 0.007032 | | | |
| hsa-miR-668-3p | -1.421 | 0.009630 | | | |
| hsa-miR-892b | -1.001 | 0.009630 | | | |
| hsa-miR-4510 | -1.462 | 0.011510 | *CIDEC* | 1.559 | 0.005503 |
| hsa-miR-551b-3p | -2.867 | 0.000153 | *CCDC3* | 1.511 | 0.016466 |
| hsa-miR-1224-3p | -1.754 | 0.004729 | *CYP27A1* | 1.483 | 0.002474 |
| hsa-miR-4510 | -1.462 | 0.011510 | | | |
| hsa-miR-296-5p | -1.105 | 0.003260 | *SCARB1* | 1.367 | 0.007276 |
| hsa-miR-296-5p | -1.105 | 0.003260 | *FGFR4* | 1.255 | 0.048591 |
| hsa-miR-3177-3p | -1.045 | 0.029363 | | | |
| hsa-miR-4728-5p | -1.012 | 0.012164 | | | |
| hsa-miR-181c-5p | -1.047 | 0.005868 | *FLT1* | 1.223 | 0.041628 |
| hsa-miR-892b | -1.001 | 0.009630 | | | |
| hsa-miR-127-5p | -2.256 | 0.00232 | *RGN* | 1.204 | 0.011920 |
| hsa-miR-382-5p | -2.08 | 0.00453 | | | |
| hsa-miR-203a-3p | -1.517 | 0.03505 | | | |
| hsa-miR-485-5p | -1.371 | 0.04171 | | | |
| hsa-miR-1224-3p | -1.754 | 0.004729 | *NR0B2* | 1.199 | 0.004540 |
| hsa-miR-539-3p | -1.692 | 0.006197 | | | |
| hsa-miR-485-5p | -1.371 | 0.041705 | | | |
| hsa-miR-301a-3p | -1.072 | 0.042411 | *LRP2* | 1.186 | 0.028674 |
| hsa-miR-4517 | -1.053 | 0.013944 | *NR1H4* | 1.101 | 0.003411 |
| hsa-miR-495-3p | -1.842 | 0.007002 | *SULT2A1* | 1.095 | 0.048243 |
| hsa-miR-4510 | -1.462 | 0.011510 | *FGFR1* | 1.008 | 0.014626 |
| hsa-miR-296-5p | -1.105 | 0.003260 | | | |
| hsa-miR-4728-5p | -1.012 | 0.012164 | | | |
| hsa-miR-493-5p | -2.297 | 0.007254 | *KIT* | 1.003 | 0.044687 |
| hsa-miR-494-3p | -2.11 | 0.007032 | | | |
| hsa-miR-539-3p | -1.692 | 0.006197 | | | |
